# Templating and confining calcium phosphate mineralization within designed protein assemblies

**DOI:** 10.64898/2026.01.14.699524

**Authors:** Le Tracy Yu, Xinqi Li, Harley Pyles, Anjali P. Patni, Andrew J. Borst, Neville P. Bethel, Paul S. Kwon, Connor Weidle, Ryan D. Kibler, Kenneth D. Carr, Yulai Liu, Dongsheng Li, Stanislav Moroz, Hannele Ruohola-Baker, Shuai Zhang, James De Yoreo, David Baker

**Author notes:** These authors contributed equally to this work (Xinqi Li, Harley Pyles).

## Abstract

Bone formation involves the deposition of ordered hydroxyapatite (HAp) on collagen fibrils, but the underlying molecular mechanisms remain largely unresolved, limiting the design of protein-apatite hybrid materials. Here we show that computationally designed de novo proteins can template and confine HAp mineralization. We design C_3_-symmetric oligomers with inner surfaces chemically complementary to the HAp {010} facet, and assemble these through additional interfaces into D_3_ oligomers, one-dimensional nanotubes, and two-dimensional arrays. Electron microscopy revealed templated mineralization and confinement at each hierarchical level, with mineral shape guided by the underlying protein architecture. Conversion of an initially formed amorphous phase to HAp is driven by the designed protein-mineral interface. Our results establish a framework for programmable protein-guided mineralization, providing a foundation for next-generation hybrid materials.

## Main Text

Bone formation requires biomineralization of calcium phosphate within a collagen matrix(*1–3*), a process regulated by non-collagenous proteins that promote ordered HAp crystal formation on collagen fibrils.(*4*) However, the molecular mechanisms underlying this process remain largely unresolved due to the structural complexity of the extracellular matrix. Biomimetic mineralization systems have been developed to investigate the physicochemical principles governing mineral nucleation and organization. Studies of peptide(*5*, *6*), protein(*7–12*), DNA(*13*, *14*) and polymer(*15*, *16*) model systems have shown that charged residues play critical roles in directing mineralization. While these studies have provided insight into the role of charge, and simple flat negatively charged interfaces have been shown to promote calcium carbonate nucleation (*17*), confined mineral growth within specific locations of a biomolecular assembly has not yet been achieved.

We reasoned that the control provided by computational protein design(*18*), which enables the construction of well-defined protein architectures with programmable surface chemistry, could provide a solution to the challenge of templating and confining mineral growth. Recent advances in deep learning based protein design methods such as RFdiffusion(*19–21*), ProteinMPNN and related methods (*22*, *23*) now allow the creation of proteins with precisely defined shapes and side-chain chemistries (*24–29*). We set out to these methods to design protein subunits with interfaces chemically and geometrically complementary to HAp lattice planes, and to explore the capability of these proteins to direct and confine HAp mineralization within a range of hierarchical assemblies.

### Design strategy

Hydroxyapatite – Ca_10_(PO_4_)_6_(OH)_2_ – is the most stable phase of calcium phosphate in physiological conditions and has a hexagonal crystal structure (space group P6_3_/m)(*30*). We built a model of a HAp crystal with a diameter of 3.8 nm as the design target. The model presents {001} facets along the c-axis and terminates at {010} facets along the a and b axes (Fig. 1A), which exposes calcium ions(*31*) to be templated using carboxylate sidechains (Fig. 1A). This HAp crystal has C_3_ rotational symmetry, and we set out to design trimeric protein oligomers with matching C_3_ symmetry with inner surfaces optimized to nucleate the target crystal. We reasoned that such HAp templating trimers could serve as building blocks for hierarchical architectures, enabling assembly into D_3_ oligomers, one-dimensional nanotubes and two-dimensional arrays (Fig. 1B). Each assembly would present spatially arranged aspartate and glutamate residues to promote HAp formation at specific locations, giving rise to ordered protein-HAp composite materials.

**Fig. 1.**
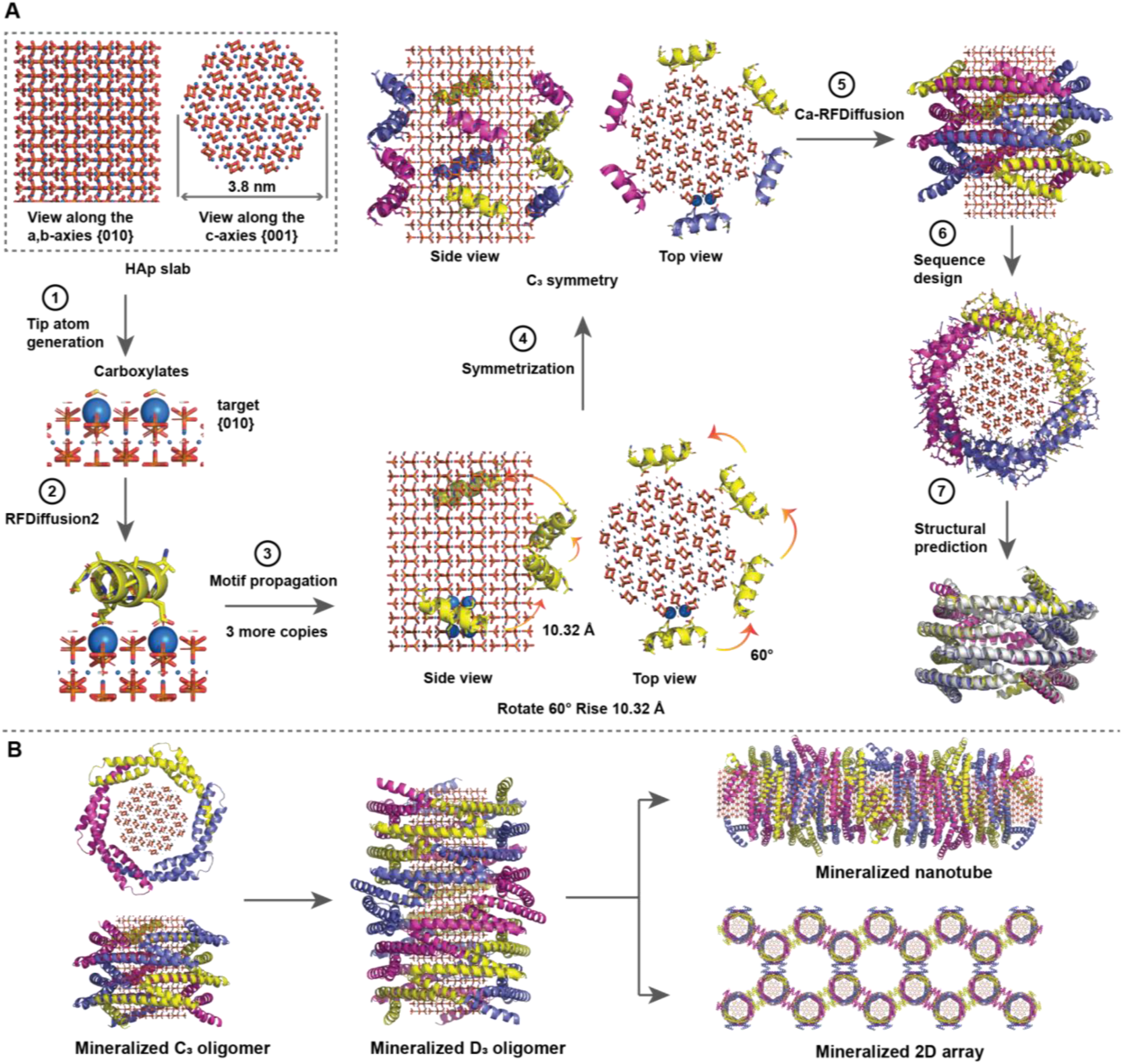
Design strategy. (**A**) Computational design work flow for generating C_3_ oligomers to stabilize HAp precursor and promote nucleation. The design starts with a hexagonal model of HAp that terminates at {010} facets at a and b directions. Carboxylates (yellow residues) were placed to interact with the calcium ions (blue balls) from HAp. The yellow alpha helix was generated with RFD2 and copies were propagated with a helical rise and twist to “wrap” the HAp model. The four helices together form an asymmetric unit which was transformed into a C_3_ oligomer (the two additional chains are in magenta and purple). Gaps between the helices were filled using RFdiffusion(*32*). In step 7, the AF2 prediction is in gray and the design model is colored. (**B**) Hierarchical design plan. The first step of the hierarchy is the mineralized C_3_ oligomer; the feature in the middle of the ring represents HAp. C_3_ oligomers were then designed into D_3_ oligomers which were then designed into one-dimensional protein nanotubes or two-dimensional protein arrays.

Carboxylate residues were placed at coordinates matching the phosphate groups surrounding the outermost calcium ions of the target surfaces, mimicking their calcium-coordinating geometry (Fig. 1A, step 1), and we used RFdiffusion2(*21*) (RFD2) to generate protein backbone segments hosting those carboxylate positions in aspartate or glutamate side chains.(Fig. 1A, step 2). These segments were then propagated to equivalent positions; in total three additional copies were generated, each sequentially rotated by 60° and translated by 10.32 Å along the c-axis—the spacing between neighboring calcium ions in the HAp crystal (Fig. 1A, step 3). We connected these four segments using Ca-RFdiffusion(*32*) to generate the monomeric subunit and then applied C_3_ symmetry around the crystal c-axis to generate a C_3_ oligomer (Fig. 1A, steps 4-5). LigandMPNN(*22*) was used to design sequences that promote folding to the target structure while keeping the HAp interacting carboxylate residues fixed (Fig. 1A, Step 6). Designed trimers whose AlphaFold2(*33*) predicted structures deviated by less than 3 Å C_α_ Root Mean Square Deviation (RMSD) from the designed structures (Fig. 1A, Step 7) were selected for experimental characterization.

### C_3_ oligomer characterization and mineralization

73 designed C_3_ trimers were expressed in *Escherichia coli* and purified (*34, see methods*). Of these, 42 were produced at high levels, and six folded into the intended oligomeric structure as determined by mass spectroscopy, Size Exclusion Chromatography (SEC), Circular Dichroism (CD), and negative stain transmission electron microscopy (nsEM) (Fig S1-S14, Table S1-S2).

To evaluate mineralization activity, we mixed the oligomers with a supersaturated CaCl_2_/K_2_HPO_4_ solution. Attenuated Total Reflectance Fourier Transform Infrared Spectroscopy (ATR-FTIR) of the HAp1_C_3_ sample showed a strong signal at ∼1045 cm^-1^ and a weak signal at ∼1095 cm^-1^ (ν_3_ PO_4_^3-^) indicative of HAp formation(*35*) (Fig. S15A). This phosphate signal overlapped with that of commercial HAp nanoparticles, whereas control reactions lacking protein showed no such signal (Fig. S15A). The reactions with the other five C_3_ oligomers produced broader phosphate stretching signals with peak splitting in the 1000-1200 cm^-1^ region, indicating less crystalline or mixed calcium phosphate phases (Fig. S15B).

We further characterized mineralization with electron microscopy. Prior to mineralization, HAp1_C_3_ has a hollow ring structure matching the design model with a diameter of ∼9 nm (Fig. 2A). After mineralization, additional density in the middle of the ring is evident (Fig. 2B). We characterized the mineralized sample under transmission electron microscopy without negative stain (TEM) and observed ∼3.5 nm nanoparticles (Fig. 2C). High resolution transmission electron microscopy (HRTEM) revealed lattice fringes with d-spacings of ∼2.73 Å and mutual angles of ∼60°, matching the (300)-type reflections of HAp, consistent with the top-view orientation of the oligomers observed on the grids (Fig 2B), and the targeting {010} plane presented parallel to the protein-mineral interface. TEM screening of the other five C_3_ oligomers showed no mineral confinement (Fig. S16); thus, HAp1_C_3_ was selected for downstream investigations.

**Fig. 2.**
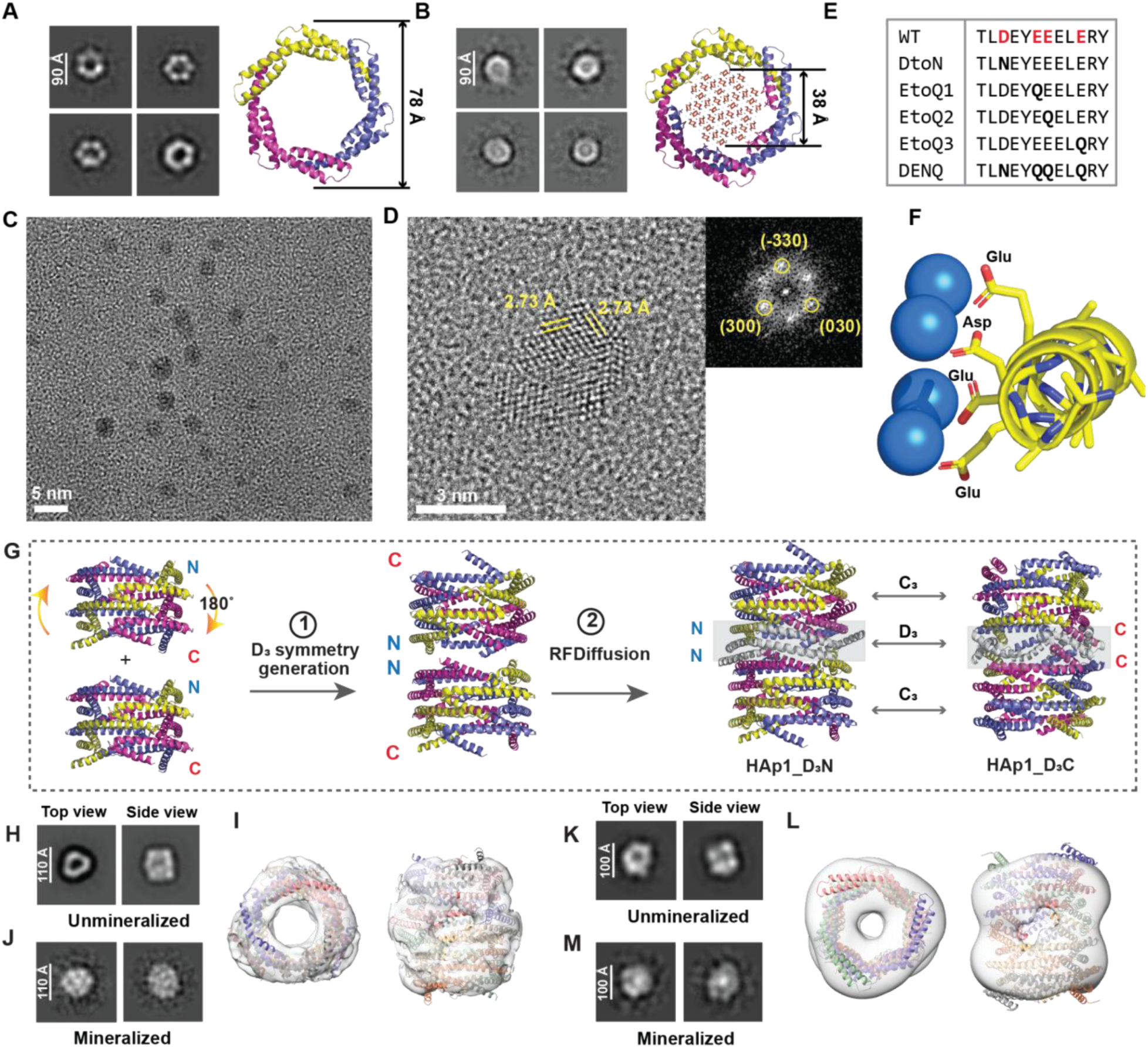
HAp crystal templating by the HAp1_C_3_ and HAp1_D_3_ designs. (**A**) Unmineralized HAp1_C_3_ nsEM 2D classes. (**B**) Mineralized HAp1_C_3_ cryoEM 2D classes (**C**) Mineralized HAp1_C_3_ TEM image. (**D**) Mineralized HAp1_C_3_ HRTEM. The top right panel shows the FFT of the nanocrystal. (**E**) Protein sequence of the repeated interfacial helix of HAp1_C_3_ and knockout mutants. Residues in bold indicate mutations. (**F**) Model illustration of designed carboxylates to interact with calcium from HAp. (**G**) HAp1_D_3_ oligomer design. The D_3_ interface does not incorporate lattice-matching sequences. (**H**) and (**K**) nsEM 2D class averages of HAp1_D_3_N and HAp1_D_3_C, respectively. (**I**) and (**L**) Design models fitted into a volume density map generated from nsEM results of HAp1_D_3_N and HAp1_D_3_C, respectively. The “pores” visible in the side views are at the D_3_. (**J**) and (**M**) nsEM 2D class of mineralized HAp1_D_3_N and HAp1_D_3_C, respectively. Raw micrographs are in Fig. S22.

HAp1_C_3_ has 4 repeats per monomer and each repeat has one aspartate and three glutamate residues designed to coordinate HAp calcium ions (Fig. 2F-G). To test their contribution to nucleation, five knockout mutants were generated with one or more carboxylates per repeat replaced with uncharged asparagine or glutamine (Fig. 2E and Table S1). All mutants retained the C_3_ oligomeric structure by SEC (Fig. S17A). ATR-FTIR analysis of nucleation reactions showed similar amide I (1600-1700 cm^-1^) and amide II (1500-1600 cm^-1^) signals in all reactions containing protein, whereas the protein-free control showed no such signals. Only the original design produced the strong phosphate stretching at ∼1063 cm^-1^ and a weaker signal at ∼1092 cm^-1^ (Fig. S17B, C), characteristic of biological apatite(*36*), demonstrating that the designed carboxylate residues are essential for mineralization.

### D_3_ oligomer design, characterization and mineralization

We next converted the HAp1_C_3_ oligomers into a D_3_ assemblies as a stepping stone to extended nanotubes and 2D arrays. To generate D_3_ assemblies, we stacked two C_3_ oligomers by arranging either their N-termini or C-termini against each other with C_2_ symmetry (Fig. 2G). We then used RFdiffusion(*20*) to extend the backbone under symmetry constraints to bridge the inter-ring gap, generating new D_3_ interfaces. The amino acid sequences at the new D_3_ Interface and adjacent regions of the structure were optimized by proteinMPNN(*23*), without consideration of HAp lattice matching. Sequences predicted by AlphaFold2(*33*) and AlphaFold3(*37*) to form D_3_ oligomers were selected for experimental characterization.

We experimentally screened 22 N-terminally stacked designs and 26 C-terminally stacked designs of D_3_ oligomers. Among them, designs HAp1_D_3_N and HAp1_D_3_C were chosen for more detailed characterization based on SEC (Fig. S18-19), MP (Fig. S12C-F), and nsEM (Fig. 2H-L, and Fig S20-21) results indicating the correct oligomeric assembly, alpha-helical secondary structure, and hollow ring morphology with shape matching the design.

We next tested the nucleation activity of HAp1_D_3_N and HAp1_D_3_C. Prior to mineralization, the designs present as hollow rings under nsEM (Fig. 2H and K). After mineralization, we observed black dots under TEM for both D_3_ oligomers (Fig. S22A and C). Staining the mineralized samples revealed filled rings (Fig. 2J and M; Fig. S22B and D), consistent with mineral occupying the protein cavity. HRTEM of the mineralized designs revealed crystalline lattice fringes matching with the HAp (040) lattice plane (Fig. S22E-F).

### Protein nanotube design and characterization

Unbounded protein nanotube architectures were designed by combining the N- and C-termini interfaces of HAp1_D_3_N and HAp1_D_3_C respectively (Fig. 3A) to generate HAp1_Nt (Fig. S23). Cryogenic transmission electron microscopy (cryoEM) micrographs of HAp1_Nt and 2D class averages showed extended nanotube structures with clearly defined secondary structure elements (Fig. 3B). 3D reconstruction of these particles generated a 3.08 Å cryoEM density map with well-resolved structural details, enabling unambiguous assignment of side-chain rotamers. Consistent with the design model, the density for HAp1_Nt represented a hollow nanotube structure containing 10-25 Å periodic pores which were incorporated during computational design to facilitate molecular transport of mineral precursors into the nanotubes (Fig. 3C and Fig. S24). The resulting atomic model of HAp1_Nt (Fig. 3D) is in close agreement with the design model, (Fig. 3E) with a total monomer Cα RMSD of ∼1.1 Å (Fig. 3F and Table S3).

**Fig. 3.**
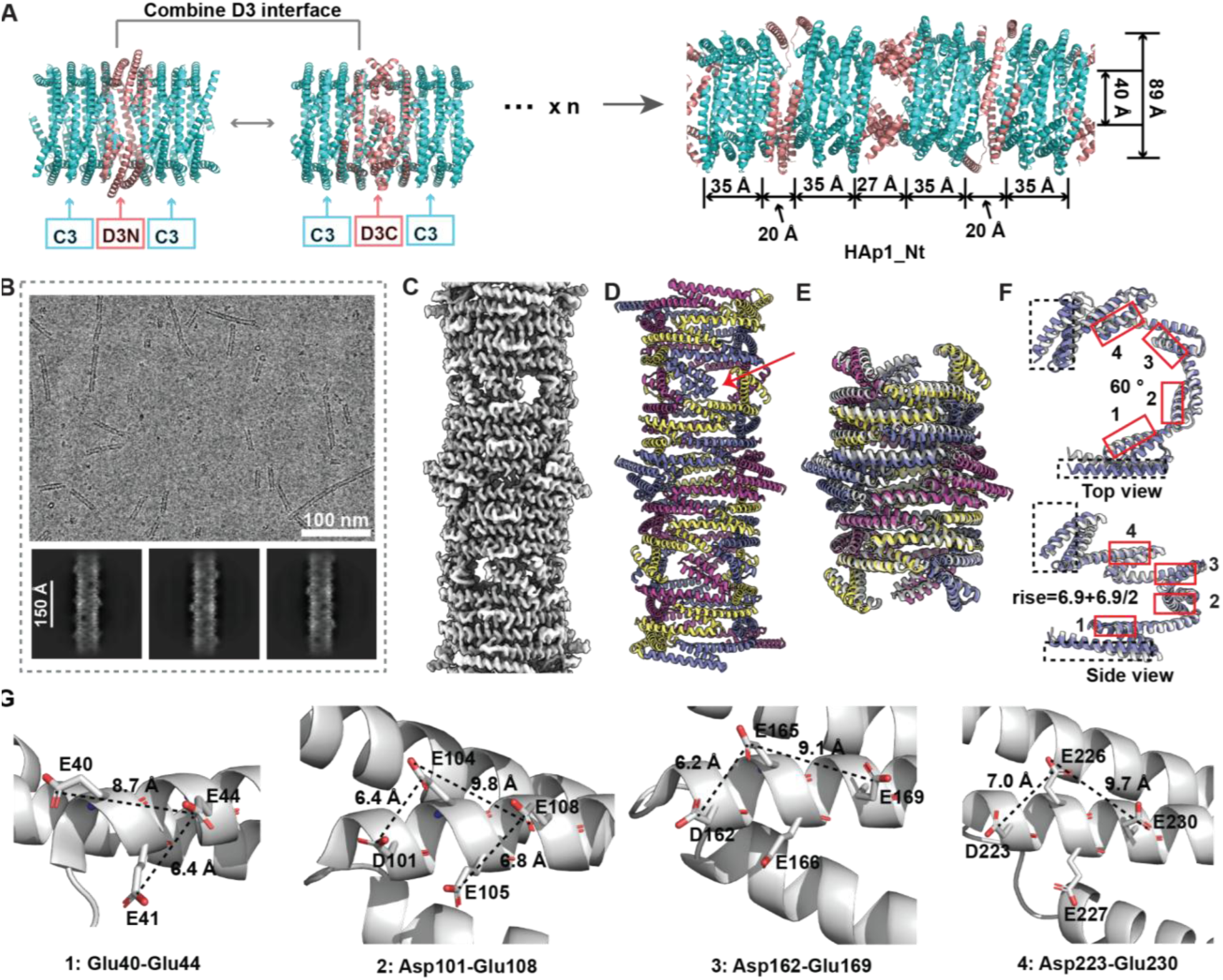
Design and cryoEM structure of the HAp1_Nt nanotube design. (**A**) Schematic of nanotube design showing the stacking of the lattice-matching region derived from the original C_3_ oligomer (cyan) and the non-lattice-matching D_3_ polymerization region (salmon). The resulting nanotubes have alternating lattice matching (3.5 nm) and non-lattice matching (2-2.7 nm) regions along the extended dimension. (**B**) cryoEM raw micrograph (top) and 2D class averages (bottom) of HAp1_Nt. (**C**) CryoEM density map. (**D**) CryoEM derived 3D structure. Arrow indicates periodic pores designed at D_3_ interfaces that may facilitate molecular transport. (**E**) Design model of the nanotube (grey) overlaps with the cryoEM structure (colored). (**F**) Monomer C_ɑ_ RMSD ∼1.1 Å. CryoEM structure is in purple and the design model in grey part. Red boxes indicate the initial mineral templating regions (see Fig 1A); black-dash boxes indicate non-nucleating D_3_ polymerization interfaces near the N-(1-39) and C-termini (276-332). (**G)** distance measurements between pairs in Glu40-Glu44, Asp101-Glu108, Asp16240-Glu169, and Asp223-Glu230.

The atomic model confirms the presence of four carboxylate-rich segments per monomer (Fig. 3F). The distances between carboxylate groups are 6.2–7.2 Å in one direction and 8.7–9.7 Å in the perpendicular direction, matching the spacing between calcium ions in the HAp {100}/{010}-type crystal lattice along the c-axis as 6.88 Å and a/b-axis as 9.42 Å, respectively (Fig. 3G). Each successive segment is related by ∼60° and shifted 10.32 Å along the nanotube axis (1.5× the HAp c-axis repeat), matching the helical symmetry of HAp. The three-fold arrangement of monomers in the nanotube mirrors the hexagonal geometry of the HAp crystal face.

### Nanotube mineralization

We mixed HAp1_Nt with CaCl_2_ and K_2_HPO_4_ and observed predominantly extrafibrillar mineralization (*assay 1*) (Fig. 4A vs Fig. S25A), with nanotubes exhibiting polycrystalline features (Fig. S25B). Polymer-induced liquid phase precursors have been demonstrated to be critical for enhancing collagen intrafibrillar mineralization(*4*, *38–41*), we therefore prepared amorphous calcium phosphate (ACP) precursors by mixing CaCl_2_/K_2_HPO_4_ with polyaspartic acid (pAsp) (*assay 2*). After incubating the pAsp-ACP precursor with HAp1_Nt, we observed electron-dense filaments with periodic banding (Fig. 4B). Staining of mineralized HAp1_Nt revealed strongly contrasting density within the nanotube lumen (Fig. 4C). HRTEM of intrafibrillarly mineralized nanotubes revealed HAp nanocrystals exposing (002) facets perpendicular to the designed {010} nucleation plane (Fig. 4D and Fig. S25C), matching the dimensions of HAp nanocrystals confined in the nanotube lumen and the orientation of the nanotubes which lie perpendicular to the beam on the TEM grids. The crystallinity is not uniform with amorphous regions coexisting along the nanotube axis. Precursor-only control showed no crystallinity (Fig. S25D). Volume density maps of mineralized HAp1_Nt with pAsp (unstained TEM) revealed coherent density of ∼40 Å diameter, matching the dimension of designed protein cavity, consistent with mineral filling the nanotube interior while the overall map diameter expands to 107 Å relative to the unmineralized protein map diameter (92 Å), showing heterogeneous features coating the nanotube exterior surface (Fig. 4E). Combined with HRTEM (Fig. 4D), these reconstructions (Fig. 4E) suggest minerals occupying the nanotube lumen and nonspecific precursors associating with the exterior of the nanotube.

**Fig. 4.**
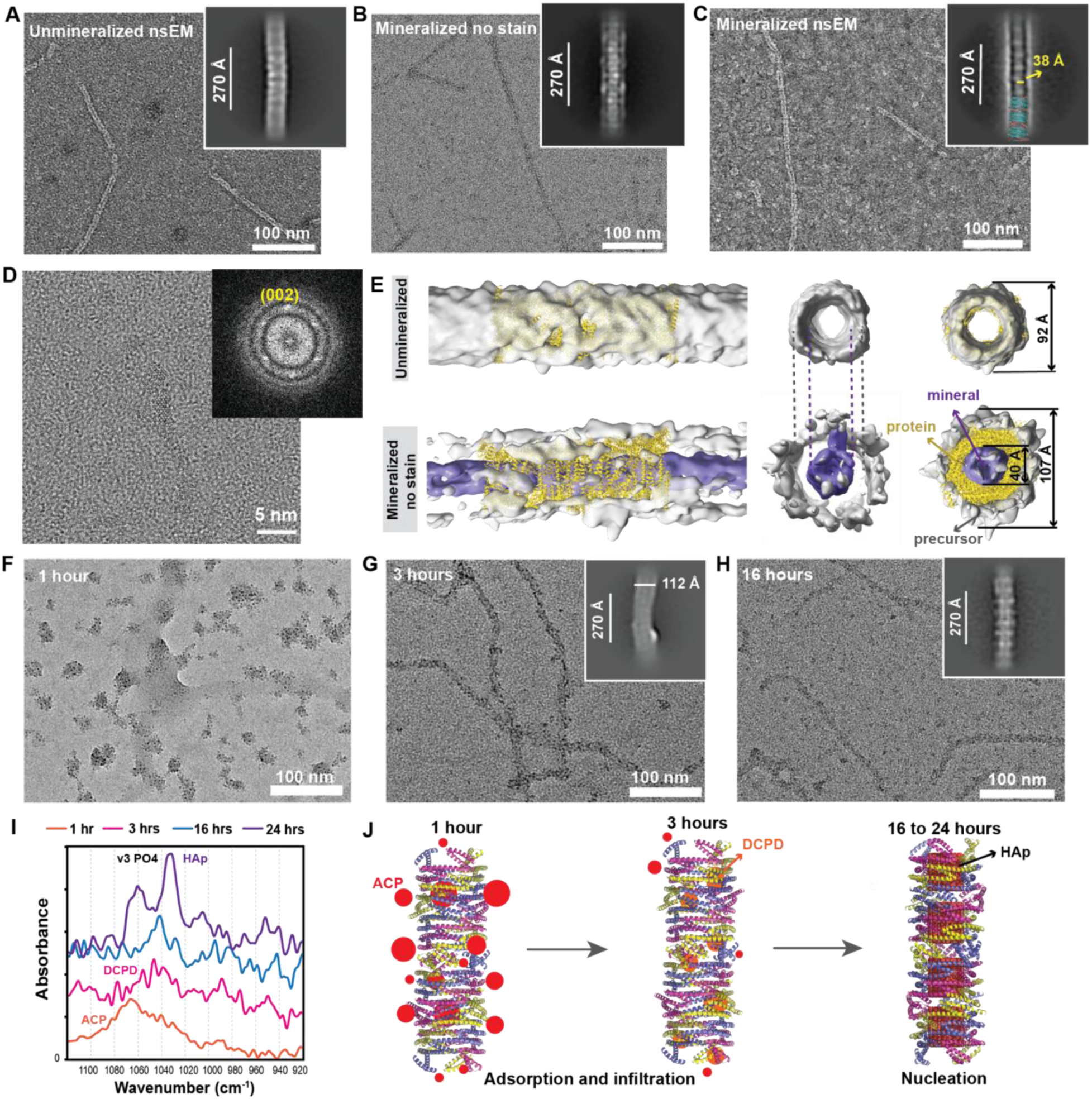
Time-course of HAp mineralization within HAp1_Nt nanotubes. **(A)** nsEM of unmineralized nanotubes, with 2D class average in inset. (**B**) TEM of HAp1_Nt after 24 hours of mineralization in presence of 180 µg/mL 27kda pAsp *(assay 2*). (**C**) nsEM of the sample in panel B. The inset shows the 2D class average superimposed on the HAp1_Nt cryoEM structure. (**D**) HRTEM of mineralized HAp1_Nt sample in panel B, with FFT in inset. (**E**) The cryo-EM structure of apo HAp1_Nt fitted into nsEM-derived density map of unmineralized nanotubes in panel A and TEM-derived density map of mineralized nanotubes with pAsp in panel C. (**F-H**) TEM of HAp1_Nt mineralized after 1 hour, 3 hours, and 16 hours mineralization, respectively. The insets are 2D class averaging. (**I**) ATR-FTIR characterization of HAp1_Nt mineralized after different mineralization times. (**J**) schematic illustration of the nanotube mineralization process.

Scanning Transmission Electron Microscopy - Energy Dispersive X-Ray Spectroscopy (STEM-EDX) of mineralized HAp1_Nt revealed co-localization of calcium and phosphorus signals within nitrogen-rich protein nanotube regions (Fig. S26). Cryo-EM of mineralized HAp1_Nt showed increased filament electron density relative to unmineralized controls (Fig. S27 vs. Fig. 3B), consistent with mineral association along the scaffold. Heterogeneous mineral and precursor distribution precluded 3D reconstruction.

We next studied the kinetics of intrafibrillar mineralization with HAp1_Nt by examining the reactions at earlier timepoints. After 1 hour, TEM revealed nanoparticle clusters (Fig. 4F) that resemble the morphology of ACP nanoparticles observed in the literature(*38*), and the ATR-FTIR spectrum showed a broad peak at 1020–1100 cm^-1^, confirming ACP formation(*42*) (Fig. 4I and Fig. S28A). After 3 hours, filamentous structures decorated with dark dots appeared (Fig. 4G), suggesting progressive adsorption of ACP onto the nanotubes and continued with infiltration (Fig. 4J). The ATR-FTIR spectrum showed multiple split bands at 1040-1056 cm^-1^ (ν_1_ PO_4_), 1080 cm^-1^ (ν_3_ PO_4_) and 990 cm^-1^ (HPO_4_^2-^) (Fig. 4I), characteristic of dicalcium phosphate dihydrate (DCPD or brushite) (*42–44*), suggesting conversion of ACP to DCPD. By 16 hours, the surface-bound particles decreased, leaving filamentous features (Fig. 4H), likely corresponding to further infiltration of precursors into the nanotube interiors (Fig. 4J). The ATR-FTIR spectrum (Fig. 4I) showed that the initially split PO_4_ bands evolved into two dominant peaks at ∼1044 and ∼1029 cm⁻¹, characteristic of poorly crystalline HAp.(*35*) After 24 hours, the filaments became thinner, consistent with densification of the mineral phase by dehydration of DCPD and HAp nucleation (Fig. 4B, D and Fig. 4J). The ATR-FTIR spectrum showed that the major peak at ∼1040 cm^-1^ intensified (Fig. 4I), confirming crystalline HAp formation(*35*). Negative controls lacking protein showed no HAp formation (Fig. S28B and S29).

We investigated what aspects of HAp formation are controlled by the designed interface by characterizing the interface knockout mutants (Table S1). While several of the mutations interfered with fiber formation, HAp1_Nt_DtoN formed filaments similar to the original design (Fig. S30A). ATR-FTIR results showed multiply split ν_3_ PO_4_ bands matching β-tricalcium phosphate (β-TCP)(*45*) with broad ACP bands at 1 and 3 hours, and better-resolved but phase-stalled β-TCP bands at 16 and 24 hours (Fig. S30 B and C). TEM showed low-electron density filamentous features after 16 and 24 hours of reaction but without mineral confinement in the nanotube (Fig. S30 D-G). These results suggest that although the HAp1_Nt_DtoN can template precursors, it fails to direct phase conversion to HAp or achieve spatial confinement of minerals, suggesting that the designed interface enhances the late-stage HAp conversion.

Liquid-phase atomic force microscopy (LP-AFM) was used to characterize the mechanical properties of HAp1_Nt before and after mineralization. Topographic imaging confirmed that both unmineralized and mineralized samples exhibit nanotubular structures, consistent with the TEM observations (Fig. 5A, B). Young’s modulus mapping indicated moduli of approximately 5.1 MPa and 16.8 MPa for unmineralized and mineralized nanotubes (Fig. 5C and Fig. S31A, B). A relatively large standard deviation for the mineralized sample likely reflects the coexistence of mineralized and unmineralized regions within individual nanotubes. The ∼3-fold increase in Young’s modulus upon mineralization is consistent with HAp mineral reinforcement of the protein scaffold (Fig. S31C-E).

**Fig. 5.**
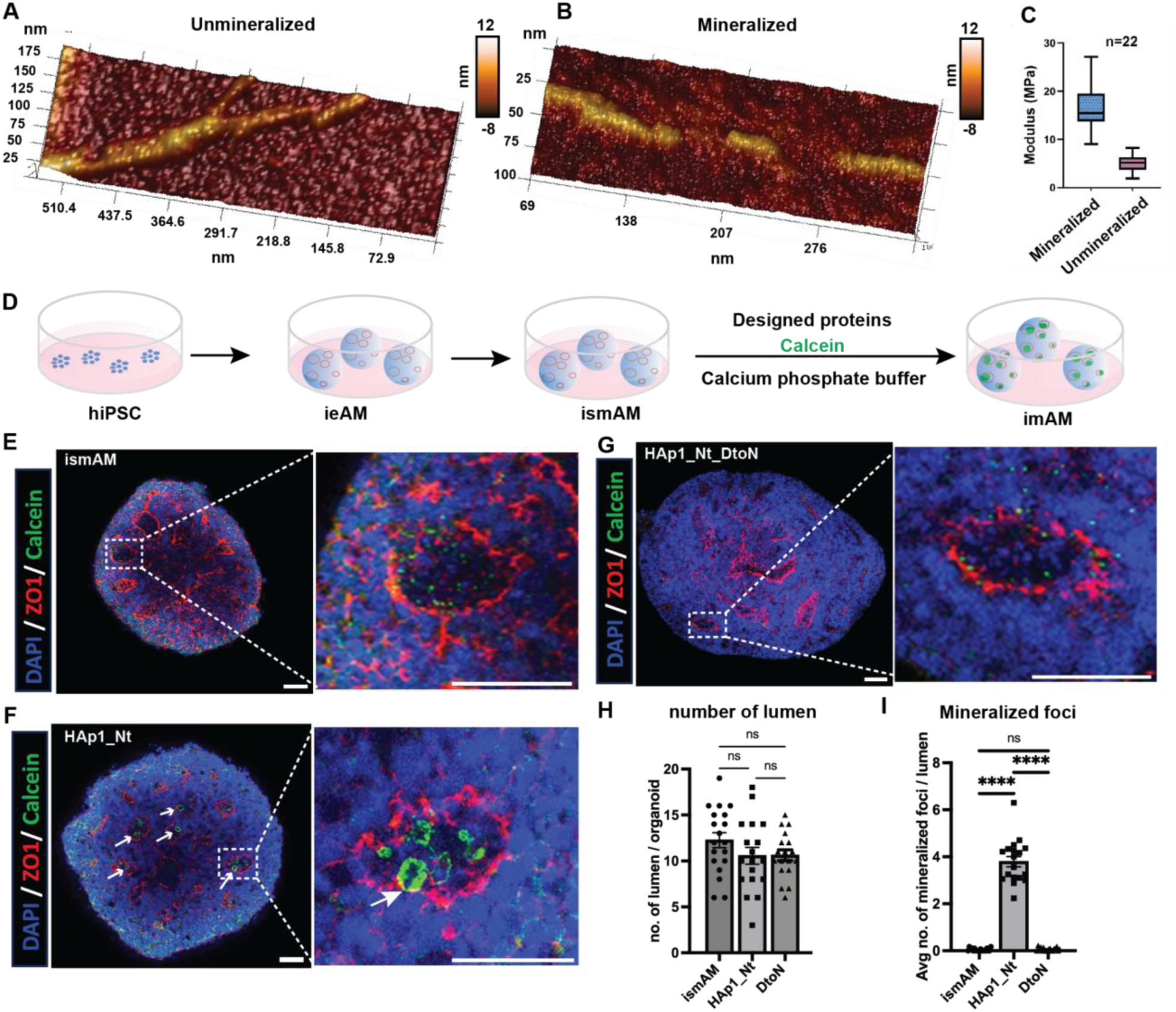
HAp1_Nt mineralization increases nanotube stiffness and promotes mineral deposition in dental organoids. (**A, B**) LP-AFM topography images of unmineralized and mineralized HAp1_Nt nanotubes, respectively. (**C**) Elastic modulus distributions calculated from Fig. S31A, B. (**D**) Schematic of the ismAM (induced secretory-stage maturation ameloblast) dental organoid mineralization assay showing the human induced pluripotent stem cell (hiPSC) differentiation trajectory from ieAM (induced early ameloblast) to ismAM and, following HAp1_Nt-driven mineralization, to imAM (induced mature ameloblast). (**E–G**) Confocal images of ismAM organoids cultured without added protein (**E**), with HAp1_Nt (**F**), or with HAp1_Nt_DtoN (**G**). Nuclei (DAPI, blue), tight junctions and lumen boundaries (ZO-1, red), and mineralized foci (Calcein, green) are shown. Scale bars: 50 μm. (**H**) Quantification of lumen number per organoid across treatment groups. (**I**) Quantification of average number of mineralized foci per lumen across treatment groups (p < 0.0001).

To evaluate biological relevance, we tested HAp1_Nt-driven mineralization in human induced pluripotent stem cell (hiPSC)-derived dental organoids differentiated to the induced secretory-stage maturation ameloblast (ismAM) (Fig 5D); in these organoids the apical side of the ameloblasts, which secrete the proteins inducing mineralization during normal tooth formation, surround a lumen like region in the organoids. We investigated mineralization in the presence of HAp1_Nt, calcein, and calcium phosphate. Compared to untreated controls (Fig. 5E), HAp1_Nt-treated organoids showed mineral deposition at the apical side of the epithelial ameloblasts (Fig. 5F), a site reflecting native enamel protein secretion and mineralization geometry (*46*). This was accompanied by a significant increase in mineralized foci per lumen of the epithelial layer (p < 0.0001, Fig. 5I) and Alizarin Red S (ARS) staining intensity (p < 0.0001, Fig. S32A), while lumen number per organoid remained unchanged (Fig. 5H), confirming that HAp1_Nt promotes spatially directed mineralization without disrupting organoid architecture. ATR-FTIR of mineralized HAp1_Nt-treated organoids revealed a ν_3_ PO_4_ band absent in untreated controls, consistent with calcium phosphate mineral deposition (Fig. S32B). The HAp1_Nt_DtoN mutant (Fig. 5G) showed no detectable calcification, demonstrating that mineralization-associated maturation depends on the designed nucleation interface rather than nonspecific protein effects.

### 2D array design, characterization and mineralization

We next explored lateral extension of the D_3_ oligomer to form a two-dimensional (2D) protein lattice. We positioned two HAp1_D_3_N oligomers side by side, thereby creating a D_2_ symmetry between the two oligomers (Fig. 6A)(*47*). We then used RFdiffusion(*20*) to generate inter-oligomer contacts (Fig. 6B) and ProteinMPNN(*23*) to design amino acid sequences compatible with the new backbones (Fig. 6C). The resulting D_2_ interface features hydrophobic interactions near the N-termini and salt bridges around the central contact region; geometric propagation generates the higher-order assembly (Fig. 6D). We screened 66 array designs among which 44 were soluble and were characterized with nsEM. HAp1_Ar had the most extended lattice pattern matching the design (Fig. 6E). The sharp peaks on the binned Fast Fourier Transform (FFT) confirm formation of ordered extended assemblies (Fig. 6E).

**Fig. 6.**
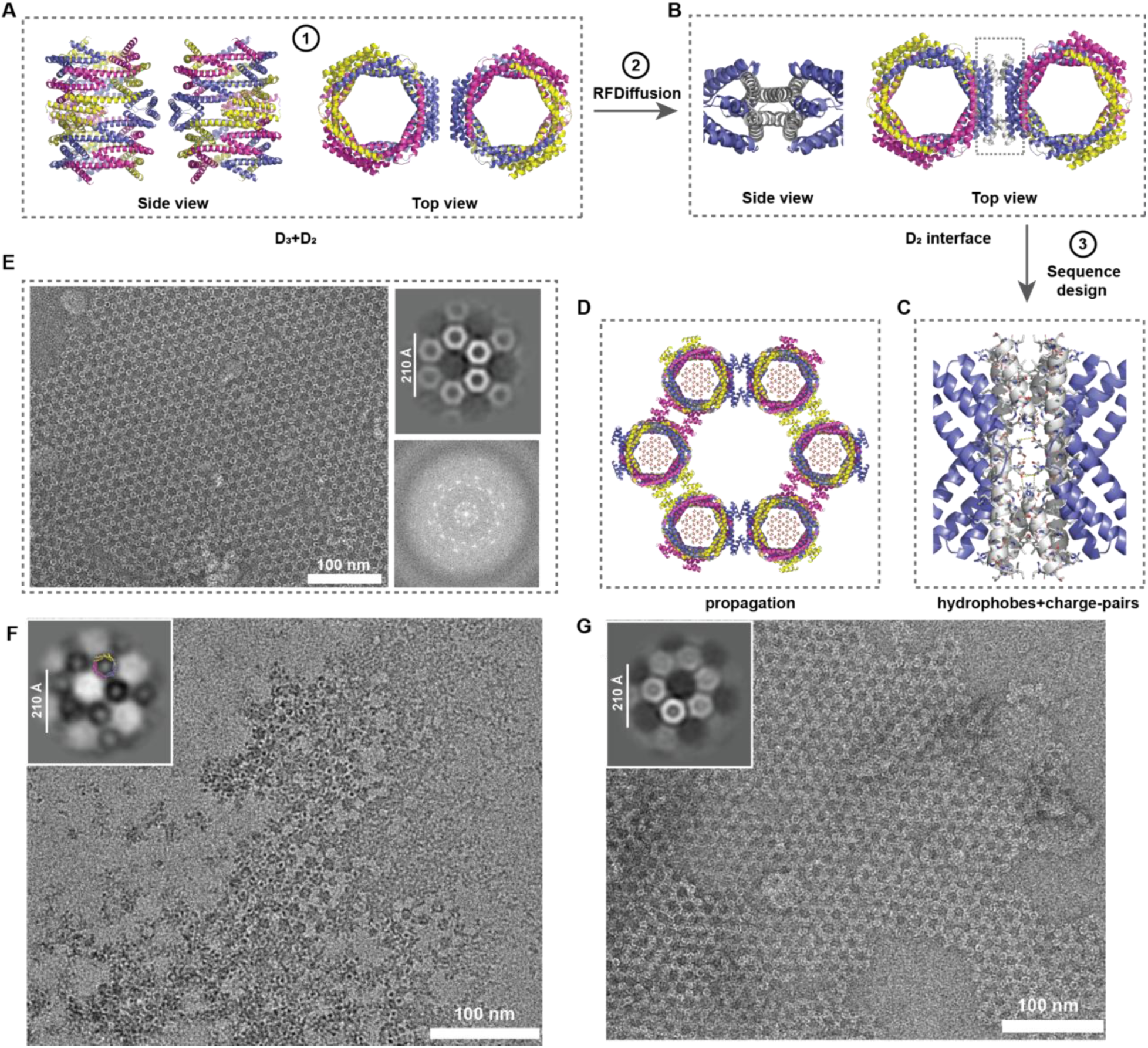
Two-dimensional array HAp1_Ar design, characterization and mineralization. (**A**) Two HAp1_D_3_N oligomers were arranged to generate D_2_ symmetry. (**B**) D_2_ interface generation using RFDiffusion (grey part). (**C**) Side chain interactions of D_2_ interface. (**D**) Propagation models with templated HAp nanocrystals generated in PyMOL(*51*). (**E**) and (**G**) nsEM results of the unmineralized and mineralized array, respectively. (**F**) TEM of mineralized array. The inset is 2D class averaging overlapping with the D_3_ design model.

We explored the mineralization activity of the HAp1_Ar array (assay 2). After 24 hrs reaction, we observed formation of nanocrystals matching the lattice under TEM (Fig. 6F). nsEM of the mineralized array showed density inside the D_3_ oligomers (Fig. 6G), suggesting templated nucleation; in contrast no particles were observed in the large hexameric regions between the D3 oligomers. As the amino acid sequence was designed to promote mineralization on the inside of the D3 assemblies, but not in the hexameric regions between them, these results provide a further demonstration of sequence specific templating of HAp formation. ATR-FTIR of mineralized HAp1_Ar revealed a strong ν_3_ PO_4_ band at ∼1020 cm^-1^ characteristic of HAp(*35*), which was absent in protein-free controls (Fig. S33).

## Discussion

Our results demonstrate that HAp-templating interfaces can be computationally designed and used to both template and confine HAp growth at specific locations within designed protein assemblies. This ability to confine mineral growth goes beyond previous efforts with calcium carbonate, where designed proteins templated but failed to confine growth, resulting in large unbounded crystals with little memory of the shape of the nucleating protein(*17*). Structural insight into this confinement is provided by the cryo-EM structure of the apo HAp1_Nt, which shows that sidechain positions at the designed interface geometrically match the HAp crystal lattice. Mutation of the designed mineral templating sidechains at this interface abolishes the phase conversion from precursors to HAp, confirming that geometric positioning of functional groups is critical for mineralization.

The mineralization observed in our synthetic systems parallels that of collagen *in vitro* mineralization in key respects. Similar to collagen, we observed extrafibrillar and enhanced intrafibrillar mineralization of HAp1_Nt without and with polymer-induced precursors, respectively(*38*) as well as a heterogeneous, multistage nucleation process(*2*, *48*). While positively charged regions in collagen recruit negatively charged ACP clusters(*49*), our designed nanotubes feature negatively charged interfaces to recruit Ca^2+^ ions that could subsequently attract phosphate-rich ACP clusters. Although differences in scaffold dimensions preclude direct kinetic comparison, the earlier appearance of HAp in HAp1_Nt nanotubes relative to collagen fibrils(*49*) suggests the designed nucleation surface is at least comparably effective at templating mineral conversion.

## Acknowledgements

We thank S. Woodbury for helping set up RFDiffusion2 scripts. LTY thanks Dr. J. Chen and the rest of Baker lab members for general discussions. We thank IPD Core Research members X. Li and C. Möller for mass spectrometry data collection and processing. We thank HHMI Janelia Research Campus for cryoEM data collection; J. Quispe, F. Zhang and W. Jiang for the management of Arnold and Mabel Beckman CryoEM Center at University of Washington. We thank IPD Yeast Production/Processing Center members for providing us with LM670 vectors. We thank the IPD lab management team led by K. VanWormer, IPD IT team led by L. Goldschmidt, IPD Electron Microscopy Research Core (EMRC) led by A. Borst for technical support. Part of the nanotubes and 2D lattices designs (HP) as well as the Liquid-phase AFM and associated mechanical mapping (XL, SZ) were based upon work supported by the US Department of Energy, Office of Science, Office of Basic Energy Sciences, as part of the Energy Frontier Research Centers (EFRC) program: CSSAS – The Center for the Science of Synthesis Across Scales under Award Number DE-SC0019288 and at Pacific Northwest National Laboratory under contract FWP 72448. PNNL is a multiprogram national laboratory operated for DOE by Battelle under Contract No. DE-AC05-76RL01830. STEM/EDS characterization performed at PNNL was supported by the U.S. DOE BES, Material Sciences and Engineering Division, Synthesis and Processing Science program, FWP 67554 (D. Li). Part of the TEM experiments were conducted at the Molecular Analysis Facility, a National Nanotechnology Coordinated Infrastructure site at the University of Washington which is supported in part by the National Science Foundation (grant NNCI-1542101), the University of Washington, the Molecular Engineering & Sciences Institute, the Clean Energy Institute, and the National Institutes of Health.

## Author contributions

Conceptualization: LTY, HP, NPB, DB

Methodology: LTY, HP, XL, NPB, AJB, CW, KDC, APP, PK

Investigation: LTY, HP, XL, AJB, YL, PK, RDK, APP, DL, SM

Data processing: LTY, XL, APP, AJB, CW, KDC

Visualization: LTY, XL, APP, AJB

Funding acquisition: HR-B, SZ, JDY, DB Supervision: HR-B, SZ, JDY, DB

Writing – original draft: LTY, HP, XL, AJB, JDY, DB

Writing – review & editing: All authors

Competing interests

Authors declare that they have no competing interests.

Data, code, and materials availability

All data are available in the main text or the supplementary materials. CryoEM structure of the HAp1_Nt was deposited in the Protein Data Bank under accession number 9ZNK.

Scripts for RFdiffusion are available at https://github.com/RosettaCommons/RFdiffusion. Scripts for ProteinMPNN and LigandMPNN are available at https://github.com/dauparas/ProteinMPNN and https://github.com/dauparas/LigandMPNN, respectively. Scripts for AlphaFold2 and AlphaFold3 are available at https://github.com/google-deepmind/alphafold and https://golgi.sandbox.google.com/ respectively. All detailed scripts, designed structures in pdb format and designed sequences in fasta files in this work are publicly available at Zenodo(*50*). All raw data of the plots presented in this work are available in the Data S1 file.

## License Information

Copyright © [xxxxxxxxxxxxxx publication year] the authors, some rights reserved; exclusive licensee American Association for the Advancement of Science. No claim to original US government works. https://www.science.org/about/science-licenses-journal-article-reuse. This article is subject to HHMI’s Open Access to Publications policy. HHMI lab heads have previously granted a nonexclusive CC BY 4.0 license to the public and a sublicensable license to HHMI in their research articles. Pursuant to those licenses, the Author Accepted Manuscript (AAM) of this article can be made freely available under a CC BY 4.0 license immediately upon publication.

