## Supplementary Materials for "Templating and confining calcium phosphate mineralization within designed protein assemblies"

Science
AAAS

### Supplementary Materials for

#### Templating and confining calcium phosphate mineralization within designed protein assemblies

Le Tracy Yu[1,2], Xinqi Li[3,4,10], Harley Pyles[1,2,10], Anjali P. Patni[5,6,7], Andrew J. Borst[1,2], Neville P. Bethel[8], Paul S. Kwon[1,2], Connor Weidle[1,2], Ryan D. Kibler[1,2], Kenneth D. Carr[1,2], Yulai Liu[1,2], Dongsheng Li[4], Stanislav Moroz[1,2], Hannele Ruohola-Baker[5,6,7], Shuai Zhang[3,4], James De Yoreo[3,4], David Baker[1,2,9]*

**Affiliations:**

[1]Department of Biochemistry, University of Washington, Seattle, WA, USA

[2]Institute for Protein Design, University of Washington, Seattle, WA, USA

[3]Department of Materials Science and Engineering, University of Washington, Seattle, WA, USA

[4]Physical Sciences Division, Pacific Northwest National Laboratory, Richland, WA, USA

[5]Department of Oral Health Sciences, University of Washington, School of Dentistry, Seattle, WA, USA

[6]Department of Biochemistry, University of Washington, School of Medicine, Seattle, WA, USA

[7]Institute for Stem Cell and Regenerative Medicine, University of Washington, School of Medicine, Seattle, WA, USA

[8]Department of Chemistry & Biochemistry, University of California San Diego, San Diego, CA, USA

[9]Howard Hughes Medical Institute, University of Washington, Seattle, WA, USA

[10]These authors contributed equally to this work

**This PDF file includes:**

Materials and Methods
Supplementary Text
Figs. S1 to S33
Tables S1 to S3
References 50-60

**Other Supplementary Materials for this manuscript include the following:**

Data S1

Materials and Methods
Scripts for RFdiffusion are available at https://github.com/RosettaCommons/RFdiffusion. Scripts for ProteinMPNN and LigandMPNN are available at https://github.com/dauparas/ProteinMPNN and https://github.com/dauparas/LigandMPNN, respectively. Scripts for AlphaFold 2 and AlphaFold3 are available at https://github.com/google-deepmind/alphafold and https://golgi.sandbox.google.com/ respectively. All detailed scripts for this design pipeline are publicly available at Zenoto(50).

***Computational Design of $C_3$ Oligomers***

*HAp crystal slabs generation*

HAp crystal structure is obtained from Crystallography Open Database (COD). HAp crystal slabs with specific facets were generated with Vesta.

*Tip atom generation*

Scripts (will be released later) were run to convert phosphates surrounding targeting calcium to be carboxylates.

*Tip atom selection*

For the design campaigns in this study, we selected four neighboring carboxylate residues at a time for RFD2. Theoretically, more carboxylate residues with different geometry can be selected. These carboxylates can be either aspartate or glutamate. After rationally selecting the tip atoms of interest, we ran "gen_inputs" scripts to generate all permutations between aspartate residue (Asp) and glutamate residue (Glu) of these tip atoms, and saved these residues and the HAp target into new pdb files. These pdb files were inputs for RFD2.

*Rosettafold Diffusion2 (RFD2)*

For diffusion, we sampled motif length between 10 to 30 residues which is the "length_range" information in the scripts. Each one residue increment will be a new job. "contig_stoms" information in the scripts refers to the atoms that users want to fix. In this study, we fixed "OE1, OE2 CD" for glutamate residue and "OD1, OD2, CG" for aspartate residue. We did two structure outputs per job.

*Motif propagation and symmetrize*

After diffusion, we ran "gen_seeds.py" scripts. This script identifies clashes between protein structures and clashes between protein and HAp crystal targets and removes those outputs. The outputs that did not have clashes were then aligned back to the HAp crystal target and propagated and symmetrized. In the propagation section, we did 60° rotation and 10.32 Å rise each propagation. These values were determined based on the target geometry. The "--ctot" value was 3 for designing $C_3$ oligomers. Users get to change this value to, for example 6, for $C_6$ oligomer design.

For targets without simple symmetry, we provided here the "propagate_and_symmetrize.py" scripts to allow users to propagate motifs based on residue positions. Users specify the starting position of the motif and the end position of the motif by entering the residue number of the tip atoms.

*Backbone generation with Ca-RFdiffusion(31, see supplementary information)*

We ran "gen_tasks.py" scripts to generate the commands for Ca-RFdiffusion. We did 25-60 residues length for diffusion. Users can customize these values with "--repeat_length_upper" and "repeat_length_lower" arguments.

*Inspect diffusion outputs*

We ran “get_breaks”.py to filter diffusion outputs based on their secondary structures. It requires structures to have no chain breaks or discontinuous secondary structures. It requires at least 75% of the protein backbone to be structured, either helix (H) or beta sheets (E). Structures that failed these requirements got filtered out.

*LigandMPNN*

We first ran “combine_pdb.py” to combine protein structure with target ligand, in this case HAp crystal. We then run the “ligandmpnn.ipynb” scripts to design protein sequences. We did “--temperature 0.3” “--batch size 5” “number_of_batches 5” and “--bias_AA ‘R:-0.9’” for the designs. The “--temperature” argument refers to the diversity of amino acid sequences, the lower, the higher diversity. “--batch size” and “number_of_batches” arguments decide the number of sequences generated, we did a total of 25 sequences per structure. “--bias_AA” determines amino acid preference, and we biased against arginine in those design campaigns.

*AlphaFold2*

We did AlphaFold2 multimer folding with three numbers of folding cycles. Designs with less than 3 Å RMSD deviation between the prediction and the design models were filtered out for experimental test.

***Computational Design of $D_3$ Oligomers and Nanotubes***

*$D_3$ symmetry generation*

We first saved chain A of the $C_3$ oligomer HAp1_$C_3$ into a new pdb file. The chain A was already centered at x=0, y=0, z=0 in PyMOL(*51*). Users will have to center their structure prior to geometry generation. We then used Rosetta to generate $D_3$ symmetry using the chain A as input. We sampled the distance between the $D_3$ interface to be 5 to 20 Å, the rotation between the interface to be 0°, 30°, 60°, and 90°. We ran “sample_dihedral_arrangments.ipynb” scripts to generate commands to sample distance and rotations for generating $D_3$ symmetry.

We then did RFdiffusion similarly as described above to generate new protein interfaces to connect the $C_3$ oligomers. We sampled diffusing amino acid lengths of 20-40.

*ProteinMPNN*

We used proteinMPNN to design the sequences of the newly generated $D_3$ interface. We ran “parse_pdb.py” scripts to generate commands for MPNN. Specifically, we used the “def_fix” module to fix the protein sequences from the original $C_3$ oligomer. Specifically, when designing structures that pack $C_3$ oligomers against their N-termini, sequences containing three functional nucleation motifs near the C-termini were fixed, and vice versa. This design process necessitated redesigning the protein fragment containing one functional nucleation motif to enable its connection with the protein-protein polymerization interface, resulting in the modification of one nucleation motif per chain in the final $D_3$ nanotube design (3 original motifs and one modified motif per chain compared to 4 in the original $C_3$ oligomer). Although this redesign did not adversely affect mineralization activity, as demonstrated experimentally, the lattice-matching constraint was not satisfied during this design process and we acknowledge this as a limitation of the current design approach.

***Computational Design of 2D Arrays***

*$D_2$ symmetry generation*

We saved chain A of the HAp1_$D_3$N oligomer into a new pdb file as the input for generating $D_2$ symmetry. We used Rosetta to generate a $D_2$ interface similarly as described above. We sampled 5-20 Å distance between the $D_2$ interface with no rotation.

***Protein Expression and Harvest***

Buffer Recipes:

Lysis buffer: 25 mM Tris-HCl, 300 mM NaCl, 30 mM Imidazole, pH 8
Elution buffer: 25 mM Tris-HCl, 300 mM NaCl, 500 mM Imidazole, 5 mM EDTA, pH 8
SEC running buffer/dialysis buffer: 25 mM Tris-HCl buffer, 150 mM NaCl, pH 7.4
All buffers were filtered through a 0.22 μm filter prior to use.

All chemicals and supplies were ordered from Thermo Fisher unless specified otherwise.
The selected designs were reverse translated into DNA sequences. Gene fragments were ordered from IDT or Twist Bioscience. Gene fragments were cloned into a pET29b+ vector between NdeI and XhoI sites with a C-terminal hexahistidine tag using Golden Gate assembly kit (New England Biolabs, Ipswich, MA). The cloned genes were transformed into *Escherichia coli* (*E. coli*) BL21 strain. Designs identified as active during screening were ordered from Genscript as sequence-verified clonal genes and used for subsequent protein expression.

*Small-scale Protein Expression and Harvest*
After transformation, designs were expressed in 4 mL autoinduction Terrific Broth II (TBII) media in 96 deep-well plates at 37 °C for 20-24 hours. The plates were shaken at ~1000 revolutions per minute (RPM) in a plate shaker (Southern Labware). After expression, cells were pelleted by centrifuging the plates at ~4000 relative centrifugal force (rcf) for 15 minutes. The supernatants were discarded. Cell pellets were then resuspended into 200 μL of lysis buffer with addition of 1X bugbuster, 0.1 mg/mL lysozyme, 10 μg/mL DNAse, and protease inhibitor (1 tablet/50 mL). Cell pellets were lysed by shaking at room temperature at 1000 rcf for 15 minutes. Lysed cells were centrifuged at 4000 rcf for 20 minutes and the supernatants were collected and purified in a 96-well format using magnetic nickel-NTA (Ni-NTA)) resins. In detail, in each well in a 0.7 mL semi-deep 96-well plate, add 20 μL of magnetic Ni-NTA resin (pre-washed with lysis buffer) and 200 μL of the supernatant from the cell lysate. The plates were covered with plate seal and shaken at ~1000 rcf at room temperature for 15-30 minutes to allow binding. The magnetic beads were then extracted with a magnetic extractor and resuspended in a lysis buffer for wash. After washing 3 times, the magnetic Ni-NTA beads were resuspended in 120 μL of the elution buffer for 15 minutes to allow elution. The magnetic beads were extracted after elution. And the eluents were subject to down-stream purification and characterization.
*Medium-scale Protein expression and Harvest*
Designs were expressed in 50 mL autoinduction TBII media in 250 mL baffled erlenmeyer flasks. The flasks were shaken at ~225 rcf in New Brunswick Innova® 44 shakers at 37 °C for 4 hours and then at 18 °C for 20 hours. Cultures were transferred into 50 mL conical tubes and centrifuged at 4000 rcf for 15 minutes to pellet cells. Cell pellets were resuspended in a 10 mL lysis buffer containing 0.1 mg/mL lysozyme, 10 μg/mL DNAse, and protease inhibitor (1 tablet/50 mL). Cells were lysed by sonication with cooling using a 4 horn tip sonicator (QSonica Sonicators, CT, USA) for 5 minutes at 10 seconds on and off pulse and 65% amplitude.
Lysed cells were centrifuged at 14000 rcf for 20 minutes and the supernatants were collected and purified using Ni-NTA resin. In detail, 2 mL of Ni-NTA resin was added into 10 mL chromatography columns (BIO-RAD Econo-Pac gravity flow columns) to result in a 1 mL of bead volume. The beads were washed one time with 3 column volume (CV) of Milli-Q (MQ) water and then 10 CV of lysis buffer. The supernatants of cell lysate were added into the columns and allowed for binding for 10 minutes. The resins were then washed 3 times with 10 CV of lysis buffer each. The proteins were pre-eluted with 600 μL of elution buffer and then eluted and collected with 1.5 mL of elution buffer.

##### *Protein Purification with Size Exclusion Chromatography*

Samples were loaded to an Agilent 1260 Infinity II LC System equipped with Superdex 200 Increase 10/300 GL (Cytiva) columns (medium scale), or Superdex 200 Increase 5/150 GL (Cytiva) columns (small scale). Samples were run with a flow rate of 0.75 mL/minute and signals were monitored at 220 nm and 280 nm.

##### *Dialysis*

After Ni-NTA column purification, designed nanotubes or arrays were loaded into D-Tube Dialyzer units (MWCO 6-8 kDa, Sigma-Aldrich) and dialyzed against SEC buffer to remove small molecule impurities. Dialysis was performed in three cycles, each with fresh buffer replacement.

##### *Higher-order assembly folding*

After dialysis, nanotube samples were concentrated using an Amicon® Ultra centrifugal filter (MWCO 10 kDa) and the sample was incubated at 4 °C overnight to allow assembly. Nanotube assembly exhibits a critical concentration of 0.1 mg/mL; below this threshold, oligomeric species predominate.

For the array samples, after dialysis, the solutions were incubated at 4 °C for a minimum of 48 hours to allow folding. During this period, phase separation occurred, and arrays were enriched in the insoluble fraction. The critical concentration for array formation was approximately 0.5 mg/mL.

##### *Mass Photometry*

Mass photometry results were collected using a Refeyn Two MP Auto Mass photometer (Refeyn). The instrument was first calibrated with a Beta amylase standard (56, 112 and 224 kDa). Designed proteins were diluted to be ~20 nM into SEC buffers. 20 µL of protein solutions were loaded into each well of a clean cassette. Samples were scanned with AquireMP software (Refeyn) with a 3000 frame video, and analyzed to fit the molecular weight distributions to multiple Gaussians with DiscoverMP software (Refeyn).

##### *Mass Spectrometry*

To identify the molecular mass of each protein, intact mass spectra was obtained via reverse-phase Liquid Chromatography-Mass Spectrometry (LC-MS) on an Agilent G6230B TOF on an AdvanceBio RP-Desalting column, and subsequently deconvoluted by way of Bioconfirm using a total entropy algorithm.

##### *Attenuated Total Reflectance Fourier Transform Infrared spectroscopy*

ATR-FTIR experiments were performed with a Thermo Scientific Nicolet iS50R Research FTIR Spectrometer instrument. ATR mode was used with a scanning range of 700-1900 $cm^{-1}$. Each sample was collected with 32 scans at 4 $cm^{-1}$ resolution using SEC buffer as the blank.

##### *Circular Dichroism Measurements*

CD experiments were performed with the JASCO-1500 Spectrophotometer equipped with a temperature-controlled multi-cell holder. In detail, 250 µL of protein samples of ~ 0.2 mg/mL were added into the cuvettes with a pathlength of 1 mm. Spectrum-wavelength scan was recorded from 190-260 nm with 1 nm increment. Temperature-wavelength scan experiments were conducted from 25 °C to 75 °C with a heating rate of 1 °C/minute. The molar residue ellipticity (MRE) value was calculated with the equation, MRE[deg cm^2 dmol^-1] = CDsignal [mdeg] / ( pathlength[mm] * concentration[M] * number of amino acids).

***Protein Negative Staining***
Proteins were diluted to 0.01–0.05 mg/mL in the SEC buffer. A 3 µL aliquot of each sample was applied to freshly glow-discharged carbon-coated 400-mesh copper grids (Electron Microscopy Sciences). Samples were wicked away with filter papers after 30 seconds of adsorption. 3 µL of 2 % uranyl formate was added onto the grids and wicked away after 30 seconds. The staining process was repeated 2 more times. Grids were left air dry before imaging.

Screening nsEM images of 2D protein array design have been deposited to Zenodo(*50*) as the volume of images precludes inclusion in the SI.

***Transmission Electron Microscopy***
TEM and nsEM data were collected on a FEI Talos L120C TEM, 120 kV (FEI Thermo Scientific, Hillsboro, OR) equipped with a Ceta 4K CCD camera. Micrographs collection was automated using the EPU software (FEI Thermo Scientific).

***High Resolution Transmission Electron Microscopy***
HRTEM imaging of the mineralized sample was performed in conventional TEM mode with aberration-corrected FEI Titan ETEM G2 at a voltage of 300 kV. Images were recorded with Gatan's Metro 300 camera, which enables low-dose and dose-rate imaging.

***Scanning Transmission Electron Microscopy – Energy-Dispersive X-ray Spectroscopy***
STEM/EDS characterization of mineralized samples was performed using an aberration-corrected Thermo Fisher Scientific Themis Z scanning/transmission electron microscope (S/TEM) operated at 300 kV. Samples were imaged in scanning transmission electron microscopy mode using high-angle annular dark-field (HAADF) imaging, with a probe convergence semi-angle of 25 mrad. To minimize beam-induced damage, HAADF images were acquired under dose-minimized conditions using multi-frame integration. Image acquisition and basic processing were performed using Thermo Fisher Velox software. Elemental mapping was performed by energy-dispersive X-ray spectroscopy (EDS) using a Thermo Fisher Super-X silicon drift detector (SDD) system.

***nsEM results analysis with CryoSPARC***
For negative-stain electron microscopy (nsEM) data processing, micrographs were imported into CryoSPARC, followed by patch contrast transfer function (CTF) estimation and blob picking using a specified particle diameter range. The upper and lower bounds were set to 10 Å above and below the designed oligomer diameter, respectively. After initial particle picking, micrographs were inspected, particles were extracted, and 2D classification was performed without CTF correction. Selected 2D classes were then used as templates for a second round of template-based particle picking, followed by inspection, extraction, and another round of 2D classification. The classes from this step were selected as the final particle set and used for ab initio reconstruction. Ab initio models were generated with and without symmetry enforcement to evaluate consistency of the resulting oligomer structures.

For data processing of the nanotubes, instead of blob picker/template picker, we did filament tracer. 2D classification was performed similarly. Instead of *ab initio*, we did helical refinement for reconstruction.

***Cryogenic Electron Microscopy Sample Preparation***

For the mineralized HAp1_$C_3$ oligomer, 3 µL of ~0.2 mg/mL protein in SEC buffer was applied to a glow-discharged Quantifoil 2 nm thin carbon R 2.0/2.0 300 mesh grid (Q3100CR2-2nm, EMS). After 30 seconds of adsorption, the excessive protein solution was wicked away with filter paper. The grid was air dry and then flipped upside down on top of the precursor solution in a 1.5 mL eppendorf tube (Sigma Aldrich) containing 800 µL of 2.5 mM $CaCl_2$, 1.0 mM $K_2HPO_4$. The reaction was kept at 37 ºC for 24 hours. After reaction, the grid was picked and subsequently rinsed three times with fresh MQ water to remove excessive precursors. The grid was transferred into a Vitrobot Mark IV (Thermo Fisher Scientific), and was blotted and plunge-frozen into liquid ethane with a blotting time of 0.5 seconds, blotting force 0 and wait time 2 seconds at 22 °C and 100% humidity.

For nanotubes, 3 µL of ~2 mg/mL protein in the SEC buffer was applied to a glow-discharged Au Holey Quantifoil R1.2/1.3 300 mesh grid (Q3100AR1.3, EMS). The grid was blotted and plunge-frozen into liquid ethane using a Vitrobot Mark IV (Thermo Fisher Scientific) at 22 °C and 100% humidity with blotting time of 6 seconds, blotting force 0, and wait time 7 seconds.

For mineralized HAp1_Nt, 3 µL of reaction solution (assay 2) was applied to a glow-discharged Quantifoil 2 nm thin carbon R 2.0/2.0 300 mesh grid (Q3100CR2-2nm, EMS). The grid was blotted and plunge-frozen into liquid ethane using a Vitrobot Mark IV (Thermo Fisher Scientific) at 22 °C and 100% humidity with blotting time of 0.5 seconds, blotting force 0, and wait time 7 seconds.

***Cryogenic Electron Microscopy***

Data of the mineralized HAp1_$C_3$ oligomer were collected automatically using SerialEM(*52*) with a Thermo Fisher Glacios transmission electron microscope 200 kV equipped with a K3 direct electron detector (Gatan). A total dose of 50 $e^-/Å^2$ was applied over a 5.009-s exposure, fractionated into 99 frames with a frame time of 0.0505 s. Data were collected with random defocus ranges spanning between −0.8 and −1.8 µm in six equal steps using beam-image shift three shots per hole and nine holes per stage.

CryoEM data for the HAp1_Nt nanotube were collected at the Janelia Research Campus on a 300 kV transmission electron microscope equipped with a spherical aberration corrector (Cs = 2.7 mm) and an energy filter operated with a 20 eV slit width. Data were acquired at a nominal magnification of 105,000×, corresponding to a calibrated physical pixel size of 1.061 Å. Movies were recorded in super-resolution mode with a calibrated super-resolution pixel size of 0.5305 Å. Each movie was collected with a total electron exposure of 50 $e^-/Å^2$ fractionated over 75 frames, corresponding to a dose of approximately 0.67 $e^-/Å^2$ per frame. Data were collected using a three-shots-per-hole acquisition strategy with a target defocus range of −1.0 to −1.8 µm. An objective aperture of 70 µm was used during data acquisition.

***CryoEM data processing with CryoSPARC***

4030 raw movies of HAp1_Nt were imported into CryoSPARC and were subjected to patch motion correction to correct for beam-induced motion. Subsequently, patch CTF estimation was performed to determine the CTF parameters for each micrograph. Micrographs were curated

using Curate Exposure job with a threshold of maximum CTF fit resolution of 4 Å and maximum total full-frame motion distance of 78.42 pixels. Following curation, 2039 micrographs were accepted and used for particle picking.
The initial particle picking utilized the Filament Tracer tool in CryoSPARC, employing the following parameters: Filament Diameter: 102 Å, Separation Distance: 0.5, Minimum Filament Length: 1. ~790k particles were picked, inspected and extracted with a ~433 Å extraction box size, Fourier-croped by half across 1600 micrographs. Particles were then subjected to 2D class averaging into50 classes. High-quality 2D class averages containing ~85k particles were then selected and used as templates for a second, more refined round of particle picking with the Filament Tracer tool. For the second round of picking, we used parameters of Filament Diameter: 100 Å, Separation Distance: 0.5, Minimum Filament Length: 1. ~457k particles were picked from 1490 micrographs, inspected and extracted with the same ~433 Å extraction box size, and Fourier-cropped by half. The particles were then classified with 2D class averaging and high quality 2D classes containing ~130k particles were selected as new templates for the last round of Filament Tracer. For the last round of picking, we used parameters of Filament Diameter: 92 Å, Separation Distance: 0.5, Minimum Filament Length: 4. ~346k particles were picked from 2039 micrographs, inspected and extracted with ~371 Å extraction box size, Fourier-cropped by half. Particles were classified with 2D class averaging and high quality 2D classes containing ~97k particles were selected. The resulting particle stack was then subjected to *ab initio* 3D reconstruction with helix refinement tool, followed by removing duplicate particles. The resulting map was used for the subsequent homogenous refinement with $D_3$ symmetry applied. The refined map was then continued to non-uniform refinement, yielding a 3.08 Å resolution map. The map was used to build the final three-dimensional structure of the nanotube sample HAp1_Nt. The final cryoEM density maps were deposited in the EMDB under accession number EMD-74450.

##### *CryoEM model building and validation*

An initial atomic model of the HAp1_Nt nanotube was generated by rigid-body docking of the computational design model into the final sharpened cryoEM density map using UCSF ChimeraX. The docked model was then subjected to automated real-space refinement using NAMDINATOR(*53*), which performs short molecular dynamics simulations under real-space restraints to improve model-to-map agreement while preserving stereochemistry. Following NAMDINATOR refinement, a single asymmetric unit chain was manually edited in Coot(*54, 55*). Manual adjustments focused on optimizing backbone placement, correcting side-chain rotamers, and improving local agreement with the density. This chain was further refined using ISOLDE(*56*) within UCSF ChimeraX(*57*) to resolve Ramachandran outliers, reduce local strain, and improve overall model quality under interactive molecular dynamics-based restraints. The refined chain was then symmetrized in UCSF ChimeraX to generate the full nanotube assembly. The symmetrized model was subsequently subjected to additional refinement using ISOLDE to resolve symmetry-related clashes, improve inter-chain interfaces, and ensure consistent geometry across the full assembly while maintaining agreement with the experimental density. A final round of Phenix real-space refinement(*58*) was performed on the symmetrized model, allowing for global minimization and optimization of geometry, including amide side-chain flips where appropriate. Final model quality was assessed using MolProbit(*59*). Figures were generated using UCSF ChimeraX. The final structure was deposited in the Protein Data Bank under accession number 9ZNK.

***Liquid Phase Atomic Force Microscopy***

All LP-AFM experiments were conducted in a pH 7.4 TBS buffer containing 150 mM NaCl using a Bruker MultiMode VIII microscope. Topographic imaging was performed in tapping mode using BL-AC40TS probes ($k \approx 0.1$ N/m; Olympus). The mechanical properties of protein nanotubes before and after mineralization were characterized using PeakForce Tapping mode using ScanAsyst Fluid+ probes ($k \approx 0.7$ N/m, Bruker).

For the protein sample preparation, 10 µL of the protein stock solution was deposited onto a freshly cleaved mica substrate and incubated for 10 min in a humid environment to promote adsorption. Subsequently, 90 µL of the buffer solution was gently added to fully immerse the sample for measurements.

AFM data were processed using Bruker NanoScope Analysis software (v2.0). For the mechanical analysis, force–distance curves were fitted using the Derjaguin–Muller–Toporov (DMT) model to extract the effective elastic modulus. For each sample region, more than 20 individual measurement points were collected to obtain statistically meaningful modulus distributions.

***Mineralization in human dental organoids***

The induced secretory mature ameloblast (ismAM) organoids were generated as previously described(*46*): hiPSCs were differentiated into 2D induced ameloblasts (iAMs) from day 0 to day 16, transferred to low-attachment plates to form 3D induced early ameloblast (ieAM) organoids in Epicult Plus medium (Stemcell Technologies #06070), and matured to the induced secretory-stage maturation ameloblast (ismAM) stage by treatment with the C3-DLL4 Notch activator(*60*). ismAM organoids were treated with 50 µM HAp1_Nt or 100 µM HAp1_Nt_DtoN, a twofold higher concentration used to provide a stringent test of the nucleation-dead control, together with 1 µM calcein (Sigma-Aldrich) and calcium phosphate buffer (2.5 mM Calcium Chloride ($CaCl_2$), 1 mM potassium dibasic phosphate ($K_2HPO_4$) in HEPES buffer, pH 7.4) for 24 hours at 37 °C. Organoids were then maintained for an additional 6 days in media containing calcein and calcium phosphate buffer, with media changes every other day, to assess mineralization and transition to the induced mature ameloblast (imAM) state. Untreated ismAM organoids cultured in calcium phosphate buffer and calcein alone served as controls.

***Confocal imaging***

Following the 7-day mineralization assay, organoids were fixed in 4% paraformaldehyde (PFA), washed 3× for 5 minutes in 1× PBS, and permeabilized in 0.5% Triton X-100 at room temperature for 10 minutes. Organoids were blocked for 2 hours in blocking solution (1× PBS containing 10% BSA, 5% normal goat serum, and 0.1% Triton-X), then incubated overnight at 4 °C with primary antibody against ZO-1 (Invitrogen, #33-9100; RRID: AB_87181), diluted in antibody dilution buffer (1× PBS containing 10% BSA, 5% normal goat serum, and 0.2% Triton-X) at the manufacturer-recommended concentration. The following day, organoids were washed 3× for 6 minutes in 1× PBS, incubated overnight at 4 °C with Mouse IgG secondary antibody conjugated to Alexa Fluor 568 (Thermo Fisher, #A-11004; RRID: AB_2534072, 1:200) and DAPI (Thermo Fisher, #D1306; RRID: AB_2629482, 1:50) in antibody dilution buffer, then washed 3× for 6 minutes in 1× PBS and mounted in Vectashield Antifade Mounting Media on a glass concavity microscope slide, one to three organoids per well. Calcein fluorescence (Sigma-Aldrich, #C0875, 1 μM), retained from the live mineralization assay, was imaged alongside DAPI and ZO-1 without additional processing. Organoids were imaged using a Leica (DMi8) SP8 LIGHTNING confocal microscope (25× and 40× objectives) equipped with HyD and PMT spectral detectors and Leica LAS X acquisition software (version 3.5.5IR).

***Alizarin Red S Staining and quantification***
Fixed organoids were stained with 40 mM Alizarin Red S (Sigma, #A5533; pH 4.1–4.3) for 1 hour at room temperature to label calcium-rich mineralized deposits. Following staining, organoids were washed extensively with deionized water until the washing solution appeared clear, ensuring removal of unbound, non-specific dye. For quantification, stained organoids were transferred to 10% acetic acid and incubated at room temperature until complete dissolution of the organoid matrix, which required approximately 5 days. The resulting acidic extract was neutralized with 10% ammonium hydroxide to restore a pH compatible with accurate spectrophotometric measurement, as Alizarin Red S absorbance is pH-dependent. The absorbance of the neutralized solution was then measured at 405 nm using a microplate reader (BioTek), with values used as a relative measure of mineral deposition across experimental conditions.

***X-ray Diffraction***
5 μL of samples were dropped on a piece of clean silicon wafer (Ted Pella, INC.) The samples were left air dry at room temperature. XRD results were collected on a Bruker D8 Discover instrument.

***Mineralization Assay***
*Assay 1: without pAsp*
1.5-3 μM protein and 2.5 mM $CaCl_2$ (prepared in SEC buffer) were mixed thoroughly in a PCR tube for binding for 5 minutes. Then, 1 mM $K_2HPO_4$ (in SEC buffer) was added and mixed well. The tube was sealed and incubated at 37 °C with shaking at 225 rcf for 24 hours.

*Assay 2: with pAsp*
A precursor solution was prepared by mixing 5 mM $CaCl_2$, 2 mM $K_2HPO_4$, and 360 μg/mL 27 kDa pAsp (Poly(L-aspartic acid sodium salt); Alamanda Polymers) or 60 μg/mL 4.1 kDa pAsp (Poly(L-aspartic acid sodium salt; Alamanda Polymers) in SEC buffer. The mixture was incubated at 37 °C for 3 hours. Then in a PCR tube, 15 μL of protein was mixed with 15 μL of precursor mixture to result in a final concentration of 1.5-3 μM protein, 2.5 mM $CaCl_2$, 1 mM $K_2HPO_4$, and 180 μg/mL pAsp (27kDa) or 30 μg/mL pAsp (4.1 kDa). The PCR tube was sealed and incubated at 37 °C with shaking at 225 rcf for 24 hours. For array mineralization, the precursor solution was added directly to the insoluble pellet of assembled arrays in a PCR tube. This sample was incubated at 4°C without agitation and remained phase-separated throughout the reaction. The insoluble fraction was subsequently characterized by TEM.

**Materials Design Analysis Reporting (MDAR) Checklist**

**Materials:**

| Newly created materials | indicate where provided: page no/section/legend) | n/a |
|---|---|---|
| The manuscript includes a dedicated "materials availability statement" providing transparent disclosure about availability of newly created materials including details on how materials can be accessed and describing any restrictions on access. | p.15, Data, code, and materials availability statement, main text | |

| Antibodies | indicate where provided: page no/section/legend) | n/a |
|---|---|---|
| For commercial reagents, provide supplier name, catalogue number and RRID, if available. | Commercial reagents and genes suppliers are described in the Material and methods section where the reagents appear | |

| DNA and RNA sequences | indicate where provided: page no/section/legend) | n/a |
|---|---|---|
| **Short novel DNA or RNA including primers, probes:** Sequences should be included or deposited in a public repository. | | n/a |

| Cell materials | indicate where provided: page no/section/legend | n/a |
|---|---|---|
| **Cell lines:** Provide species information, strain. Provide accession number in repository **OR** supplier name, catalog number, clone number, **OR** RRID. | Material and methods section in SI | |
| **Primary cultures:** Provide species, strain, sex of origin, genetic modification status. | Material and methods section in SI | |

| Experimental animals | indicate where provided: page no/section/legend) | n/a |
|---|---|---|
| **Laboratory animals or Model organisms:** Provide species, strain, sex, age, genetic modification status. Provide accession number in repository **OR** supplier name, catalog number, clone number, **OR** RRID. | | n/a |

| | | |
|---|---|---|
| **Animal observed in or captured from the field:** Provide species, sex, and age where possible. | | n/a |

| **Plants and microbes** | **indicate where provided: page no/section/legend)** | **n/a** |
|---|---|---|
| **Plants:** provide species and strain, ecotype and cultivar where relevant, unique accession number if available, and source (including location for collected wild specimens). | | n/a |
| **Microbes:** provide species and strain, unique accession number if available, and source. | | n/a |

| **Human research participants** | **indicate where provided: page no/section/legend) or state if these demographics were not collected** | **n/a** |
|---|---|---|
| If collected and within the bounds of privacy constraints report on age, sex and gender or ethnicity for all study participants. | | n/a |

**<u>Design:</u>**

| **Study protocol** | **indicate where provided: page no/section/legend)** | **n/a** |
|---|---|---|
| If study protocol has been pre-registered, provide DOI. For clinical trials, provide the trial registration number **OR** cite DOI. | p.15, Data, code, and materials availability statement, main text | |

| **Laboratory protocol** | **indicate where provided: page no/section/legend)** | **n/a** |
|---|---|---|

| | |
|---|---|
| Provide DOI **OR** other citation details if detailed step-by-step protocols are available. | Detailed step-by-step protocols are described in the Material and Methods section SI. |

| **Experimental study design (statistics details)** | | |
|---|---|---|
| **For in vivo studies:** State whether and how the following have been done | **indicate where provided: page no/section/legend. If it could have been done, but was not, write not done** | **n/a** |
| Sample size determination | | n/a |
| Randomisation | | n/a |
| Blinding | | n/a |
| Inclusion/exclusion criteria | | n/a |

| **Sample definition and in-laboratory replication** | **indicate where provided: page no/section/legend** | **n/a** |
|---|---|---|
| State number of times the experiment was replicated in laboratory. | Protein expression, purification, and mineralization experiments were each performed a minimum of 5 independent times. CryoEM structure determination was performed once from a single optimized sample preparation. | |
| Define whether data describe technical or biological replicates. | All experimental replicates represent biological replicates using independently prepared protein batches. Mineralization experiments, kinetic assays, and ATR-FTIR measurements were each performed on independently prepared protein and mineralization reaction batches (biological replicates, $n \geq 5$). AFM mechanical mapping measurements represent technical replicates across multiple regions of the same sample. CryoEM structure determination and HRTEM imaging (FEI Titan) were each performed once on a single optimized sample preparation. | |

| Ethics | indicate where provided: page no/section/legend | n/a |
|---|---|---|
| **Studies involving human participants:** State details of authority granting ethics approval (IRB or equivalent committee(s), provide reference number for approval. | | n/a |
| **Studies involving experimental animals:** State details of authority granting ethics approval (IRB or equivalent committee(s), provide reference number for approval. | | n/a |
| **Studies involving specimen and field samples:** State if relevant permits obtained, provide details of authority approving study; if none were required, explain why. | | n/a |

| Dual Use Research of Concern (DURC) | indicate where provided: page no/section/legend | n/a |
|---|---|---|
| If study is subject to dual use research of concern regulations, state the authority granting approval and reference number for the regulatory approval. | | n/a |

**Analysis:**

| Attrition | indicate where provided: page no/section/legend | n/a |
|---|---|---|
| Describe whether exclusion criteria were preestablished. Report if sample or data points were omitted from analysis. If yes report if this was due to attrition or intentional exclusion and provide justification. | No exclusion criteria were preestablished and no experimental batches or data points were omitted from analysis. CryoEM particle selection and classification followed standard processing procedures; particle numbers at each stage are detailed in Table S3 (cryoEM data collection and refinement statistics) | |

| Statistics | indicate where provided: page no/section/legend | n/a |
|---|---|---|

| | |
|---|---|
| Describe statistical tests used and justify choice of tests. | Statistical analysis was performed for AFM mechanical mapping data. Stiffness/modulus values were compared between mineralized (n=22) and unmineralized control (n=22) groups using a two-tailed Student's t-test |

| **Data availability** | **indicate where provided: page no/section/legend** | **n/a** |
|---|---|---|
| For newly created and reused datasets, the manuscript includes a data availability statement that provides details for access or notes restrictions on access. | p.15, Data, code, and materials availability statement, main text | |
| If newly created datasets are publicly available, provide accession number in repository **OR** DOI **OR** URL and licensing details where available. | p.15, Data, code, and materials availability statement, main text | |
| If reused data is publicly available provide accession number in repository **OR** DOI **OR** URL, **OR** citation. | p.15, Data, code, and materials availability statement, main text | |

| **Code availability** | **indicate where provided: page no/section/legend** | **n/a** |
|---|---|---|
| For all newly generated custom computer code/software/mathematical algorithm or re-used code essential for replicating the main findings of the study, the manuscript includes a data availability statement that provides details for access or notes restrictions. | p.15, Data, code, and materials availability statement, main text | |
| If newly generated code is publicly available, provide accession number in repository, **OR** DOI **OR** URL and licensing details where available. State any restrictions on code availability or accessibility. | p.15, Data, code, and materials availability statement, main text | |
| If reused code is publicly available provide accession number in repository **OR** DOI **OR** URL, **OR** citation. | p.15, Data, code, and materials availability statement, main text | |

#### Reporting

MDAR framework recommends adoption of discipline-specific guidelines, established and endorsed through community initiatives. Journals have their own policy about requiring specific guidelines and recommendations to complement MDAR.

| Adherence to community standards | indicate where provided: page no/section/legend | n/a |
|---|---|---|
| State if relevant guidelines (e.g., ICMJE, MIBBI, ARRIVE) have been followed, and whether a checklist (e.g., CONSORT, PRISMA, ARRIVE) is provided with the manuscript. | | n/a |

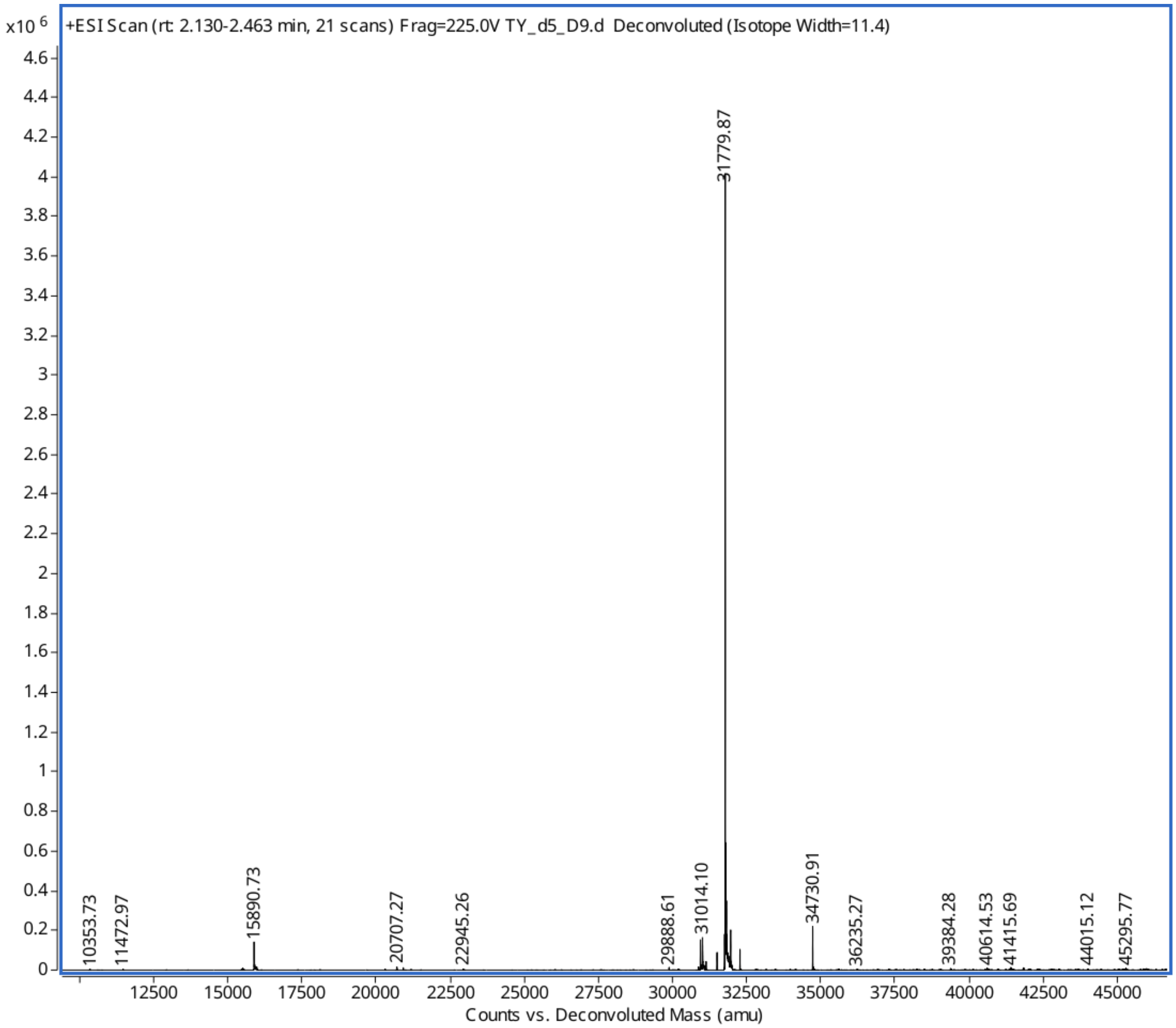

**Fig. S1 Mass Spectrometry Characterization of Designed Proteins.** HAp1_$C_3$

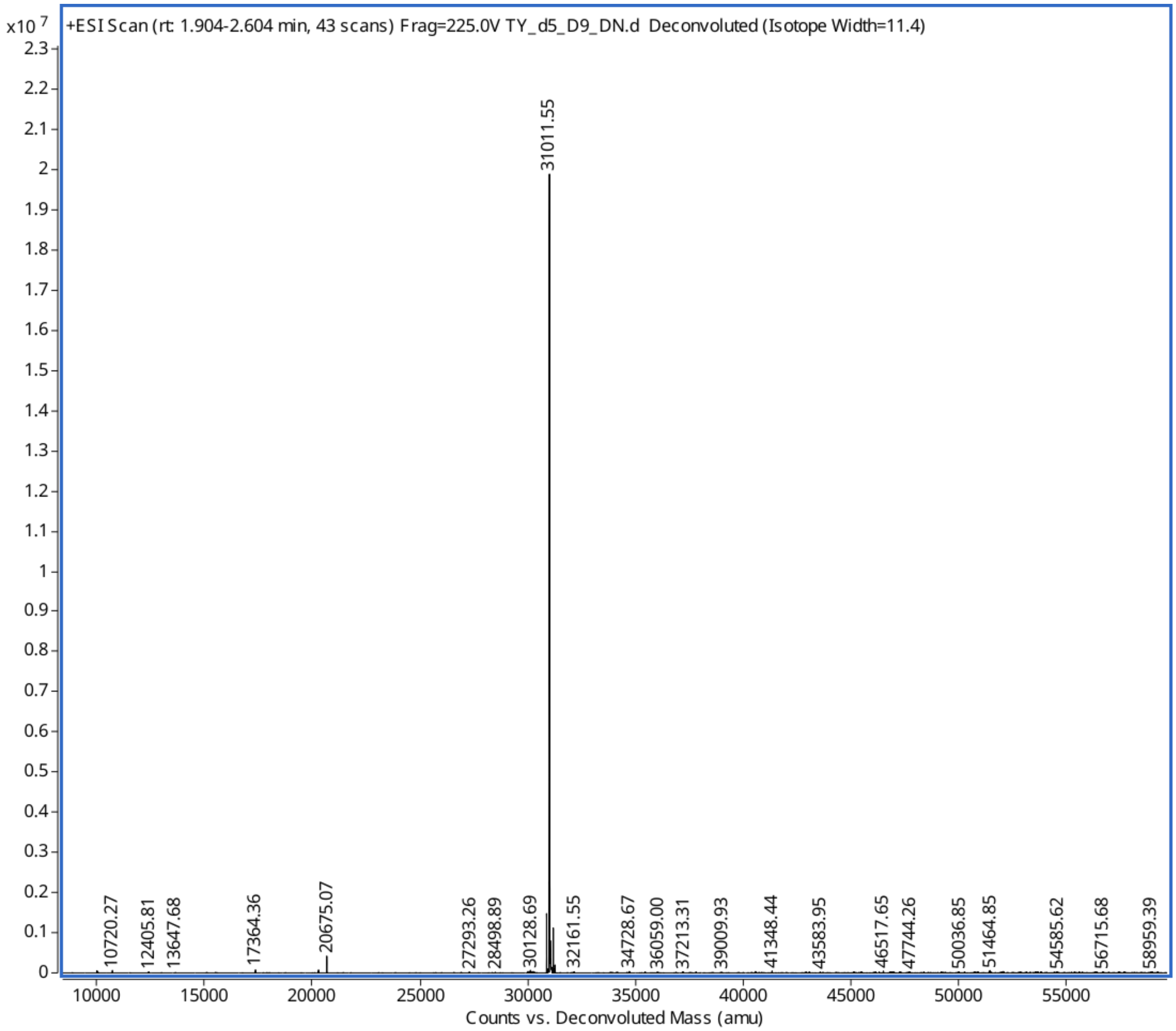

**Fig. S2 Mass Spectrometry Characterization of Designed Proteins.** HAp1_$C_3$_DtoN

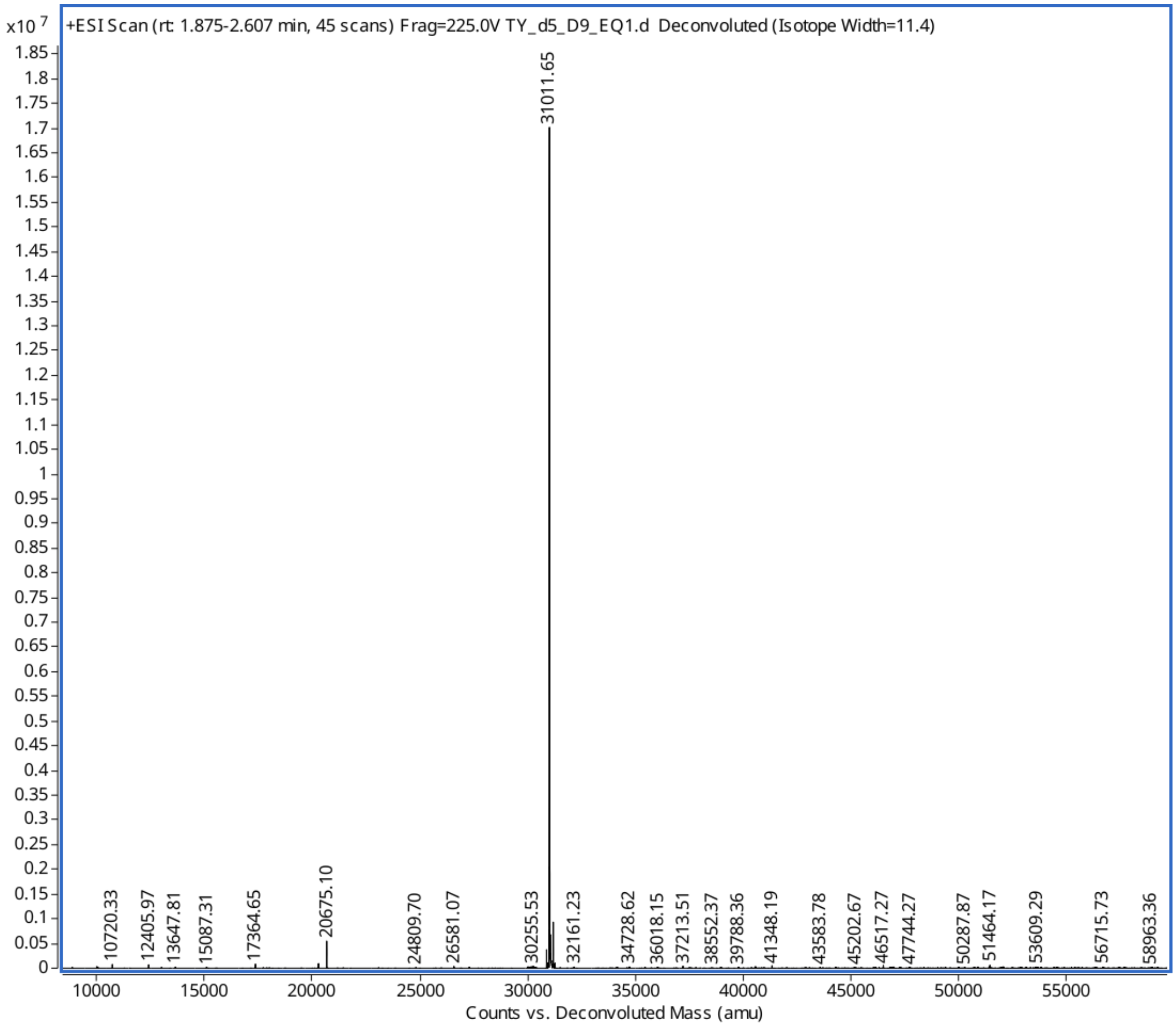

**Fig. S3 Mass Spectrometry Characterization of Designed Proteins.** HAp1_$C_3$_EtoQ1

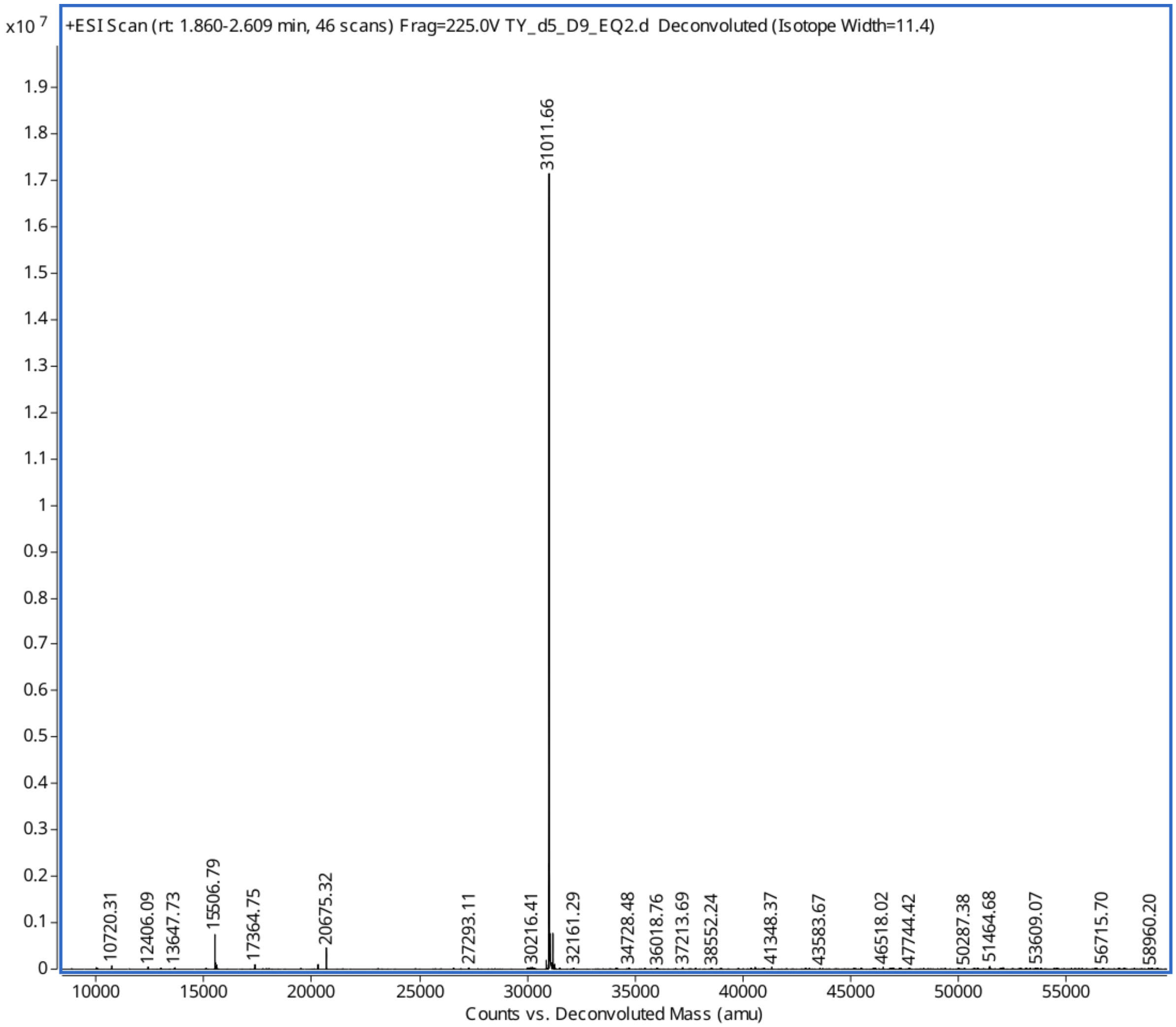

**Fig. S4 Mass Spectrometry Characterization of Designed Proteins.** HAp1_$C_3$_EtoQ2

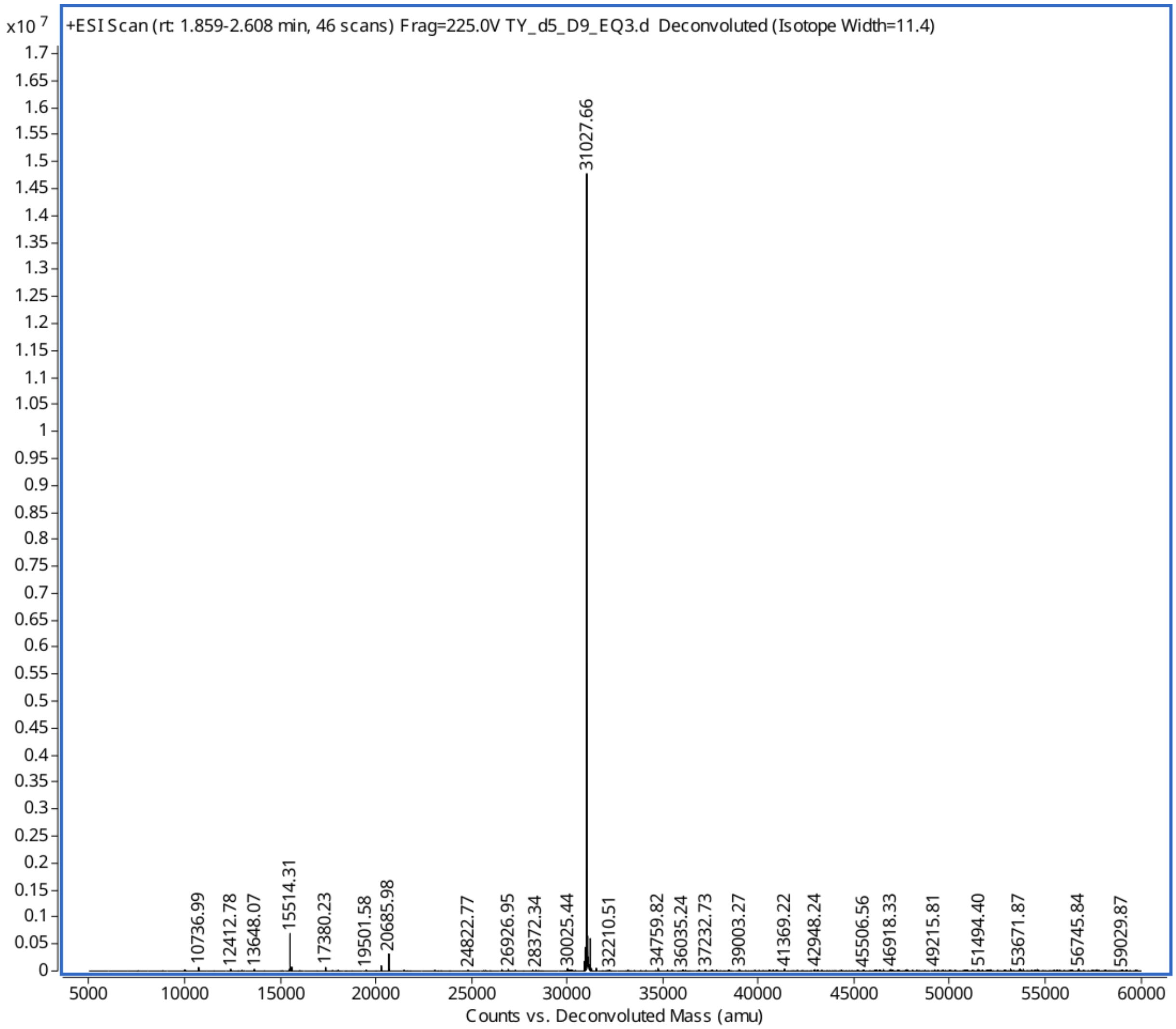

**Fig. S5 Mass Spectrometry Characterization of Designed Proteins.** HAp1_$C_3$_EtoQ3

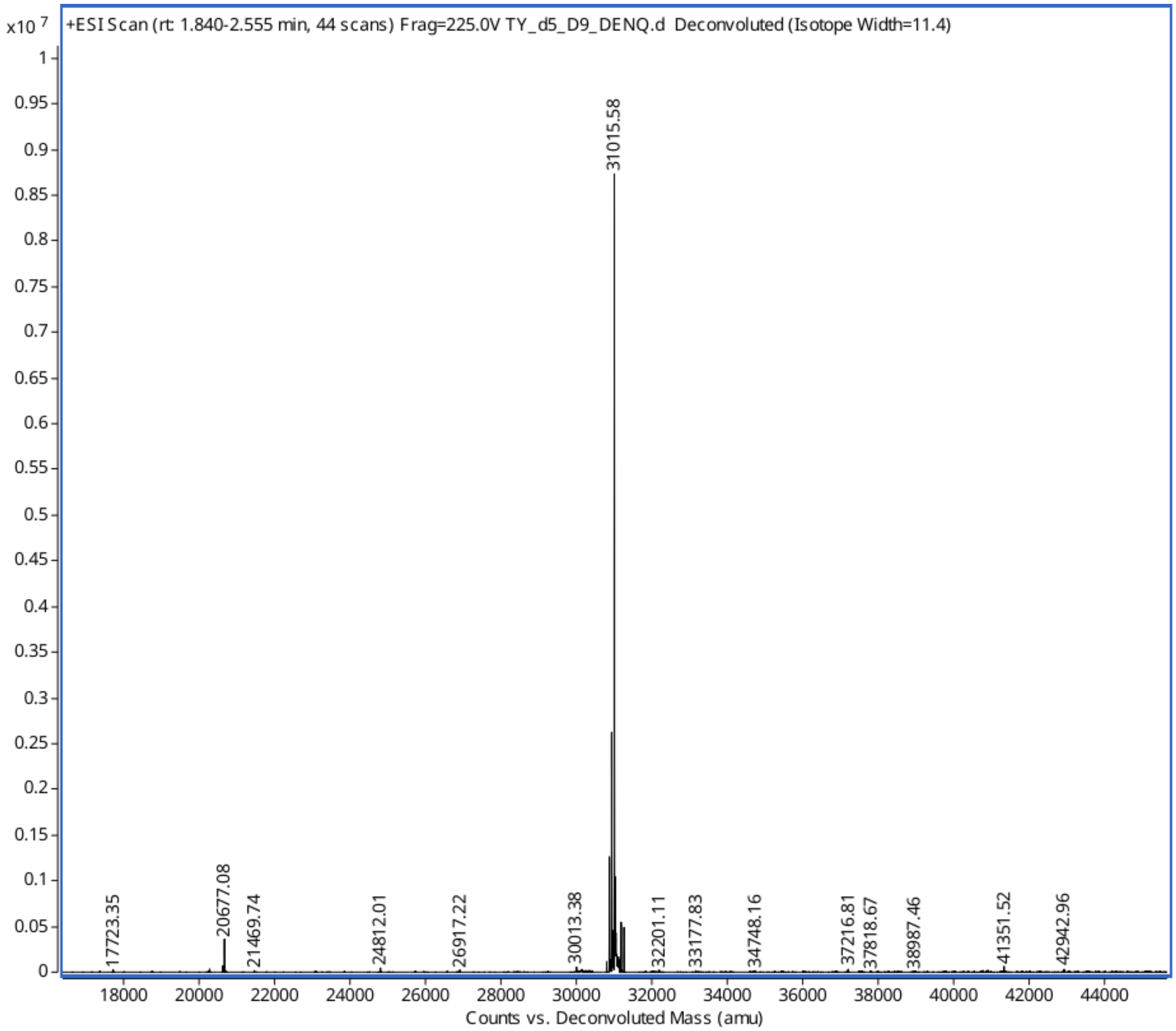

**Fig. S6 Mass Spectrometry Characterization of Designed Proteins.** HAp1_$C_3$_DENQ

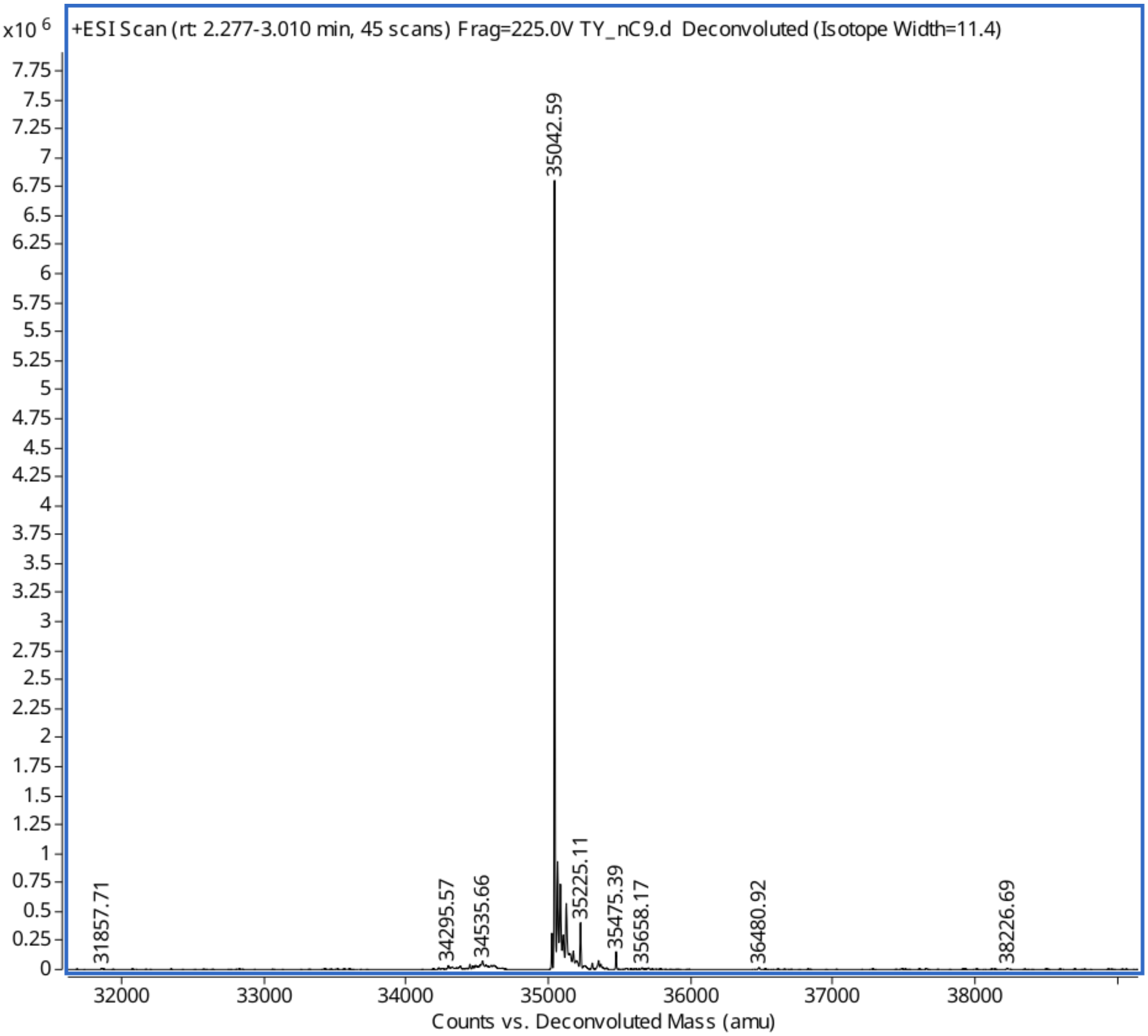

**Fig. S7 Mass Spectrometry Characterization of Designed Proteins.** HAp1_D3N

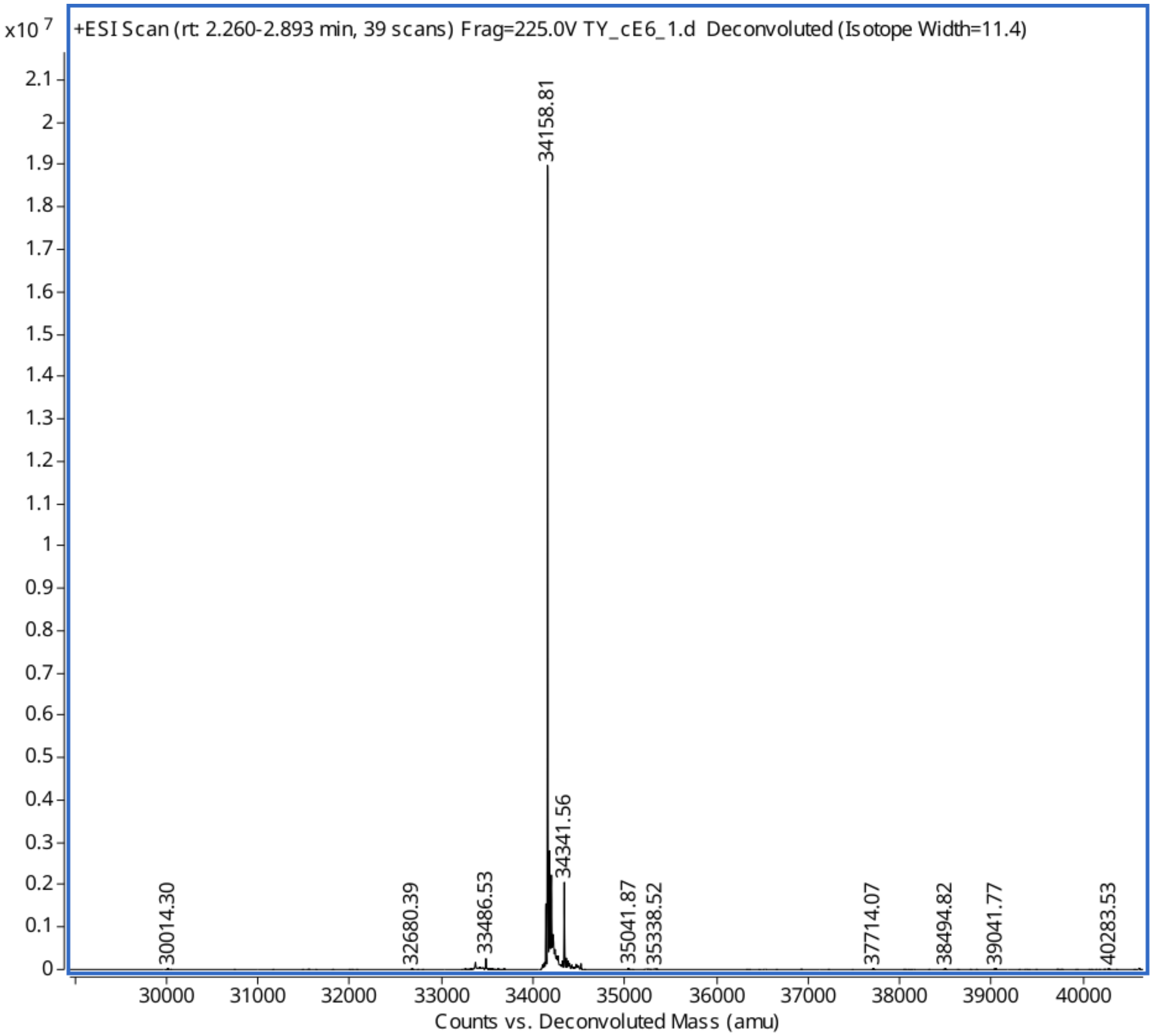

**Fig. S8 Mass Spectrometry Characterization of Designed Proteins.** HAp1_D$_3$C

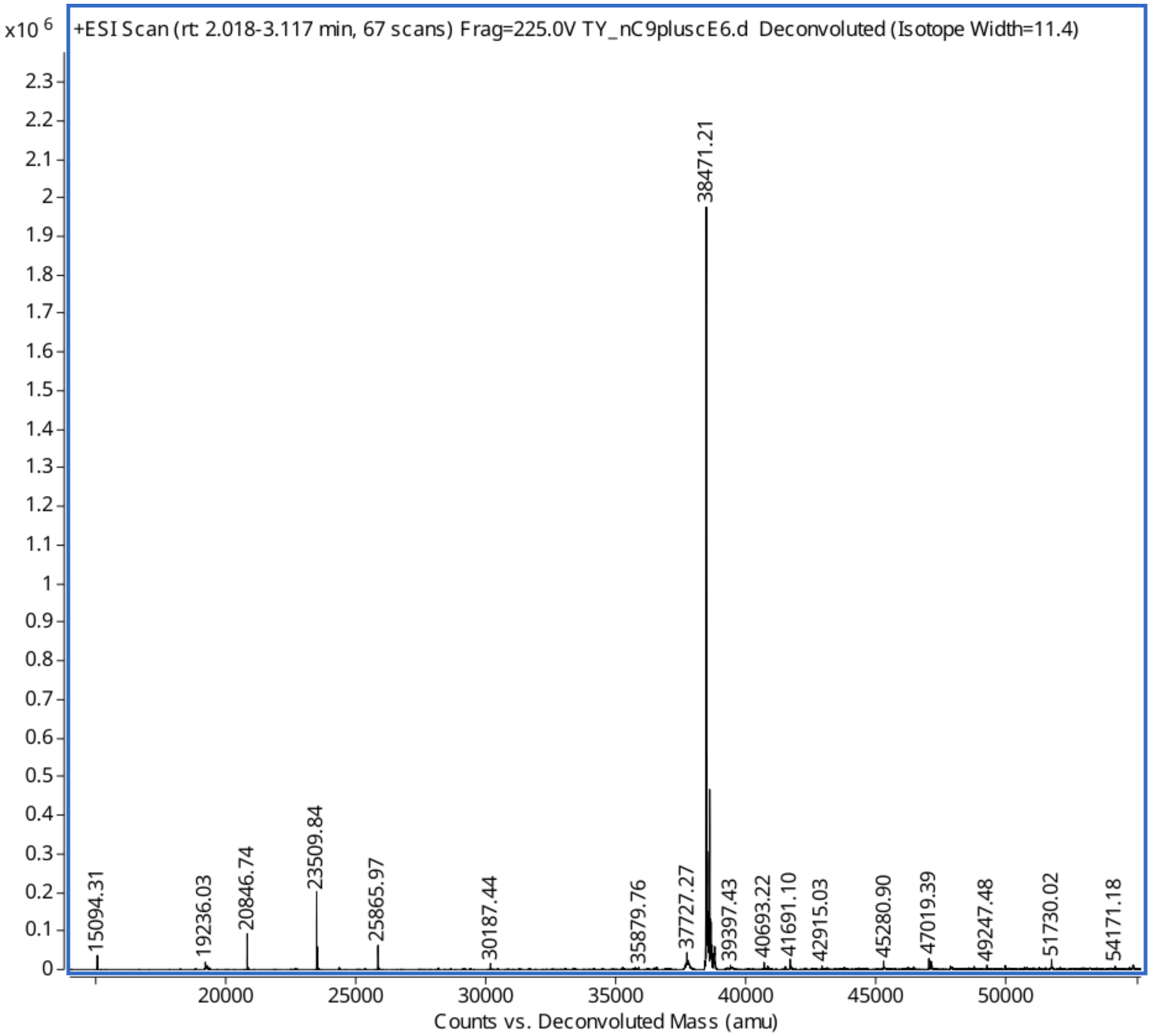

**Fig. S9 Mass Spectrometry Characterization of Designed Proteins.** HAp1_Nt.

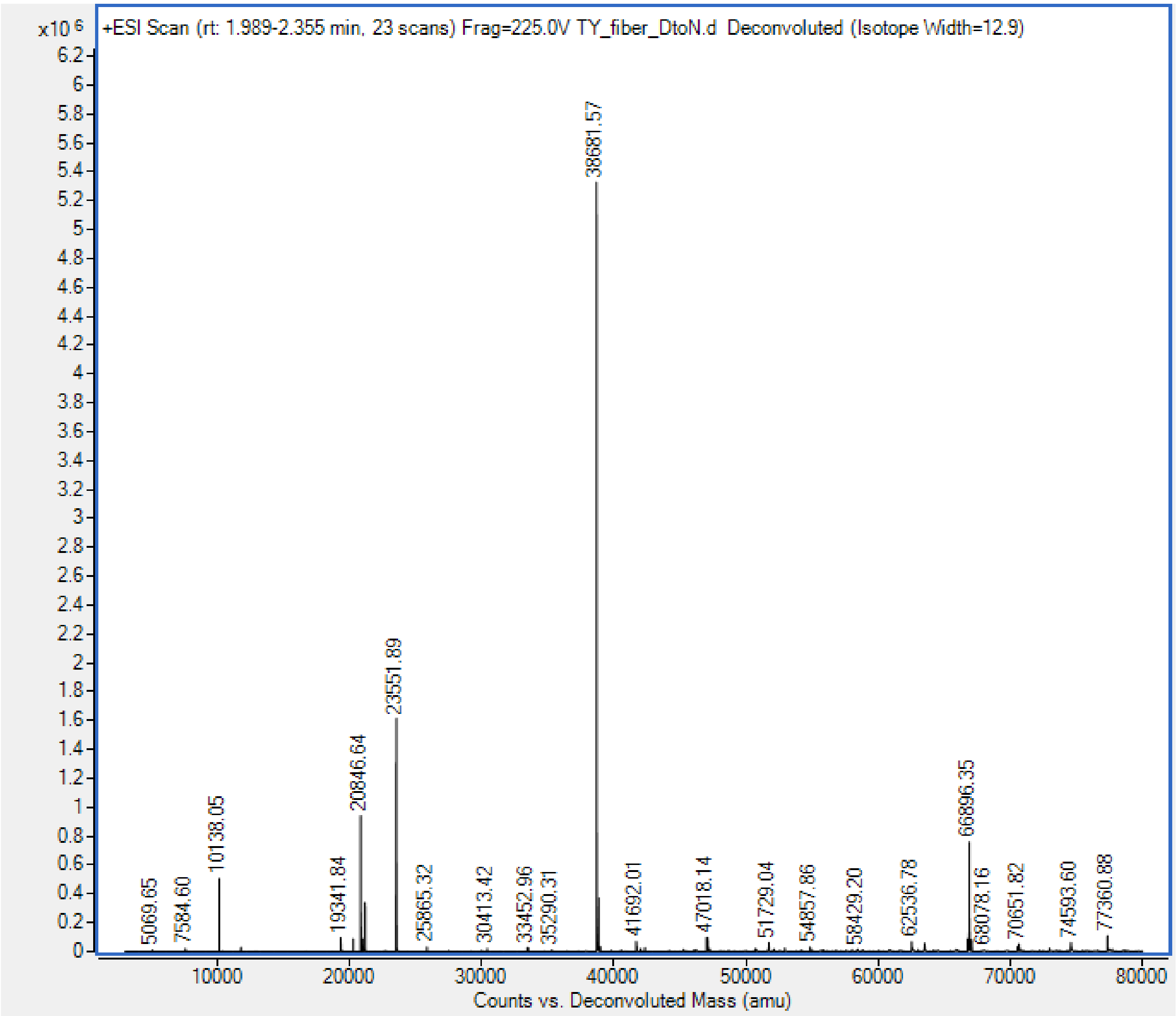

**Fig. S10 Mass Spectrometry Characterization of Designed Proteins.** HAp1_Nt_DtoN

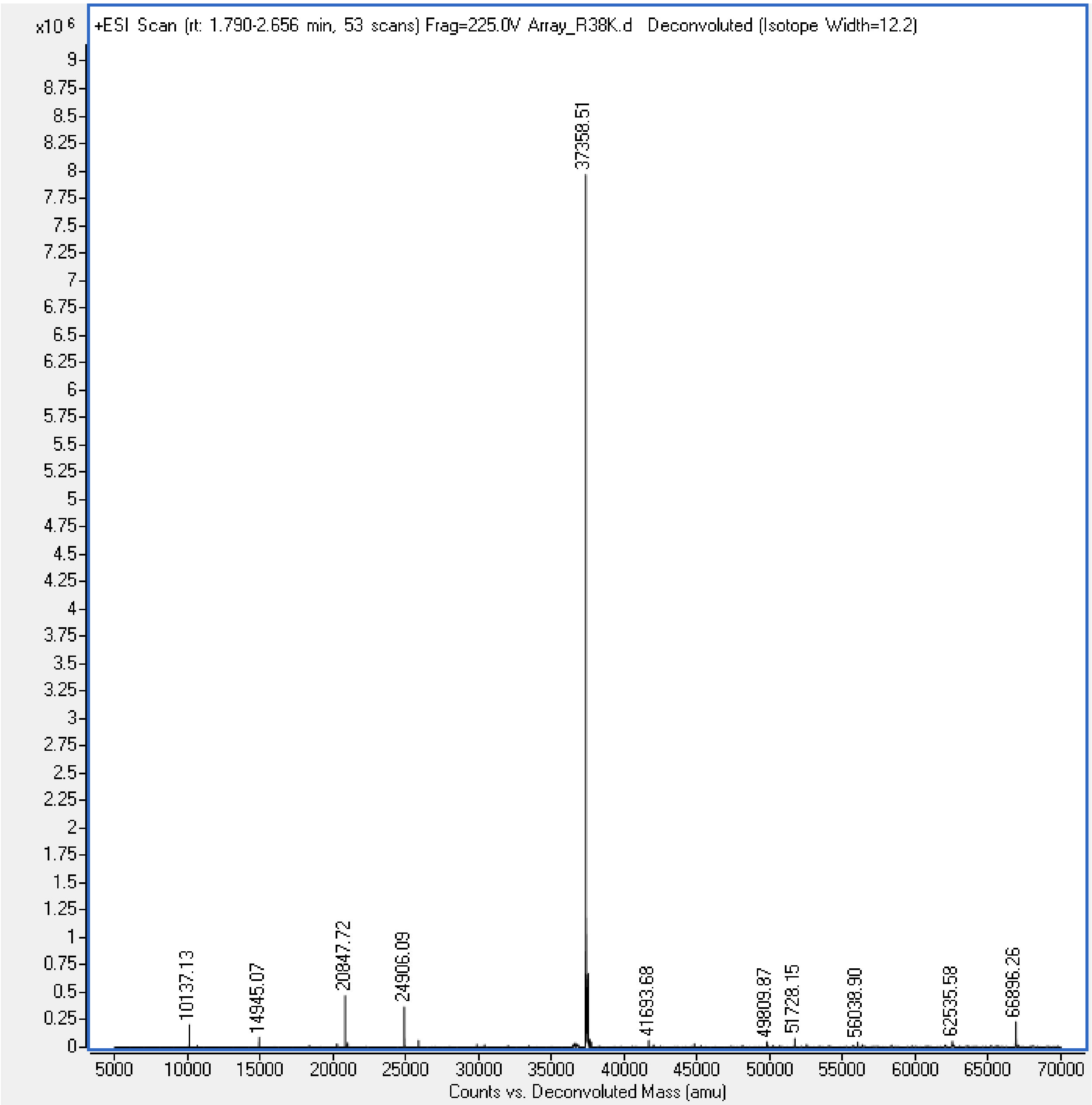

**Fig. S11 Mass Spectrometry Characterization of Designed Proteins.** HAp1_Ar.

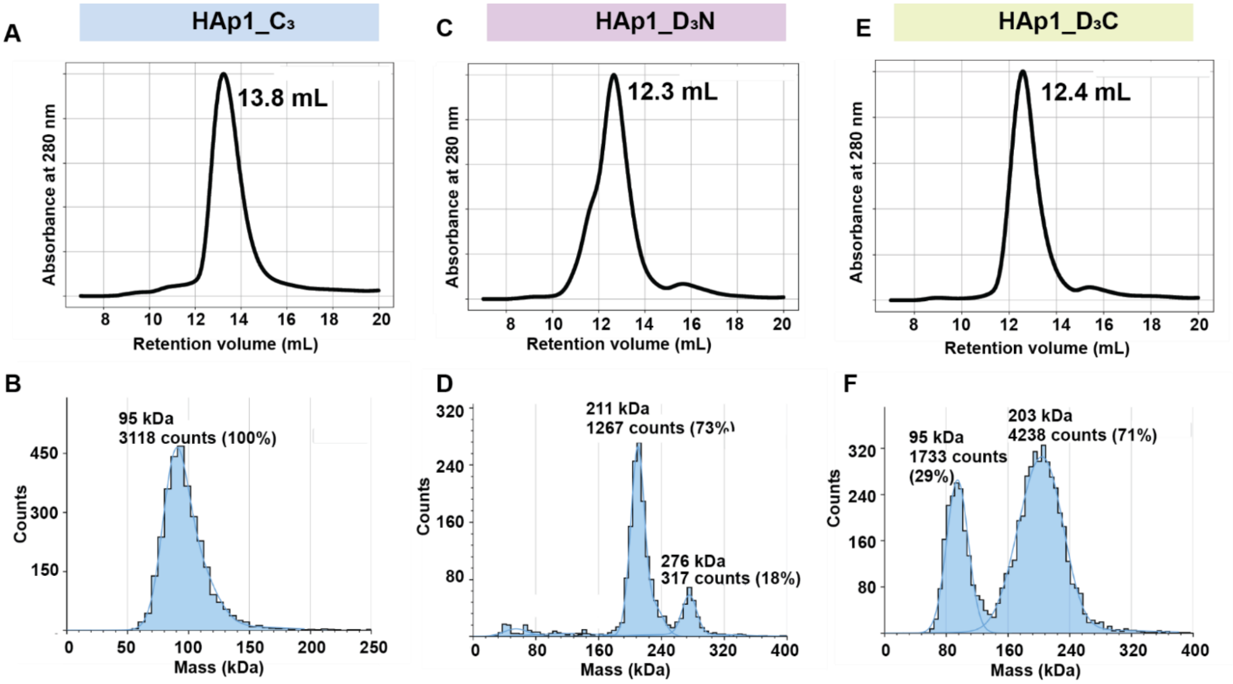

**Fig. S12 SEC and MP characterization of oligomers.** (**A**) and (**B**) SEC and MP profiles of HAp1_C3 oligomer, indicating a trimeric species. (**C**) and (**D**) SEC and MP profiles of HAp1_D3N oligomer, indicating predominantly hexameric species with minor higher-order oligomers. (**E**) and (**F**) SEC and MP profiles of HAp1_D3C oligomer, indicating predominantly hexameric species with a minor trimeric population.

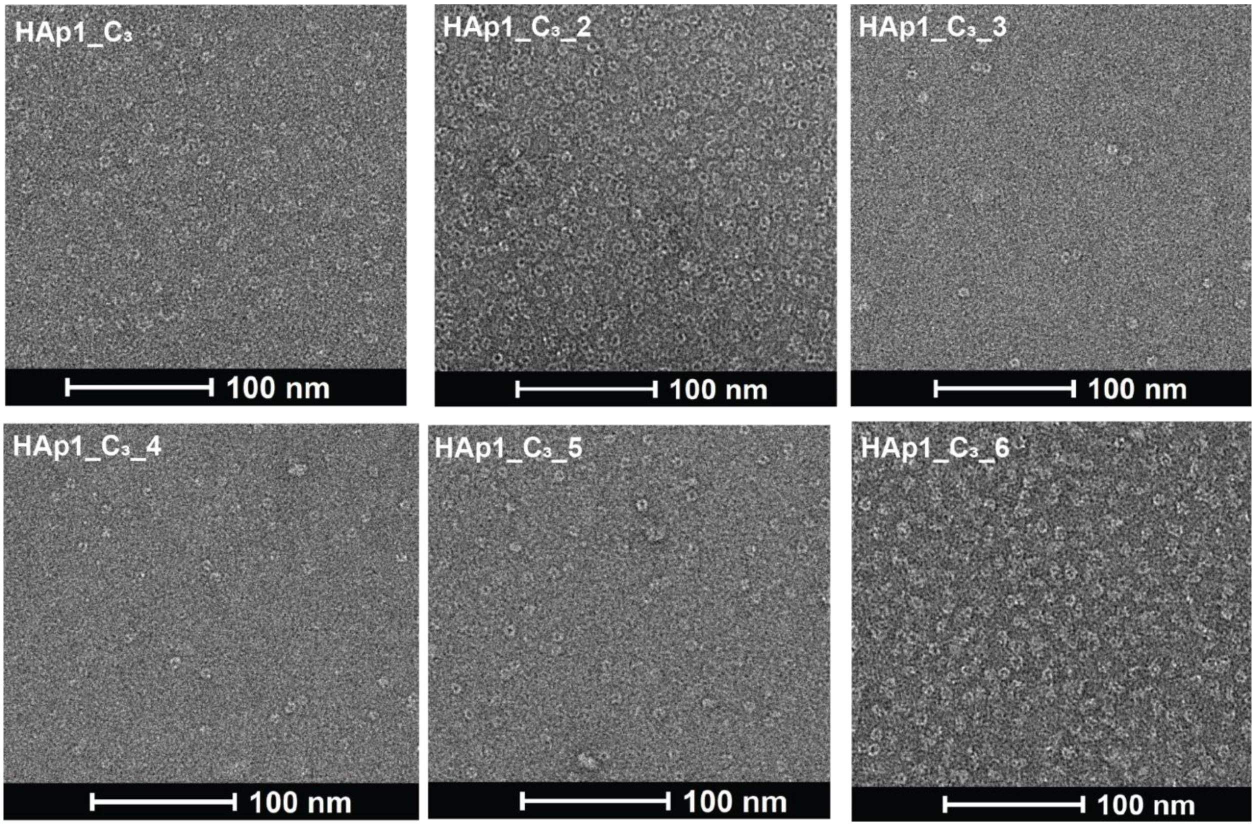

**Fig. S13 nsEM characterization of HAp1_C3 oligomers.**

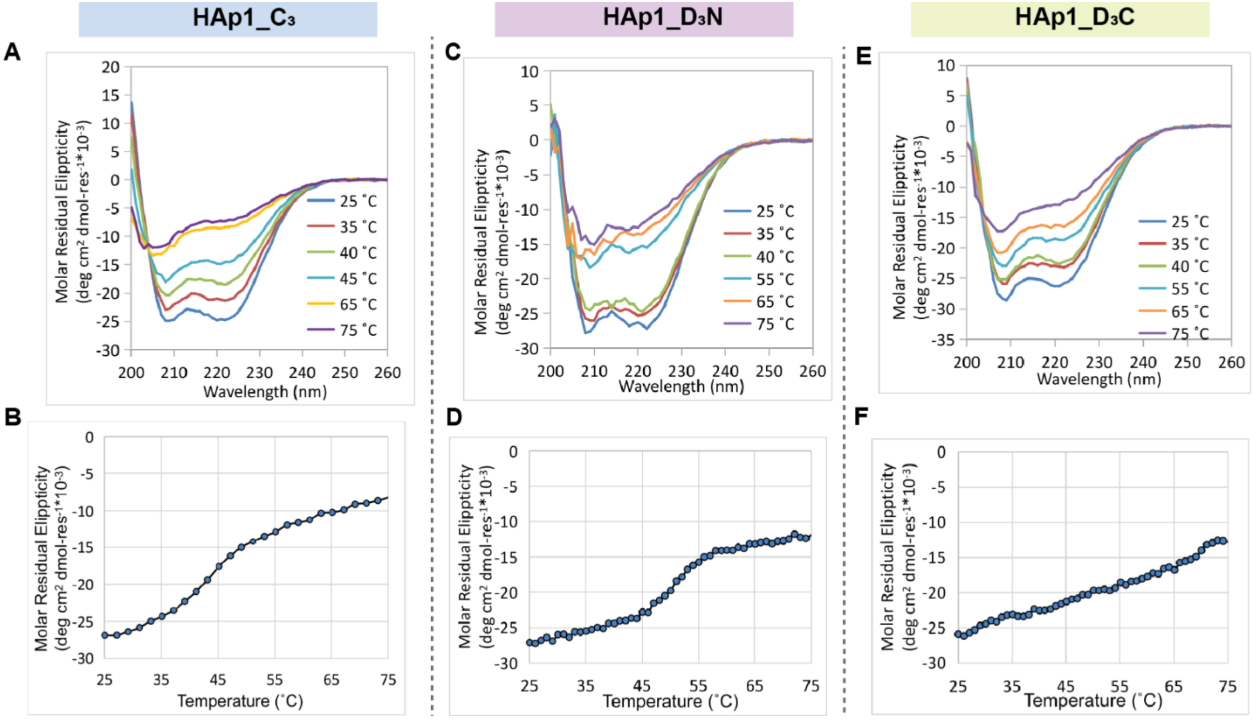

**Fig. S14 CD characterization of oligomers.** (**A-E**) spectra scan of oligomers HAp1_C3, HAp1_D3N and HAp1_D3C at different temperatures. (**B-F**) melting curves of oligomers HAp1_C3, HAp1_D3N and HAp1_D3C. Signals were monitored at 222 nm wavelength.

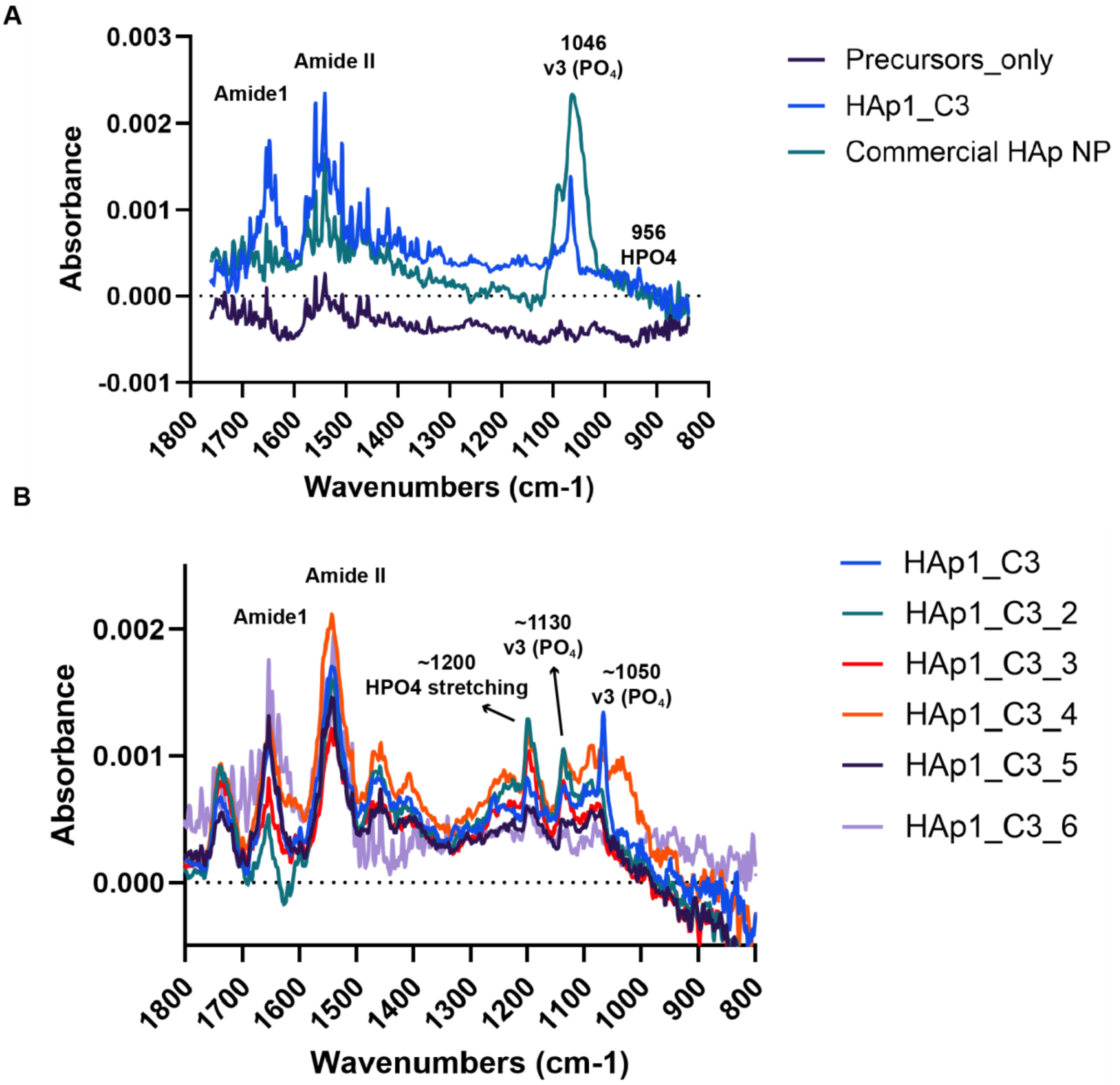

**Fig. S15 ATR-FTIR characterization of HAp1_C**$_3$ oligomers **mineralization** with assay 1**.** (**A**) ATR-FTIR comparison of reactions containing HAp1_C$_3$, no protein and commercial HAp nanoparticles. (**B**) ATR-FTIR characterization of reactions containing folded HAp1_C$_3$ oligomers.

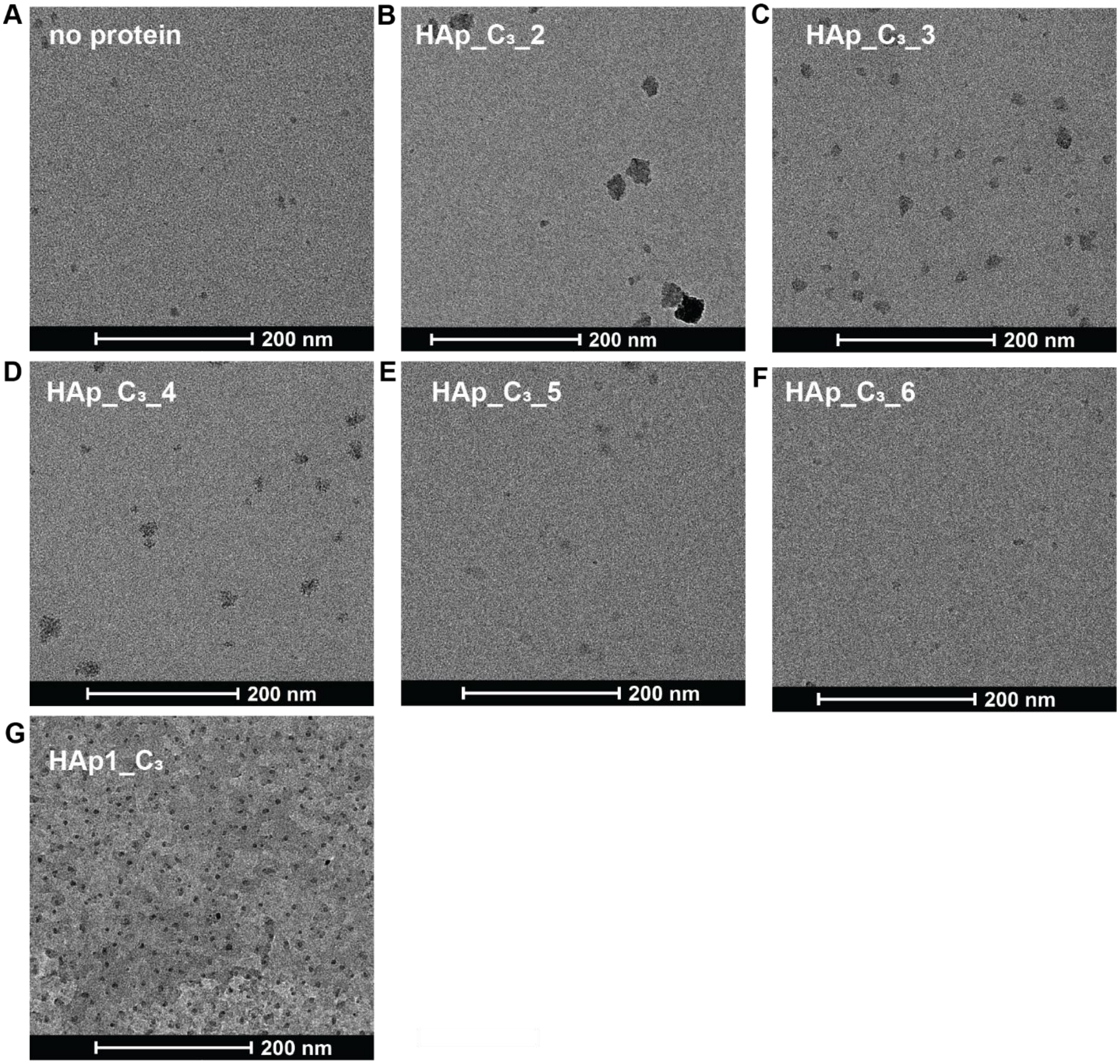

**Fig. S16 TEM characterization of HAp1_$C_3$ oligomers mineralized with assay 1.** (**A**) no protein, 2.5 mM $CaCl_2$ and 1 mM $K_2HPO_4$, (**B**) HAp1_$C_3$_2, (**C**) HAp1_$C_3$_3, (**D**) HAp1_$C_3$_4 (**E**) HAp1_$C_3$_4, (**F**) HAp1_$C_3$_6 (**G**) HAp1_$C_3$.

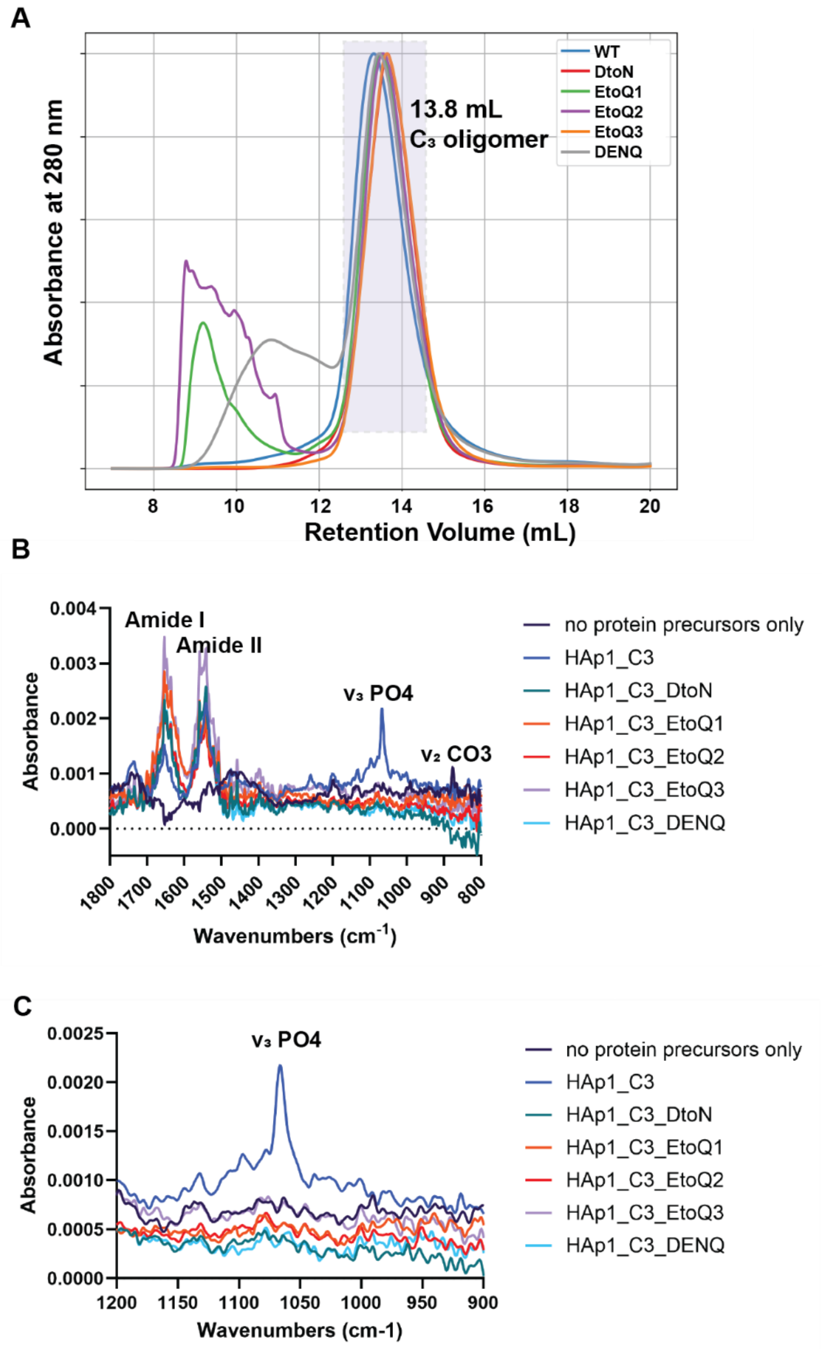

**Fig. S17 HAp1_$C_3$ knockout mutants characterization.** (**A**) SEC profiles of knockout mutants of HAp1_$C_3$. The species eluted at 9-12 minute retention volume are higher-order species. Species eluted at 13.8 mL retention volume were collected and expected to be trimers. (**B**) and (**C**) ATR-FTIR characterization of mineralized HAp1_$C_3$ oligomer and knockout mutants.

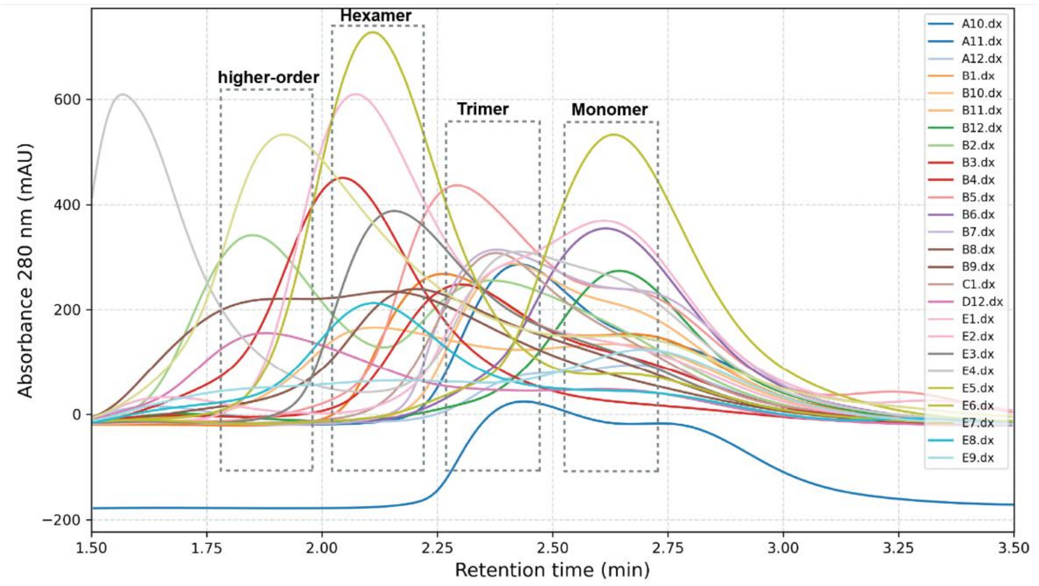

**Fig. S18 SEC traces of HAp_$D_3$ oligomers packed against their C-termini using Superdex 200 Increase 5/150 GL column.** The design HAp1_$D_3$C that's presented in the main text was named as E6 in this plot. Among these 26 designs, one formed aggregate, five formed higher-order species, four formed majorly hexamer, seven formed majorly trimers, five formed monomers, and four are ambiguous.

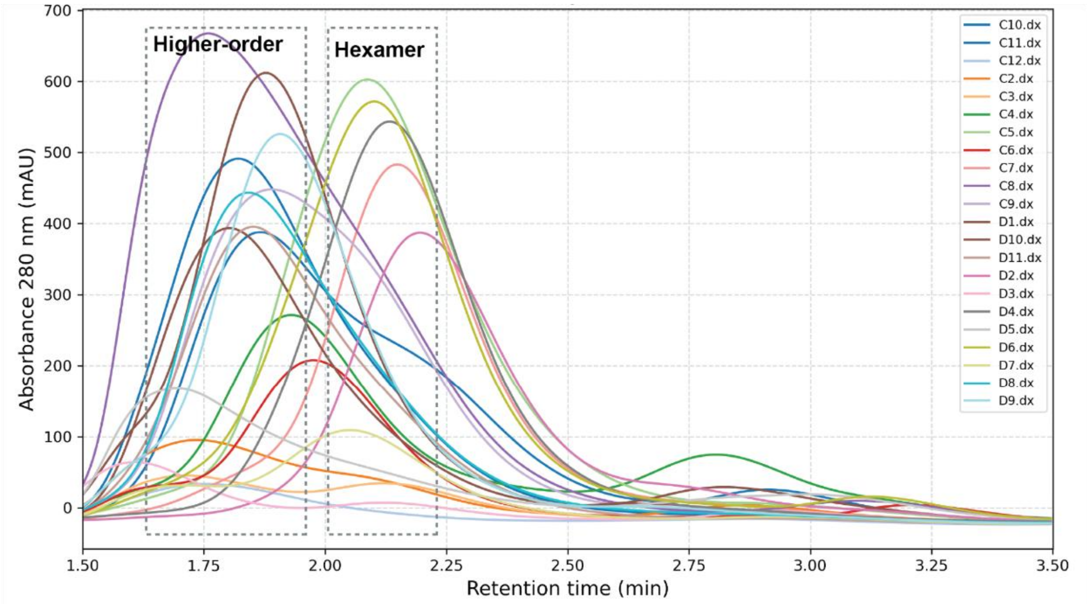

**Fig. S19 SEC traces of HAp_D3 oligomers packed against their N-termini using the Superdex 200 Increase 5/150 GL column.** The design HAp1_$D_3$C that's presented in the main text was named as C9 in this plot. Among these 22 designs, 11 formed higher-order species, five formed majorly hexamer, and six are ambiguous.

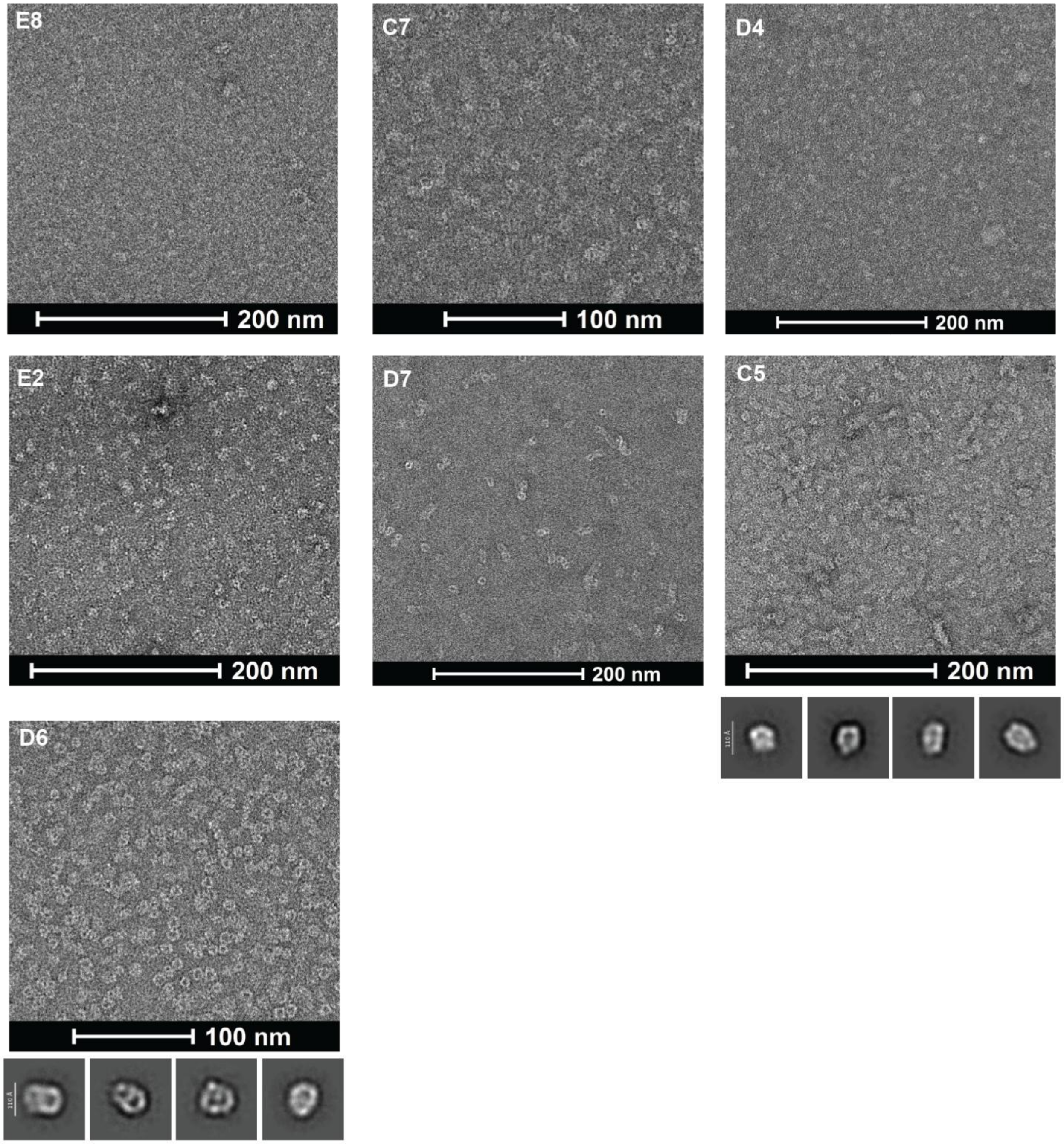

**Fig. S20 nsEM screening results of HAp1_D3 oligomers that did not fold into expected structures.** D7 sample showed short tube formation and D6 and C5 samples despite forming oligomers as screened by nsEM, 2D class averaging suggests that the protein structures do not match with the design model.

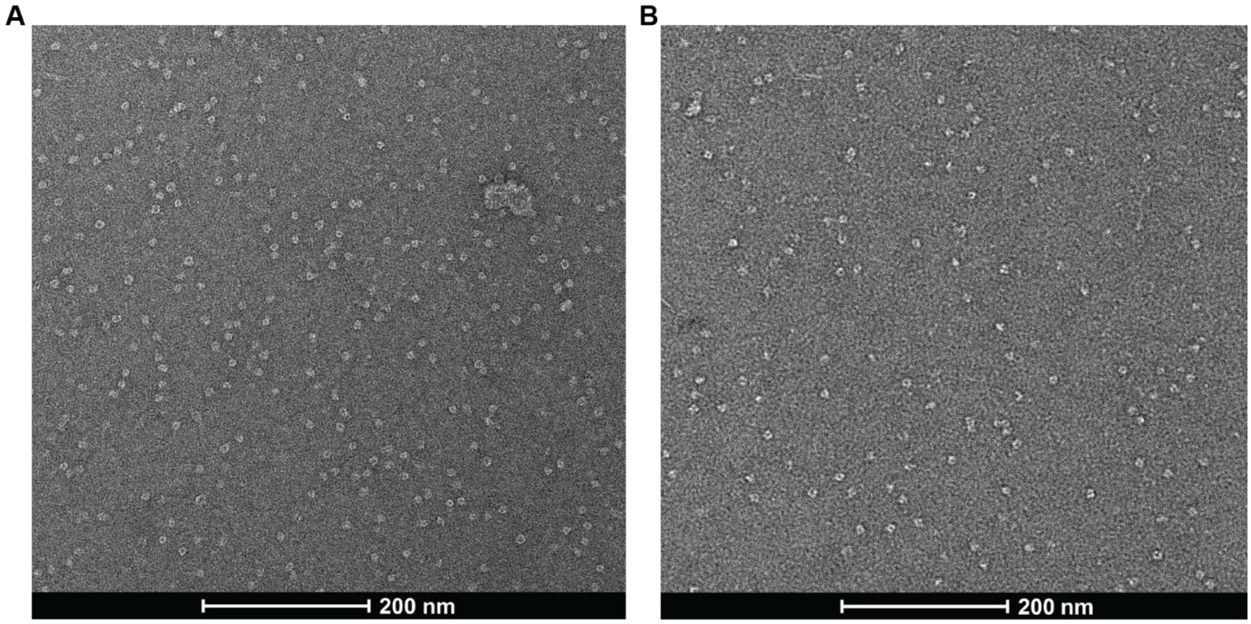

**Fig. S21 nsEM micrographs of $D_3$ oligomers.** HAp1_$D_3$N oligomer (A) and HAp1_$D_3$C oligomer (B).

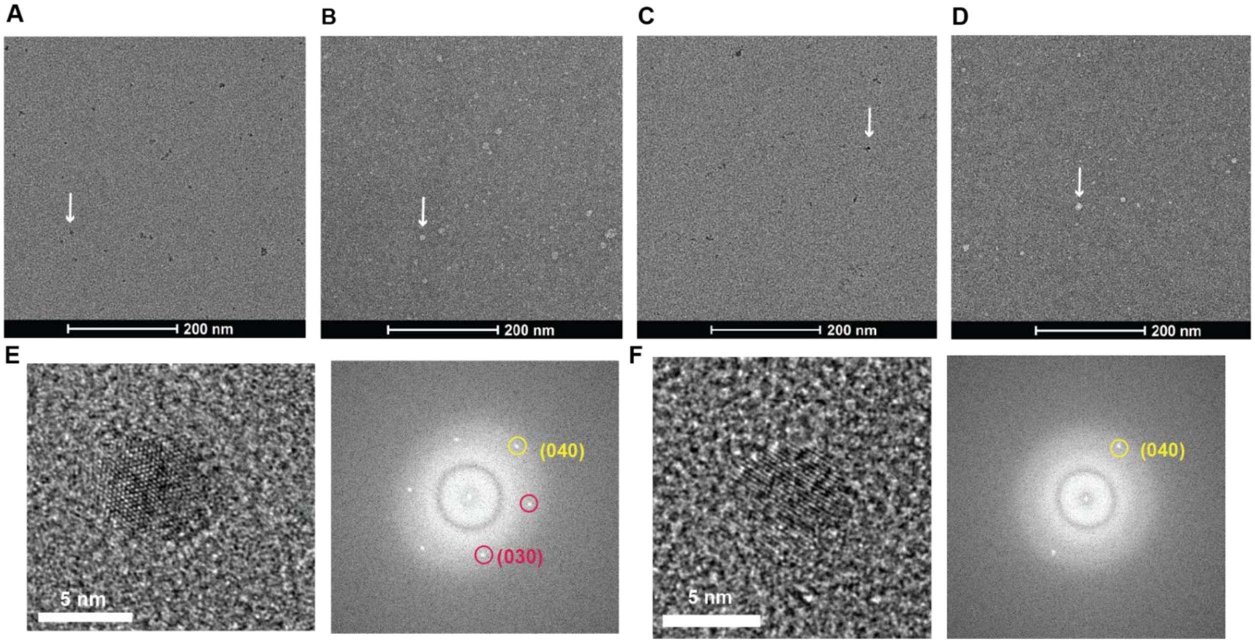

**Fig. S22 TEM results of $D_3$ oligomers mineralization (assay 1).** (**A**) and (**B**) are unstained TEM and uranyl formate stained nsEM of mineralized HAp1_$D_3$N oligomers, respectively. (**C**) and (**D**) are unstained TEM and uranyl formate stained nsEM of mineralized HAp1_$D_3$C oligomer, respectively. (**E**) HRTEM of mineralized HAp1_$D_3$N. (**F**) HRTEM of mineralized HAp1_$D_3$C oligomers.

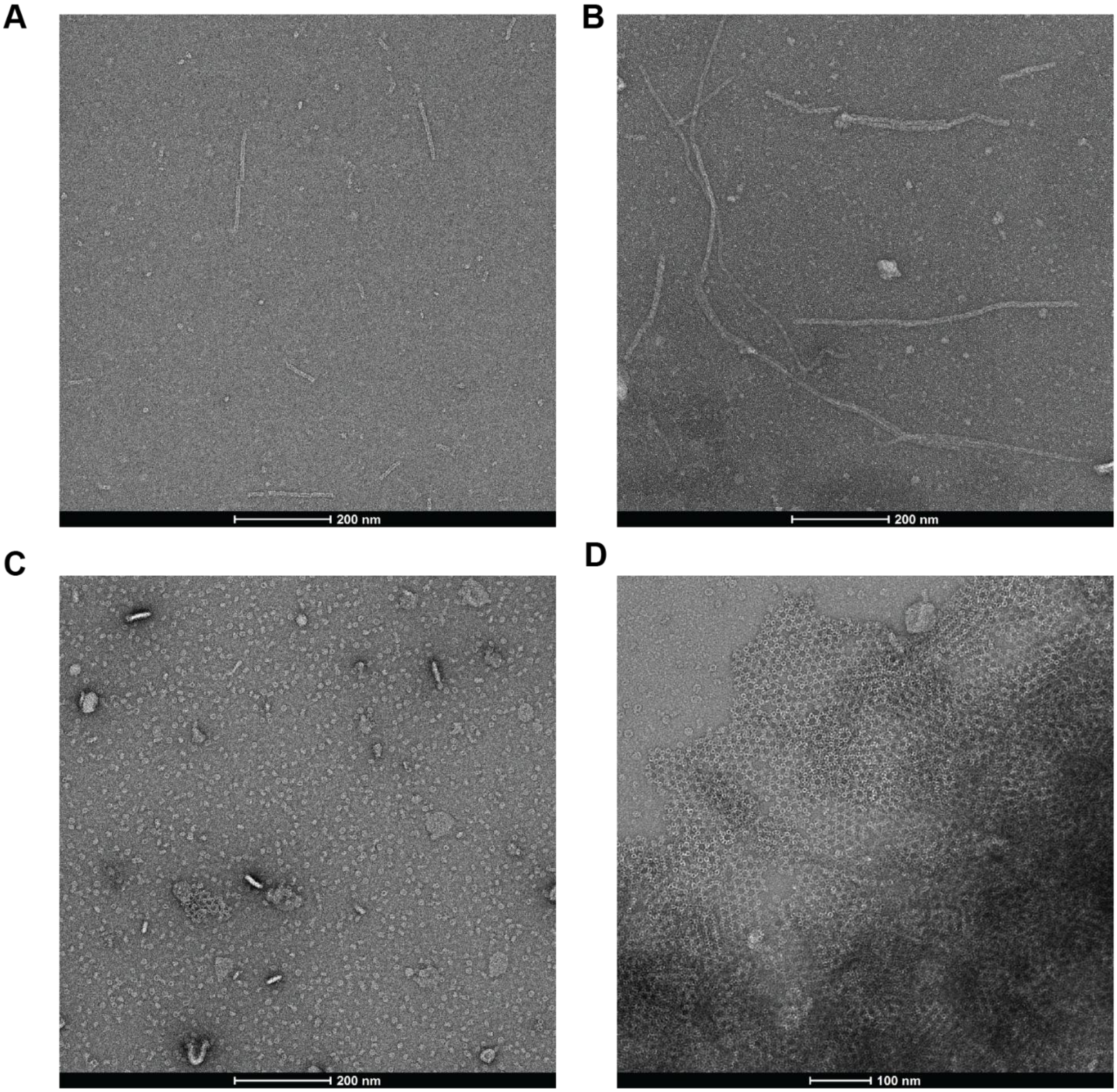

**Fig. S23. Concentration- and time-dependent assembly of HAp1_Nt and HAp1_Ar monitored by nsEM.** (A) HAp1_Nt assembled at 0.036 mg/mL. (B) HAp1_Nt assembled at 0.13 mg/mL for 1 hour. (C) HAp1_Ar immediately after IMAC purification at 0.12 mg/mL. (D) HAp1_Ar after 3 days of incubation at 0.12 mg/mL and 4°C.

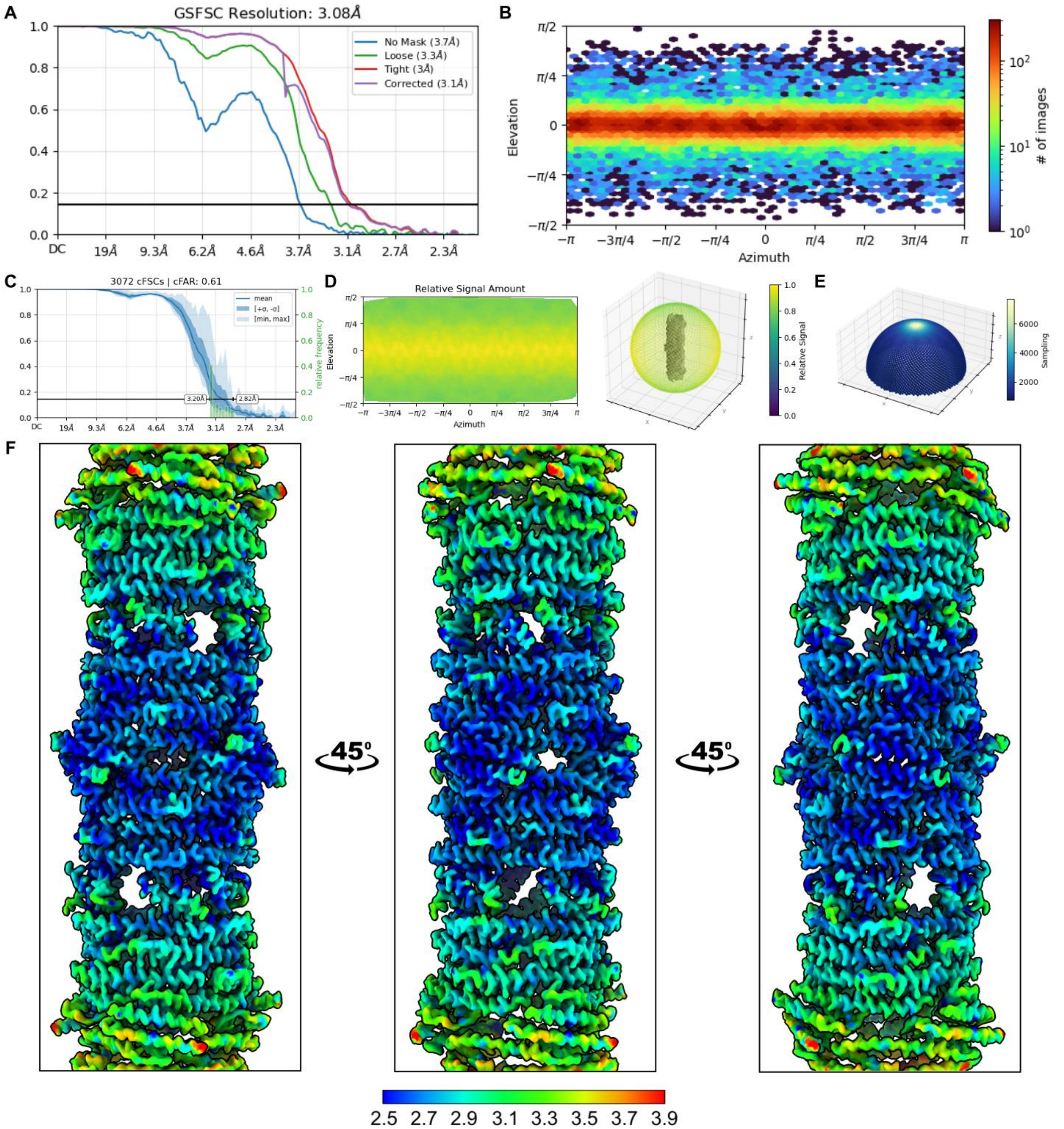

**Fig. S24 CryoEM analysis of de novo designed nanotube HAp1_Nt.**
(**A**) Global gold-standard Fourier shell correlation (GSFSC) curve indicating an estimated global resolution of 3.10 Å. (**B**) Orientational distribution plot demonstrating near-full angular sampling. (**C**) Summary of 3,072 conical FSCs. The conical FSC Area Ratio (cFAR) quantifies orientation bias as the ratio of the minimum to maximum weighted area-under-curve (wAuC) values across viewing directions. Each cFSC corresponds to a cone of 20° half-angle sampled uniformly across the sphere (Fibonacci scheme). cFAR values range from 0 to 1, with higher values indicating more uniform correlation landscapes and reduced orientation bias. (**D**) Relative signal amount as a function of viewing direction. (**E**) Fourier sampling plot (radius = 23, bins = 3,324; 0.0% empty; SCF = 0.845). The Sampling Compensation Factor (SCF) measures the

impact of the orientation distribution on spectral signal-to-noise ratio. SCF ranges from 0 to 1, with higher values reflecting more complete sampling. A value of 0.81 corresponds to a uniform side-view distribution; values above this threshold indicate adequate sampling. (**F**) Local resolution map colored by resolution and shown along the three rotational view angles.

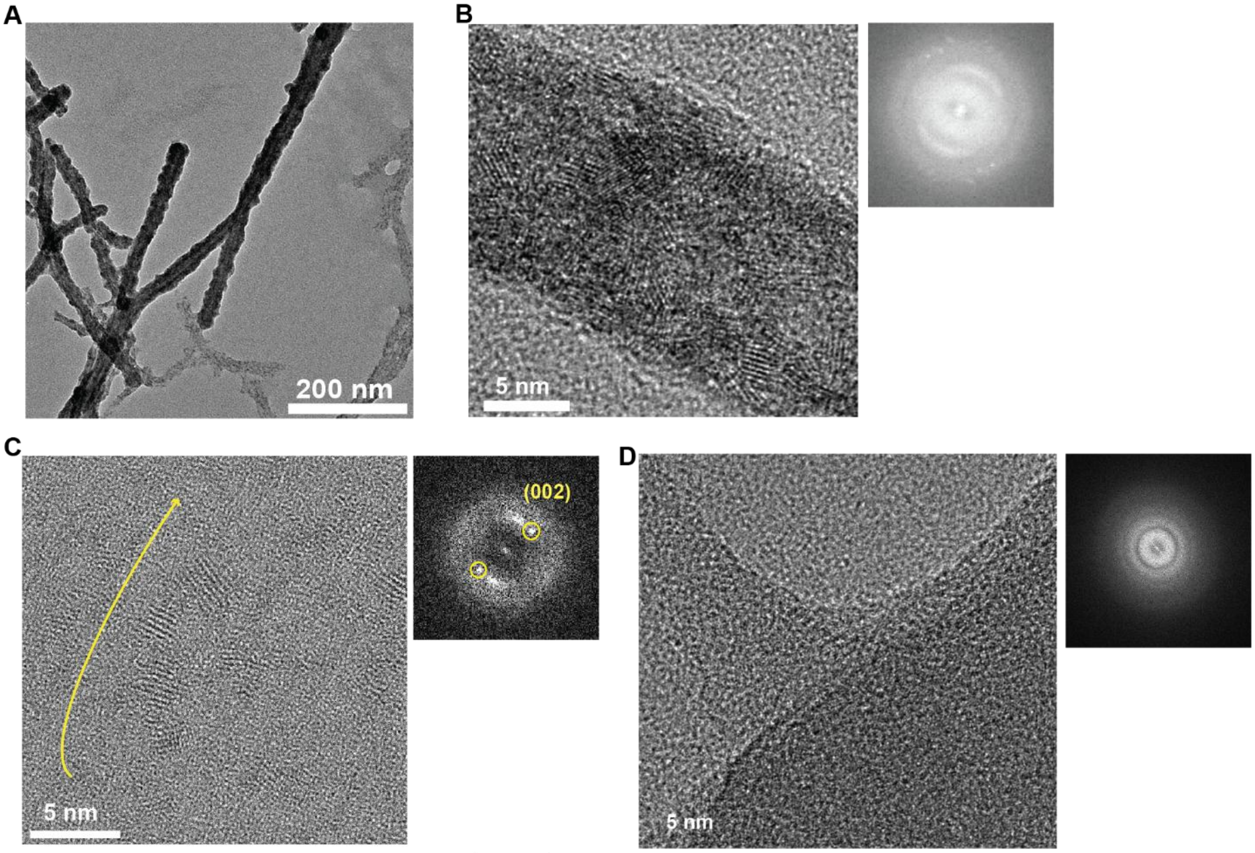

**Fig. S25. Mineralization of HAp1_Nt. (A)** TEM of mineralized HAp1_Nt with no pAsp (*assay 1*)**.** (**B**) HRTEM of mineralized HAp1_Nt (*assay 1*). (**C**) HRTEM of mineralized HAp1_Nt with pAsp (*assay 2*). (**D**) no protein control sample containing 2.5 mM $CaCl_2$, 1.0 mM $K_2HPO_4$ and 180 µg/mL pAsp (27 kDa) after 24 hours reaction. Most of the area on this grid has no features and the area that has features are noncrystalline. The right panel is FFT.

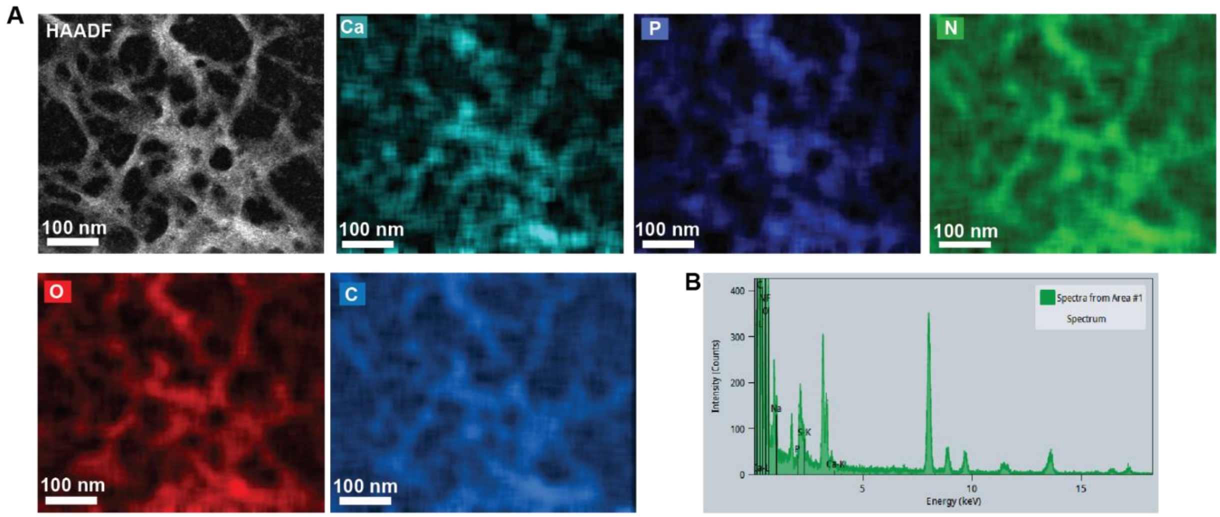

**Fig. S26 STEM-EDX characterization of HAp1_Nt.** (**A**) STEM-EDX mapping of HAp1_Nt at low magnification showing elements of calcium, phosphorus, nitrogen, oxygen and carbon overlap on filamentous features. (B) Elemental intensity of the mapping shown in A.

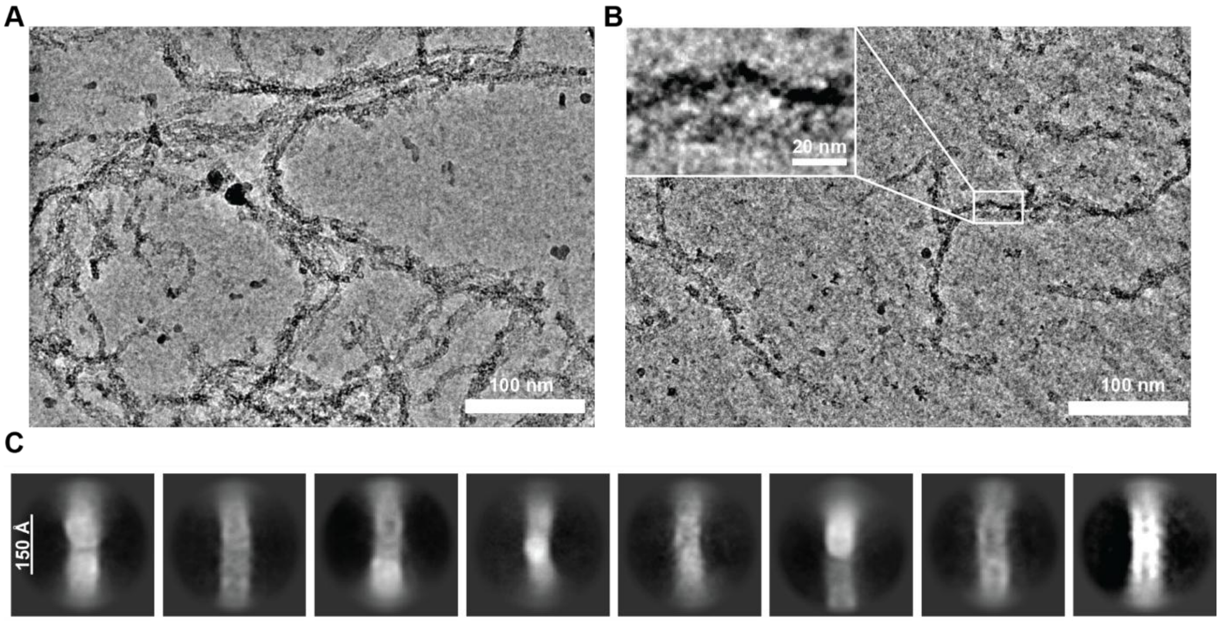

**Fig. S27 CryoEM characterization of mineralized HAp1_Nt with the presence of pAsp (assay2).** (**A**) and (**B**) Representative cryoEM micrographs of mineralized HAp1_Nt. (C) 2D class averaging of mineralized HAp1_Nt.

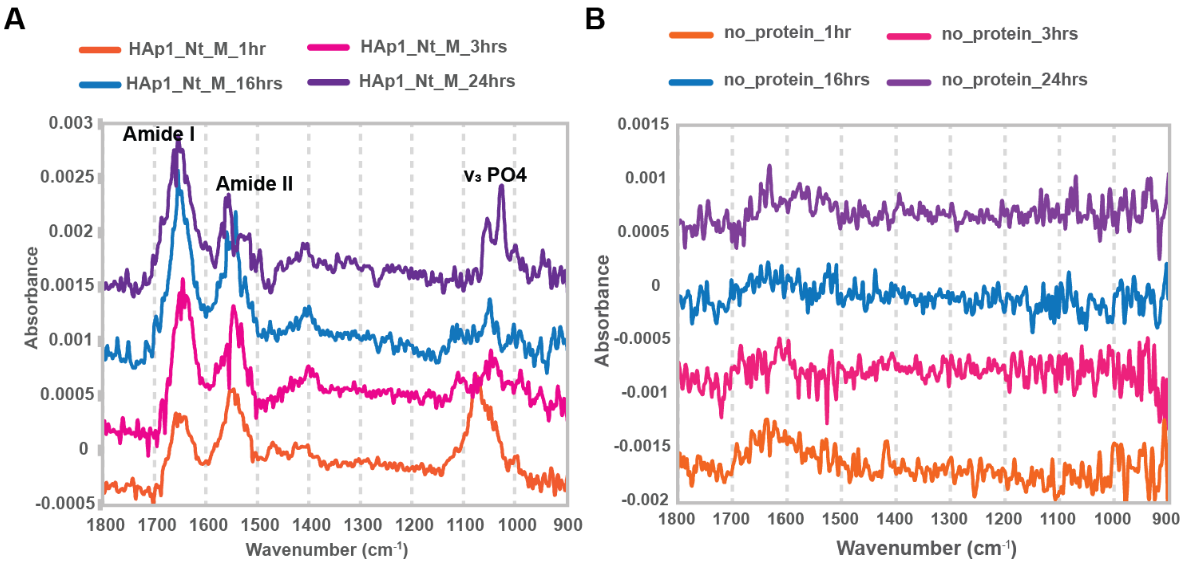

**Fig. S28. ATR-FTIR time-course study of mineralized HAp1_Nt.**
(**A**) This is the full ATR-FTIR spectrum of Fig. 4G. HAp1_Nt mineralized in 2.5 mM $CaCl_2$, 1.0 mM $K_2HPO_4$ and 180 µg/mL pAsp (27 kDa). (**B**) Reactions containing no protein but 2.5 mM $CaCl_2$, 1.0 mM $K_2HPO_4$ and 180 µg/mL pAsp (27 kDa).

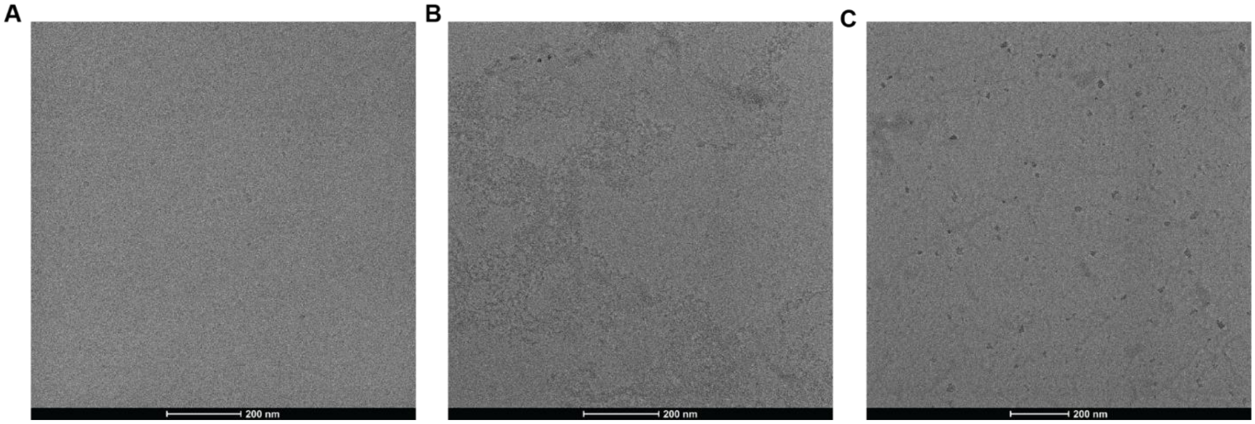

**Fig. S29 TEM of negative controls mineralization results.**
(**A**) precursors containing 2.5 mM $Ca^{2+}$, 1.0 mM $K_2HPO_4$, after 24 hours incubation at 37 ºC; (**B**) precursors containing 2.5 mM $Ca^{2+}$, 1.0 mM $K_2HPO_4$, 180 µg/mL pAsp (27 kDa), after 24 hours incubation at 37 ºC. (**C**) 3 µM HAp1_Nt and 2.5 mM $CaCl_2$ after 24 hours incubation at 37 ºC. Grids were washed 3 times with MQ $H_2O$ before drying and imaging.

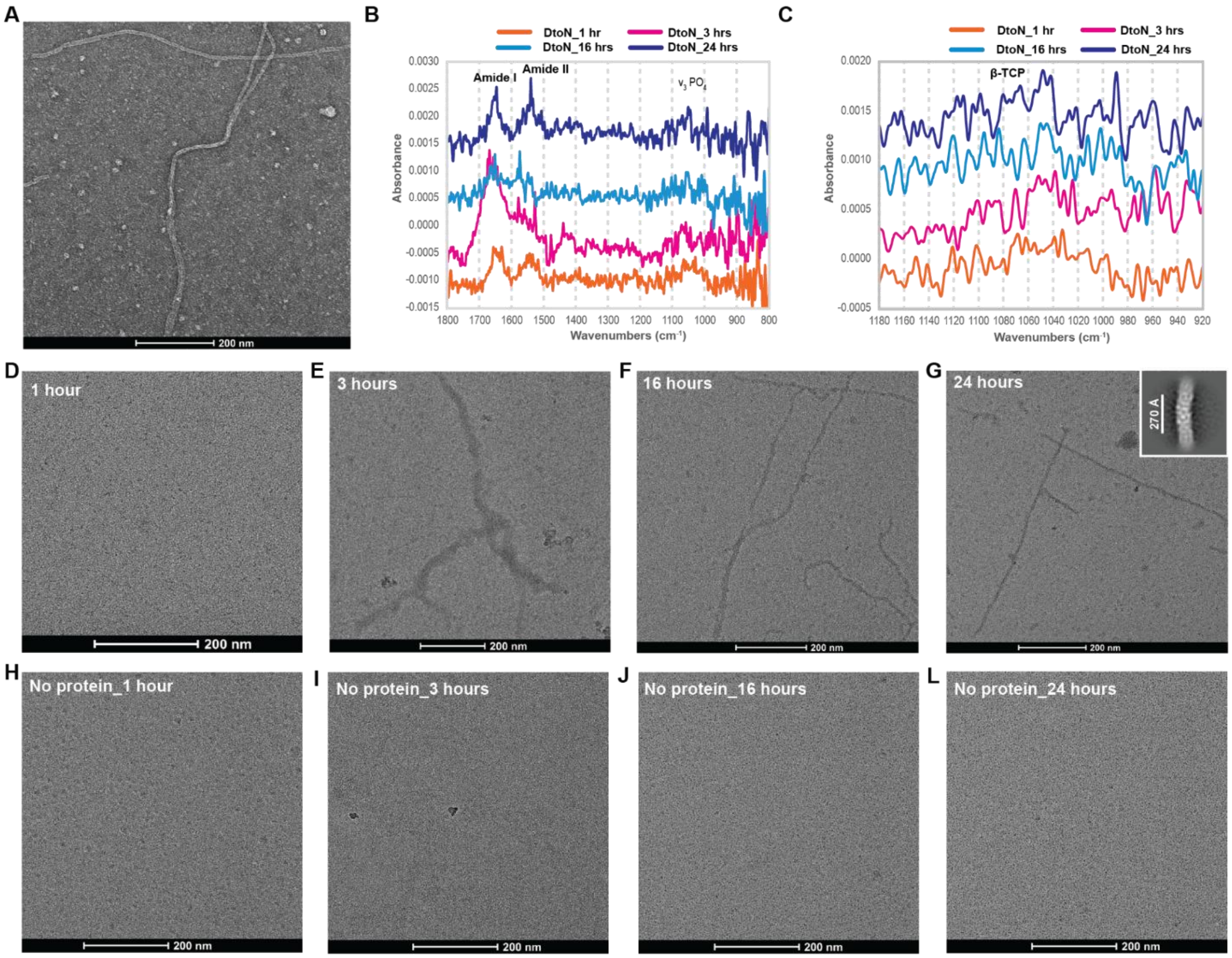

**Fig. S30 Characterization of HAp1_Nt_DtoN.**
(**A**) nsEM of HAp1_Nt_DtoN. (**B**) and (**C**) ATR-FTIR of mineralized HAp1_Nt_DtoN samples with 2.5 mM $CaCl_2$, 1.0 mM $K_2HPO_4$ and 180 µg/mL pAsp (27 kDa). (**D-G**) TEM of HAp1_Nt_DtoN mineralized with 2.5 mM $CaCl_2$, 1.0 mM $K_2HPO_4$ and 180 µg/mL pAsp (27 kDa). The inset in (**G**) is the 2D class averaging of the mineralized HAp1_Nt_DtoN. (**H-L**) TEM of the reactions with no proteins but 2.5 mM $CaCl_2$, 1.0 mM $K_2HPO_4$ and 180 µg/mL pAsp (27 kDa). TEM grids were washed three times with MQ $H_2O$ to remove salts and excessive precursors. Most of the ACP dissolved during this washing process unless stabilized by proteins attached to the grid.

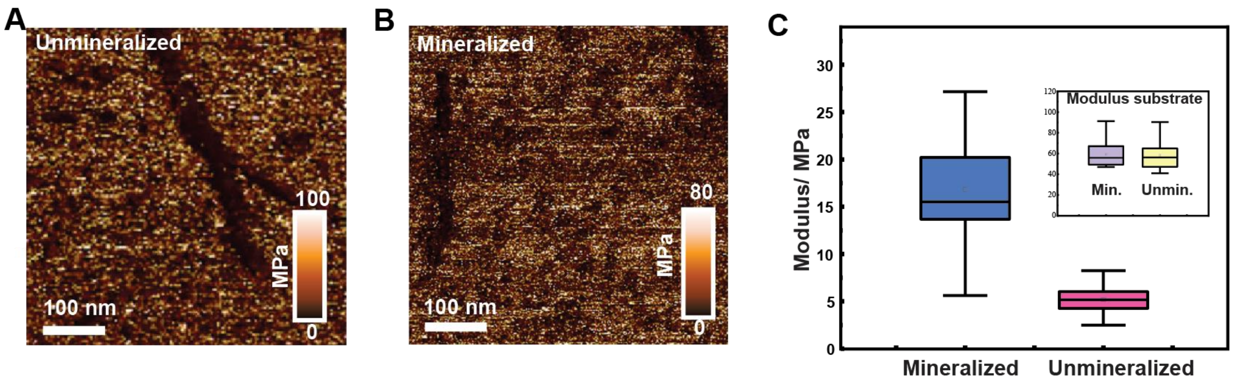

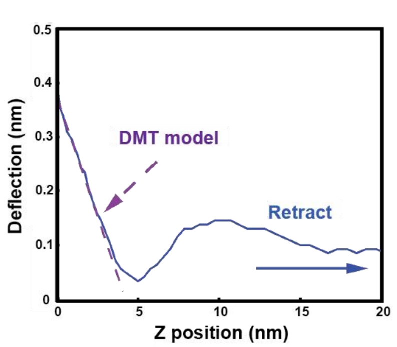

E

| Sample / Modulus | Unmin. | Min. |
|---|---|---|
| Mean (MPa) | 5.1 | 16.8 |
| Std (MPa) | 1.9 | 5.6 |
| Max (MPa) | 8.3 | 27.1 |
| Min (MPa) | 2.5 | 5.6 |

**Fig. S31 LP-AFM of nanotube HAp1_Nt.** (**A**) and (**B**) are mechanical maps of the unmineralized and mineralized nanotubes, respectively. (**C**) Modulus of the mineralized and unmineralized nanotubes, calculated based on Panel A and B. The inset is the modulus of the substrate standard. Purple is mineralized and yellow is unmineralized. (**D**) A representative force–distance curve collected on mineralized nanotubes. (**E**) Summary of LP-AFM mechanical force mapping results.

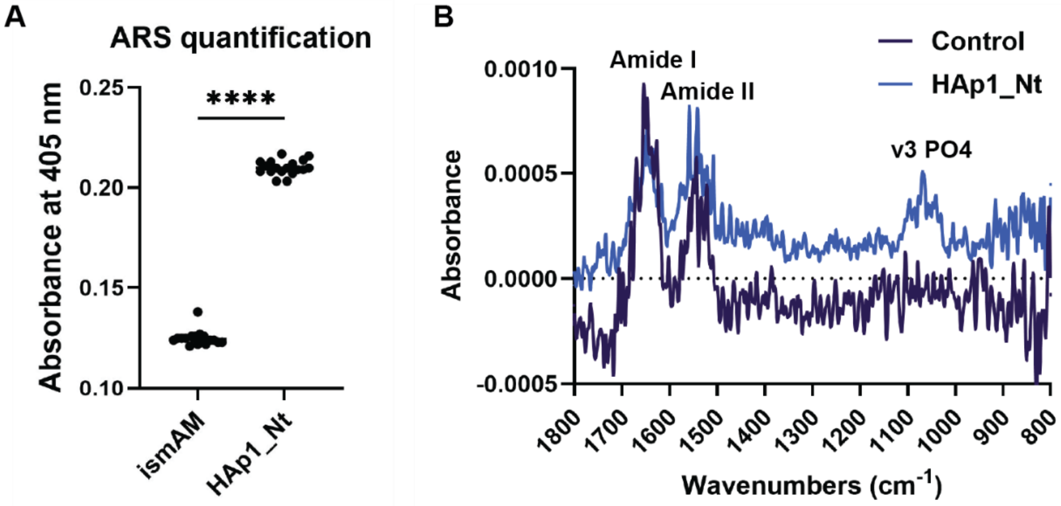

**Fig. S32 Biochemical characterization of mineralized human dental organoids.** (**A**) Alizarin Red S (ARS) quantification of organoids treated with HAp1_Nt or untreated controls ($p < 0.0001$). (**B**) ATR-FTIR spectra of organoids treated with HAp1_Nt or untreated controls.

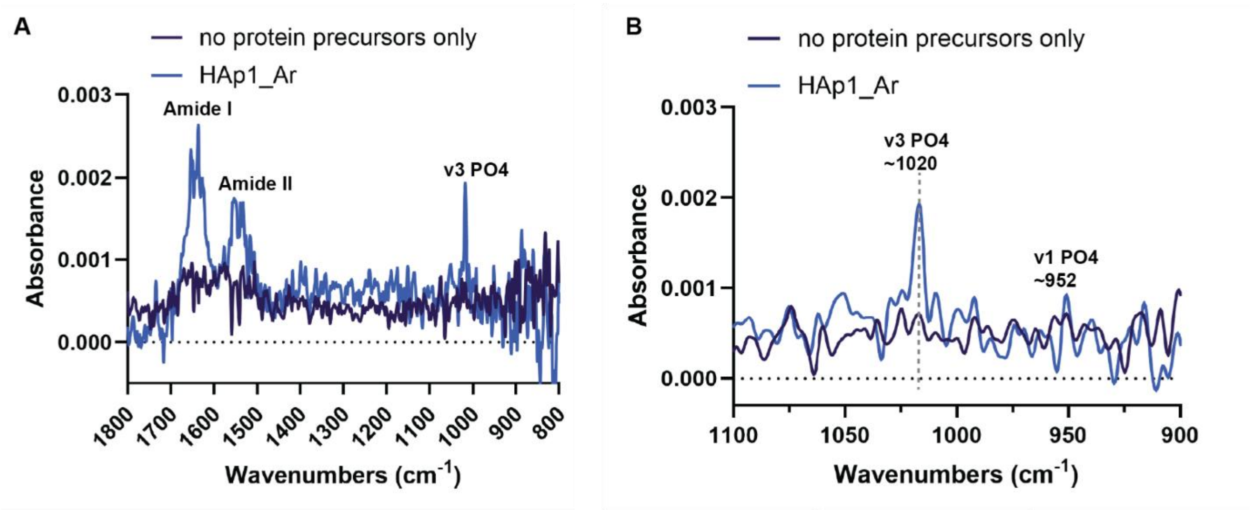

**Fig. S33 ATR-FTIR characterization of array HAp1_Ar mineralization (*assay 2*).** The no protein sample contains 2.5 mM $Ca^{2+}$, 1 mM $HPO4^{2-}$, and 180 µg/mL pAsp (27 kDa) after 24 hours reaction at 37 °C.

Table S1. Protein amino acid sequence and mass spectrometry information.

| Design name | Amino acid sequence | Expected [M+H]+ | Observed [M+H]+ |
|---|---|---|---|
| HAp1_C$_3$ | SGTLDEYEEELERYRKEAAALAAAGEDTSEANAKAVEAAKERYEAELKELEEKADELEARAPETLDACEEGLEVVYRKLAAYYGARGEDTSEVNARAQRLALLRYRHKYGELENDLALYEGLAPLTLDACEEGLEVCRKLAARAIAEGKDTSELVARAQRLGLLRFKHETAQLEHDAAVYEALAPLTLDACEEGAEVVYRKLIERAEALGEDTSEYRAKLEELQALRDRQEYGQLEFDAAVYAALAPLTKDAQEELAEVLRRLEKREKEGSGSHHWGSTHHHHHH | 31779.70 | 31779.87 |
| HAp1_C$_3$_DtoN | MTLNEYEEELERYRKEAAALAAAGEDTSEANAKAVEAAKERYEAELKELEEKADELEARAPETLNACEEGLEVVYRKLAAYYGARGEDTSEVNARAQRLALLRYRHKYGELENDLALYEGLAPLTLNACEEGLEVCRKLAARAIAEGKDTSELVARAQRLGLLRFKHETAQLEHDAAVYEALAPLTLNACEEGAEVVYRKLIERAEALGEDTSEYRAKLEELQALRDRQEYGQLEFDAAVYAALAPLTKDAQEELAEVLRRLEKREKELEHHHHHH | 31011.11 | 31011.55 |
| HAp1_C$_3$_EtoQ1 | MTLDEYQEELERYRKEAAALAAAGEDTSEANAKAVEAAKERYEAELKELEEKADELEARAPETLDACQEGLEVVYRKLAAYYGARGEDTSEVNARAQRLALLRYRHKYGELENDLALYEGLAPLTLDACQEGLEVCRKLAARAIAEGKDTSELVARAQRLGLLRFKHETAQLEHDAAVYEALAPLTLDACQEGAEVVYRKLIERAEALGEDTSEYRAKLEELQALRDRQEYGQLEFDAAVYAALAPLTKDAQEELAEVLRRLEKREKELEHHHHHH | 31011.11 | 31011.65 |
| HAp1_C$_3$_EtoQ2 | MTLDEYEQELERYRKEAAALAAAGEDTSEANAKAVEAAKERYEAELKELEEKADELEARAPETLDACEQGLEVVYRKLAAYYGARGEDTSEVNARAQRLALLRYRHKYGELENDLALYEGLAPLTLDACEQGLEVCRKLAARAIAEGKDTSELVARAQRLGLLRFKHETAQLEHDAAVYEALAPLTLDACEQGAEVVYRKLIERAEALGEDTSEYRAKLEELQALRDRQEYGQLEFDAAVYAALAPLTKDAQEELAEVLRRLEKREKELEHHHHHH | 31011.11 | 31011.66 |
| HAp1_C$_3$_EtoQ3 | MTLDEYEEELQRYRKEAAALAAAGEDTSEANAKAVEAAKERYEAELKELEEKADELEARAPETLDACEEGLQVYRKLAAYYGARGEDTSEVNARAQRLALLRYRHKYGELENDLALYEGLAPLTLDACEEGLQVCRKLAARAIAEGKDTSELVARAQRLGLLRFKHETAQLEHDAAVYEALAPQTLDACEEGAQVYRKLIERAEALGEDTSEYRAKLEELQALRDRQEYGQLEFDAAVYAALAPLTKDAQEELAEVLRRLEKREKELEHHHHHH | 31027.07 | 31027.66 |
| HAp1_C$_3$_DENQ | MTLNEYQQELQRYRKEAAALAAAGEDTSEANAKAVEAAKERYEAELKELEEKADELEARAPETLNACQQGLQVVYRKLAAYYGARGEDTSEVNARAQRLALLRYRHKYGELENDLALYEGLAPLTLNACQQGLQVCRKLAARAIAEGKDTSELVARAQRLGLLRFKHETAQLEHDAAVYEALAPQTLNACQQGAQVVYRKLIERAEALGEDTSEYRAKLEELQALRDRQEYGQLEFDAAVYAALAPLTKDAQEELAEVLRRLEKREKELEHHHHHH | 31015.31 | 31015.58 |
| HAp1_D$_3$N | SGSEAELAAAFKERAEALKEVNKAEARIWALEEEHGLTREEYRERLARYREDAALLRAAGVDTSETIANAIRAARLLHEAELAEWERDAARLDLRAPETLDACEEGLEVVYRKLAAYYGARGEDTSEVNARAQRLALLRYRHKYGELENDLALYEGLAPLTLDACEEGLEVCRKLAARAIAEGKDTSELVARAQRLGLLRFKHETAQLEHDAAVYEALAPLTLDACEEGAEVVYRKLIERAEALGEDTSEYRAKLEELQALRDRQEYGQLEFDAAVYAALAPLTKDAQEELAEVLRRLEKREKEGSHHHHHH | 34999.53 | 35042.59 [M+K]$^+$ or [M+Acetyl+H]$^+$ |
| HAp1_D$_3$C | SGTLDEYEEELERYRKEAAALAAAGEDTSEANAKAVEAAKERYEAELKELEEKADELEARAPETLDACEEGLEVVYRKLAAYYGARGEDTSEVNARAQRLALLRYRHKYGELENDLALYEGLAPLTLDACEEGLEVCRKLAARAIAEGKDTSELVARAQRLGLLRFKHETAQLEHDAAVYEALAPLTLDACEEGAEVVYRKLIERAAARGEDPKELAKKYEKLKEIGERLRWAQLERDAAIYAALAPLTVEARERLRAIIGELAKREVKEELGVSEEVAEVLGRVLAERRWTAVATELEGSHHHHHH | 34158.80 | 34158.81 |
| HAp1_Nt | SGSEQELAQAFKERAEALKEVNKAEARIWALEEEHGLTREEYRERLARYREDAALLRAAGVDTSETIQNAIRAARLLHEAELQEWERDAARLDLRAPETLDACEEGLEVVYRKLAAYYGARGEDTSEVNQRAQRLALLRYRHKYGELENDLALYEGLAPLTLDACEEGLEVCRKLAARAIAEGKDTSELVQRAQRLGLLRFKHETAQLEHDAAVYEALAPLTLDACEEGAEVVYRKLIERA | 38684.98 | 38684.56 |

| | | | |
|---|---|---|---|
| | **AARGEDPKELAKKYEKLKEIGERLRWAQLERDAAIYQALAPLTVEARERLRAIIGELAKREVKEELGVSEEVAEVLGRVLAERRWTQVATELE**GSHHHHHH | | |
| HAp1_Nt_DtoN | SG**SEQELAQAFKERAEALKEVNKAEARIWALEEEHGLTREEYRERLARYREDAALLRAAGVDTSETIQNAIRAARLLHEAELQEWERDAARLDLRAPETLNACEEGLEVYRKLAAYYGARGEDTSEVNQRAQRLALLRYRHKYGELENDLALYEGLAPLTLNACEEGLEVCRKLAARAIAEGKDTSELVQRAQRLGLLRFKHETAQLEHDAAVYEALAPLTLNACEEGAEVYRKLIERAAARGEDPKELAKKYEKLKEIGERLRWAQLERDAAIYQALAPLTVEARERLRAIIGELAKREVKEELGVSEEVAEVLGRVLAERRWTQVATELE**GSHHHHHH* | 38682.04 | 38681.48 |
| HAp1_Ar | SG**ARLALDATRYLREHITLPADVRAALRAAARRDRAEALKERNLAEAAIFLLEEEDGPTREEYRERLARYREDAALLRAAGVDTSETIANAIRAARLLHEAELAEWERDAARLDLRAPETLDACEEGLEVYRKLAAYYGARGEDTSEVNARAQRLALLRYRHKYGELENDLALYEGLAPLTLDACEEGLEVCRKLAARAIAEGKDTSELVARAQRLGLLRFKHETAQLEHDAAVYEALAPLTLDACEEGAEVYRKLIERAEALGEDTSEYRAKLEELQALRDRQEYGQLEFDAAVYAALAPLTKDAQEELAEVLRRLEKREKE**GSHHHHHH | 37359.22 | 37358.51 |

Sequences in bold are designed sequences. Depending on the vectors that the designs were cloned into, the expressed protein sequences can contain methionine as a starting residue, "GS" linker, "LE" linker, SNAC tag, or histidine tag near either C or N termini. Yellow highlighted residues are the designed calcium-coordinating residues. These follow the 'DXXEEXXE' motif (X = any amino acid), with the first entry corresponding to the structure shown in Fig. 2E. Such residues were mutated to noncharged residues during the mutation studies (highlighted in green). The linkers or tags do not interfere with protein assembly or functionalization. After combining the sequence of HAp1_$D_3$N and HAp1_$D_3$C for nanotube design, a few alanine residues on the surface of the nanotube were rationally redesigned into glutamine residues to improve solubility and reduce aggregation.

Table S2. Protein amino acid sequence information of $C_3$ oligomer designs that folded into oligomeric structure but did not exhibit confined HAp mineralization.

| | |
|---|---|
| HAp1_$C_3$_2 | KEKDEKEEEDDRAAKEVIDELRAEARLLFAEGKSIEEIGEEVEKLYEEIK<br>PSLSDRETRKVERAIPDIMEEIEDIAAKEVIDRIRAEAILLYAAGKSIEEIKE<br>EVEKIAKEIEPSLTEKERKKVRRAIPDIIEEIEDRAAKKVIDELRKTAEEMF<br>KEGKSIEEIQQEVVALAREVFPSLTPYERRKVARAILDIIEEIEDKAAKKV<br>IDELKERALELFEEGKSIEEIQKEVVKLAKEKMPSLSAYEQRKVARAILDI<br>IEEIQDRKDKEEEDEERAAEKEAEE |
| HAp1_$C_3$_3 | GPADALLAELEALRAEVERAEDVADEWGTRLILAEGSGSAEVRAEVEA<br>ELEARLAALPPWQALLARLRLREFRVRRAEDVADEFGTLLILARGSGDP<br>EVQAEVEALIDRLLAALPPWQRLLAELRLRRYEVEYAEDVADEFGELRI<br>LAAGSGDAAVRAEVDALLQELLDALPPWQRLLAERRLERYEVRYAED<br>VADEFGELLLLAEESGDPEVRAEVEARLAALLAALTEDLAALARARLA<br>EYKERRKEDEEDEFGLLEELRAEGS |
| HAp1_$C_3$_4 | SAVEEVRARLEERRAAEDAAEEAGRELVLAELAGDEELLARLRAEIARQ<br>DAVFQVRVRRAIRRAWEDLAEELGRELVDAELAGDEERLAALRAEIEK<br>LPPVLQVRVERAIRRAWEDLEEELGEEIVAAELAGDEELLARLRAEVER<br>LPAVLRVGVEEAIERAKEDLAEEERRRLERARRAGDEELVAALEARLEG<br>S |
| HAp1_$C_3$_5 | GPADALLAEFEALRAQVEGAEDLADERGTRLILAEGSGDAAVRAEVEA<br>ELEAALAALPPWQALLAKFRLRRFRVRFAEDLADEFGTLLILARGSGDA<br>AVQAEVEALIQRLLDFLPPWLRLLAEFRLRRHEVEFAEDLADEFGELRIL<br>AAGSGDAEVRAEVDALIQELLDALPPWQRLLAERRLERHEVRFAEDLA<br>DEFGELALLAEASGDAAVRAEVDAALDALEAALTEDLAALFRARLAEY<br>RVRRKEDEEDEFGLLKKLRAEGS |
| HAp1_$C_3$_6 | TREEAEDKYEEARKRLAEAGELDPELVAEAAKYEIERQEKKLEELEEKI<br>EELEEKLKEGATREECEDGYEACRKVIAYKGKIDDDLVEKASRFEIARQ<br>EQKRVELEFKLAVYAALLEHGMTREQAEDGAEAARRLLAYAGEVRPD<br>LVAALSRFEIARQEQKLVELEYKVRVLQALLEHGMTREQAEDGAEACR<br>KILEYKGEEDPEILALLERFEEARRRQKLVELQVKVAVYSALEKYGMTK<br>EEKEDRDEAIRRIKEYGS |

**Table S3. CryoEM Data Collection, Processing, and Modeling Statistics**

| | **HAp1_Nt PDB: 9ZNK** |
|---|---|
| **Data Collection** | |
| Microscope | Titan Krios |
| Voltage (kV) | 300 |
| Detector | Gatan K3 |
| Energy Filter | Gatan BioQuantum Gif |
| Recording mode | SuperResolution |
| Magnification | 105,000 X |
| Movie micrograph pixel size (Å) | 0.5305 |
| Dose rate ($e^{-}/Å^2/s$) | 13.2 |
| No. of frames per movie micrograph | 75 |
| Frame exposure time (s) | 0.0505 |
| Movie micrograph exposure time (s) | 5.009 |
| Total dose ($e^{-}/Å^2$) | 50.0 |
| Under focus range (µm) | 0.8 - 1.8 |
| Number of movie micrographs | 4,018 |
| Symmetry Applied | D3 |
| Extraction Box Size (pix) | 700→350 |
| Initial particle images (no.) | 358,483 |
| Final particle images (no.) | 69,261 |
| Map resolution (Å) | 3.08 |
| FCS threshold | 0.143 |
| Map resolution range (Å) | 2.5-3.3 |
| Initial model used | Design Model |
| Map resolution (Å) | 3.08 |
| FCS threshold | 0.143 |
| Model resolution range (Å) | 2.5-3.3 |
| Map sharpening B factor | -87.6 |
| Model composition | |
| Non-hydrogen atoms | 31,872 |

| | |
|---|---|
| Protein Residues | 3,984 |
| Ligands | Na |
| *B* factors (Å) | |
| Protein | 22.23 |
| Ligands | Na |
| R.M.S. deviations | |
| Bond lengths (Å) | 0.003 |
| Bond angles (º) | 0.424 |
| Validation | |
| MolProbity score | 0.98 |
| Clashscore | 1.59 |
| Rotamer Outliers (%) | 1.38 |
| Ramachandran plot | |
| Favored (%) | 98.41 |
| Allowed (%) | 1.59 |
| Disallowed (%) | 0 |
